# Mechanistic insights into the functioning of GMP synthetase: a two-subunit, allosterically regulated, ammonia tunnelling enzyme

**DOI:** 10.1101/2022.02.27.481963

**Authors:** Santosh Shivakumaraswamy, Sanjeev Kumar, Asutosh Bellur, Satya Dev Polisetty, Hemalatha Balaram

## Abstract

Guanosine 5’-monophosphate (GMP) synthetases, enzymes that catalyze the conversion of xanthosine 5’-monophosphate (XMP) to GMP are comprised of two different catalytic units, which are either two domains of a polypeptide chain or two subunits that associate to form a complex. The glutamine amidotransferase (GATase) unit hydrolyzes glutamine generating ammonia and the ATP pyrophosphatase (ATPPase) unit catalyzes the formation of AMP-XMP intermediate. The substrate-bound ATPPase allosterically activates GATase and the ammonia thus generated is tunnelled to the ATPPase active site where it reacts with AMP-XMP generating GMP. In ammonia tunnelling enzymes reported thus far, a tight complex of the two subunits is observed, while the interaction of the two subunits of *Methanocaldococcus jannaschii* GMP synthetase (MjGMPS) is transient with the underlying mechanism of allostery and substrate channelling largely unclear. Here, we present a mechanistic model encompassing the various steps in the catalytic cycle of MjGMPS based on biochemical experiments, crystal structure and cross-linking mass spectrometry guided integrative modelling. pH dependence of enzyme kinetics establish that ammonia is tunnelled across the subunits with the lifetime of the complex being ≤ 0.5 s. The crystal structure of XMP-bound ATPPase subunit reported herein highlights the role of conformationally dynamic loops in enabling catalysis. The structure of MjGMPS derived using restraints obtained from cross-linking mass spectrometry has enabled the visualization of subunit interactions that enable allostery under catalytic conditions. We integrate the results and propose a functional mechanism for MjGMPS detailing the various steps involved in catalysis.

**Graphical Abstract:** 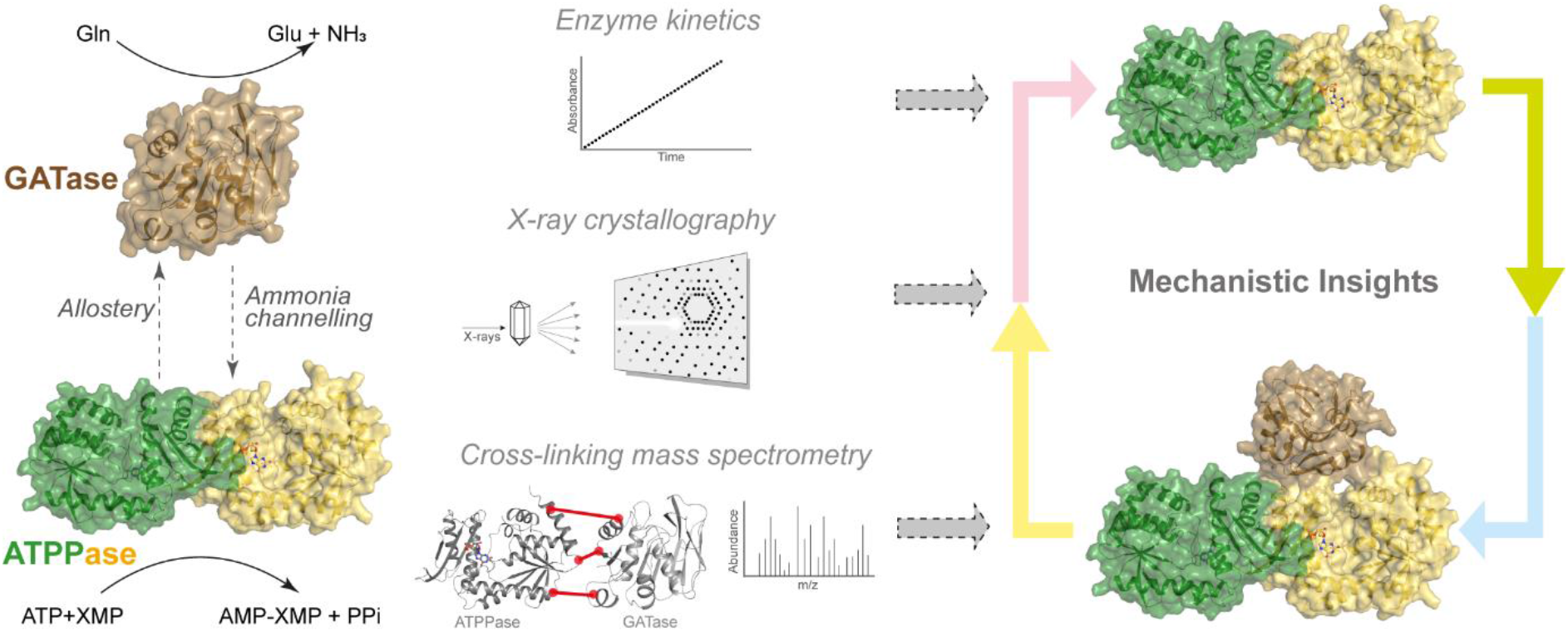

## Introduction

Cascade reactions catalyzed by bifunctional enzymes and multienzyme complexes are nature’s equivalent of one-pot chemical synthesis.^1, 2^ In these reactions, the intermediates that are labile/toxic are sequestered from the cellular milieu by shuttling them from one active site to the other through a physically constrained tunnel, a process termed tunnelling or channelling.^3, 4^ Among the intermediates that are tunnelled, ammonia is by far the most common.^5–7^ Enzymes of the glutamine (Gln) amidotransferase family are ammonia tunnellers that generate ammonia by hydrolysing the amide group of Gln and utilize it to replace an oxo group on acceptor metabolites that can be amino acids, sugars, nucleotides, cofactors and antibiotics.^5, 7, 8^ The production of ammonia and its incorporation occur on separate catalytic units, which can be either two domains of a protein chain or two subunits that associate to form a complex. The Gln amidotransferase (GATase) unit catalyzes the hydrolysis of the side chain of Gln generating ammonia (Figure 1A), while the acceptor unit catalyzes the incorporation of nitrogen (ammonia) into the acceptor molecule.^6^ The GATase unit is inactive and the binding of the substrate(s) to the acceptor unit allosterically activates GATase resulting in very large increase in activity.^9^ The mechanism of ammonia tunnelling is diverse with preformed tunnels observed in some Gln amidotransferases whereas in others, the conduit is formed through substrate induced local conformational changes or reorganization of entire domains.^3, 10, 11^

**Figure 1:**
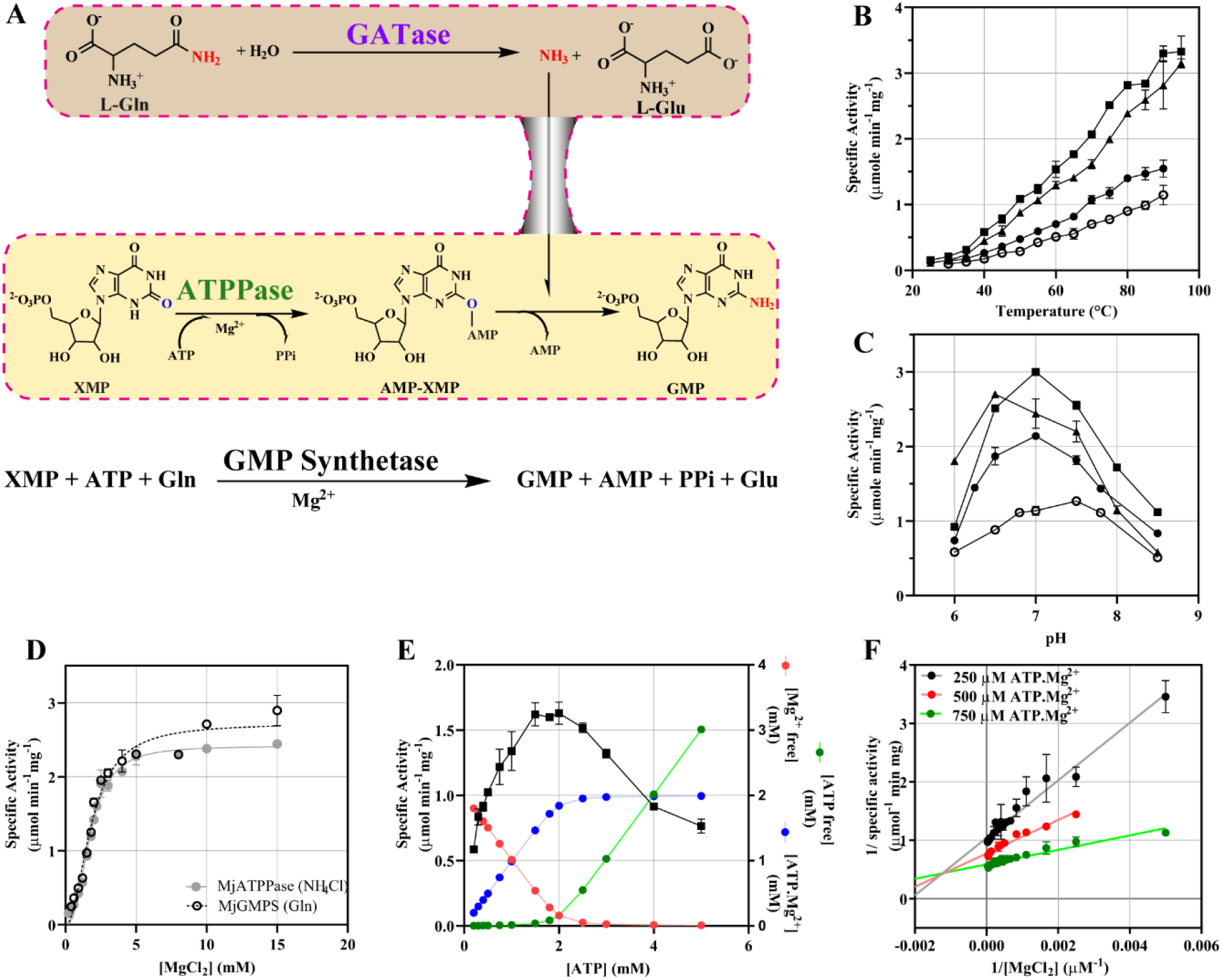
Reaction scheme and biochemical characterization of MjATPPase. (A) Reaction catalyzed by the GATase unit, ATPPase unit and GMPS. Effect of (B) temperature and (C) pH on NH_4_Cl dependent GMP formation by MjATPPase (▪) and fused MjGMPS (●), and Gln dependent GMP formation by MjGMPS (▴) and fused MjGMPS (○). (D) Effect of increasing [MgCl_2_] on the activity of MjATPPase and MjGMPS at fixed [ATP] of 2 mM. The data best fitted to positive cooperativity. (E) Dependence of MjATPPase activity on increasing [ATP] at fixed [MgCl_2_] of 2 mM. (F) Lineweaver-Burk plot of MjATPPase activity at different fixed [ATP.Mg^2+^] and varying [MgCl_2_]. Intersecting pattern suggests that ATP does not compete with ATP.Mg^2+^. The mean and standard deviation from two experiments is plotted in panels B to F.

The GATase unit forms the basis for classifying Gln amidotransferases into class I, where the fold resembles that of α/β hydrolases and class II, that possess the fold of N-terminal nucleophile hydrolases.^6^ GMP synthetase (GMPS) is a class I Gln amidotransferase that functions in *de novo* purine biosynthesis catalysing the conversion of xanthosine 5’-monophosphate (XMP) to guanosine 5’-monophosphate (GMP) (Figure 1A).^12^ The acceptor unit termed the ATP pyrophosphatase (ATPPase) binds a molecule of ATP.Mg^2+^ and XMP, and catalyzes the formation of AMP-XMP intermediate (Figure 1A).^13^ In GMP synthetases from bacteria and eukaryotes, the GATase and ATPPase units are two domains of a polypeptide chain (two-domain type) whereas the two units are independent proteins in many archaea (two-subunit type).^14^ *In vitro,* the two-domain type GMP synthetases as well as the ATPPase subunit of the two-subunit type enzymes can also utilize external ammonia to synthesize GMP.^14–18^ In both forms, the GATase domain/subunit is inactive or weakly active and the binding of ATP.Mg^2+^ and XMP to the ATPPase domain/subunit allosterically activates the GATase domain/subunit leading to Gln binding and hydrolysis. The ammonia thus generated is tunnelled to the ATPPase active site where it replaces the adenylate group in AMP-XMP through a nucleophilic attack, generating GMP (Figure 1A).^13, 15, 16, 19^ In the two-domain type GMPS from *Plasmodium falciparum* (PfGMPS), the products of the two domains are produced in stoichiometric amounts suggesting that the reactions on the two domains are synchronized^19^.

Structural and biochemical studies on two-domain type GMP synthetases, in particular, the studies on PfGMPS have considerably advanced our understanding of the functional mechanism. However, the two-subunit type GMP synthetases remain poorly characterized as only GATase activation and subunit association have been examined for the enzymes from *Methanocaldococcus jannaschii* (MjGMPS) and *Pyrococcus horikoshii* (PhGMPS).^14, 20^ In PhGMPS, the complex comprising the two subunits can be detected only in the presence of ATP.Mg^2+^ and XMP and the interaction leads to allosteric activation of GATase subunit and catalysis of GMP formation.^14^ In MjGMPS however, the interaction between the GATase (MjGATase) and ATPPase (MjATPPase) subunits is transient even in the presence of substrates.^20^

Among the nine class I Gln amidotransferases reported till date, except for cytidine triphosphate synthetase and 2-amino-2-desoxyisochorismate synthase where only two-domain type enzymes have been characterized, two-subunit type enzymes have been studied for the seven others. Considering that domain crosstalk underlies allostery and ammonia tunnelling, the GATase and the acceptor subunits form a tight complex in six Gln amidotransferases namely carbamoyl phosphate synthetase^21^, imidazole glycerol phosphate synthase^9^, formylglycinamide ribonucleotide amidotransferase^22^, pyridoxal 5’-phosphate synthase^23^, anthranilate synthase^24^ and 4-amino 4-deoxychorismate synthase^25^, excluding GMP synthetases. Hence, PhGMPS and MjGMPS are unique compared to all characterized two-subunit type Gln amidotransferases wherein the transient interaction between the subunits runs counter to the strong association of subunits enabling the activity of these bifunctional enzymes.

Herein, we investigate the mechanism underlying the functioning of MjGMPS using enzymatic assays, X-ray crystallography, cross-linking mass spectrometry (XL-MS) and integrative modelling. Using kinetic experiments, we estimate the lifetime of the transient complex and thereafter establish that ammonia is channelled across the subunits despite the small time scales of association. We then report the crystal structure of the ATPPase subunit which reveals novel insights into the catalytic mechanism and the obligate dimeric nature of the ATPPase domains/subunits. To understand the structural basis of subunit crosstalk, we trap the complex by covalent cross-linking, identify the cross-linked residues using mass spectrometry and use the crosslinks as distance restraints to reconstruct the structure of the complex. The study provides structural details of the interaction between MjGATase and MjATPPase subunits that enables allosteric activation of the GATase subunit. The results from the kinetic assays in conjunction with XL-MS guided model of MjGMPS structure has enabled us to map out the different steps of catalysis. This study also demonstrates the applicability of XL-MS to probe the structural details of transient complexes that are elusive to other methods.

## Results

### MjATPPase displays unusually high affinity for NH_4_Cl and harbours a Mg^2+^ binding site

Similar to other GMP synthetases^15, 17, 18^, and Gln amidotransferases^5^, MjATPPase subunit alone catalyzed the synthesis of GMP using NH_4_Cl as the nitrogen source. Consistent with the thermophilic nature of *M. jannaschii*, the specific activity increased with temperature, with the activity at 95 °C being 23-fold higher than at 25 °C (Figure 1B). The optimum pH for activity at 70 °C was 7 (Figure 1C), differing from GMP synthetases that prefer a pH of 8.5-9.2 for NH_4_Cl dependent GMP formation.^15, 19^ Kinetic studies performed at 70 °C and pH 7 revealed an absence of cooperativity in substrate binding. The *K*_m_ value for NH_4_Cl was 4.1 ± 0.2 mM (Table S1), the lowest reported for any GMP synthetase.^16–18, 26^ The *k*_cat_ for NH_4_Cl dependent GMP formation was 1.91 ± 0.02 s^-1^ and the *K*_m_ values for ATP.Mg^2+^ and XMP were 447 ± 5 µM and 30 ± 2 µM, respectively (Table S1).

GMP synthetases are known to possess an additional Mg^2+^ binding site apart from the ATP.Mg^2+^ site.^13, 16–18^ However, it remains unclear if this site is in the GATase or the ATPPase domains/subunits. We utilized the two-subunit arrangement of MjGMPS to resolve the location of the Mg^2+^ binding site. In kinetic experiments with the MjATPPase subunit, we noticed a sigmoidal increase in NH_4_Cl dependent GMP formation with increasing [Mg^2+^] (Figure 1D). Probing further, we measured MjATPPase activity at fixed [Mg^2+^] of 2 mM and increasing [ATP]. The activity dropped at [ATP] exceeding 2 mM, which could be due to the competition of ATP with ATP.Mg^2+^ binding site or a reduction in free [Mg^2+^] on account of chelation by ATP (Figure 1E). To distinguish between the two scenarios, the [Mg^2+^] was varied at various fixed [ATP.Mg^2+^] and saturating [XMP] (Figure 1F). The lines in the double reciprocal plot displayed an intersecting pattern suggesting that ATP does not compete with ATP.Mg^2+^. Hence, the reduction in activity upon increasing [ATP] is due to the chelation of free Mg^2+^ which is required by the MjATPPase subunit for maximal activity. The Hill coefficient and *K*_0.5,_ which were 2.45 ± 0.05 and 1.75 ± 0.02 mM, respectively did not change in the presence of the MjGATase subunit (Hill coefficient = 2.05 ± 0.07 and *K*_0.5_ = 1.84 ± 0.01 mM), corroborating the presence of an additional Mg^2+^ binding site in the ATPPase subunit (Figure 1D).

### Evidence for ammonia tunnelling in MjGMPS

Upon screening a combination of various ligands, we identified that apart from the substrate pair ATP.Mg^2+^+XMP, even AMP-PNP+XMP and XMP+pyrophosphate (PPi) also activated MjGATase, albeit to lower levels (Figure 2A). This suggests that formation of AMP-XMP intermediate is not a precondition for activation. It has been shown that Gln dependent GMP formation is maximum when the ratio of MjGATase and MjATPPase is 1:1.^20^ Upon measuring the temperature dependence of GMP formation by MjGMPS (equimolar concentrations of MjGATase+MjATPPase), we observed a trend similar to MjATPPase, as the activity increased linearly with temperature up to 95 °C and the optimum pH at 70 °C was 7.0 (Figure 1B, C). Kinetic studies performed at a temperature of 70 °C and pH of 7 indicated that all substrates exhibited Michaelis-Menten behaviour. The *K*_m_ values of 452 ± 3 µM for ATP.Mg^2+^ and 61 ± 3 µM for XMP were similar to and two-fold higher, respectively when compared to MjATPPase (Table S1). The *K*_m_ for Gln was 520 ± 8 µM, a value similar to other GMP synthetases. The *k*_cat_ for Gln dependent GMP formation being 1.94 ± 0.02 s^-1^ (Table S1) was similar to NH_4_Cl dependent GMP formation by the MjATPPase subunit.

**Figure 2.**
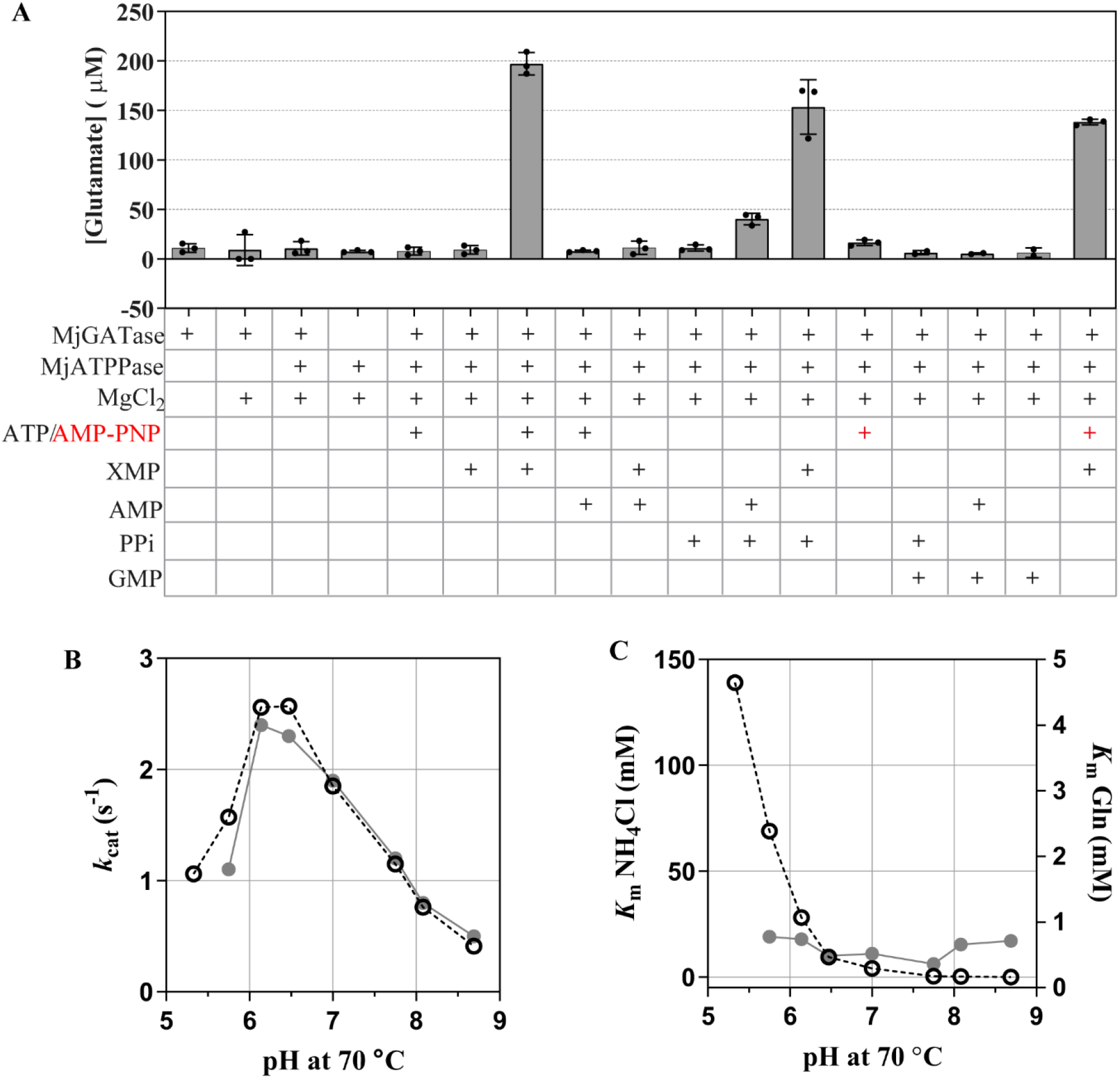
GATase activation and ammonia tunnelling in MjGMPS. (A) Screening of ligand/ligand combinations for GATase activation in MjGMPS. GATase activation was examined using an end point assay that involved measurement of Glu formed using glutamate dehydrogenase as the coupling enzyme as described in the methods section. (B and C) pH dependence of kinetic parameters *k*_cat_ (B) and *K*_m_ (C) for Gln (●) and NH_4_Cl (○) dependent GMP formation by MjGMPS. Each data point is from a Michaelis-Menten plot generated by measuring the initial velocity at various concentrations of either Gln or NH_4_Cl and saturating concentrations of XMP and ATP.Mg^2+^.

The two subunits of MjGMPS associate to form a complex strictly in the presence of ATP.Mg^2+^+XMP.^20^ The complex dissociates upon product release and hence the interaction of the two subunits is transient with the life-times of association governed by catalysis. Considering that the *k*_cat_ for Gln dependent GMP formation is 1.94 ± 0.02 s^-1^, the upper limit of the time scales over which MjATPPase-MjGATase remain associated during one catalytic cycle is 0.5 s (inverse of *k*_cat_). To ascertain if ammonia is channelled across the subunits despite this transient association, we determined the pH dependence of the Michaelis-Menten parameters for Gln and NH_4_Cl dependent GMP formation. The variation in *k*_cat_ with the two substrates followed similar trends, however, the changes in *K*_m_ were contrasting (Figure 2B, C). The *K*_m_ for NH_4_Cl dropped more than 1000-fold from 139 mM at pH 5.3 to 0.1 mM at pH 8.7 while the *K*_m_ for Gln remained unchanged across this pH range. The variation in the *K*_m_ for Gln would have followed a trend similar to the *K*_m_ for NH_4_Cl if ammonia were released out of the enzyme. Hence the ammonia generated in the MjGATase subunit is channelled to the MjATPPase subunit.

Tunnelling of ammonia can also be proven by comparing the experimentally determined *k*_cat_ for Gln dependent GMP formation with a theoretical value calculated for a scenario where ammonia is released into the solvent.^15, 19^ The experimentally determined values of *k*_cat_ and *K*_m_ for Gln dependent GMP formation at pH 6.2 is 2.4 s^-1^ and 0.7 mM, respectively, while the same parameters for NH_4_Cl dependent GMP formation are 2.5 s^-1^ and 69 mM, respectively. In an assay with a substrate concentration of 7 mM Gln (saturating), assuming that 10 % of Gln is hydrolyzed and the ammonia generated is released to the solvent, the calculated *k*_cat_ for Gln dependent activity will be 0.025 s^-1^. (2.5×0.7/69 = 0.025 s^-1^). This value of *k*_cat_ is 100-fold lower than the actual value validating that ammonia generated in the GATase subunit is indeed not released to the solvent.

### Crystal structure of MjATPPase

To dissect the structural details of the crosstalk between MjGATase and MjATPPase leading to allostery and ammonia tunnelling, structures of the independent subunits are a prerequisite. We had earlier solved the structure of the MjGATase subunit by NMR^20^ and subsequently by X-ray crystallography.^27^ Here we report the crystal structure of XMP complexed MjATPPase subunit solved to a resolution of 2.1 Å (Table 1). The model has four chains in the asymmetric unit with chains A and B, and C and D forming two homodimers, and each protomer bound to a molecule of XMP (Figure 3A and S1A). The electron density for the first 180 residues in chain C is weak and hence the residuals, Rfactor (0.23) and Rfree (0.27) are slightly higher. A maximum RMSD of 1.34 Å (Table S2) is observed upon superposing the chains indicating that the conformation of the backbone is similar across the four chains. Also, the backbone conformation of MjATPPase/XMP is similar to the apo structures of PhATPPase (PDB ID: 2DPL, 3A4I) as well as the apo and XMP bound ATPPase domains of two-domain type GMPS (Figure 3B and Table S2). However, certain flexible loops that are disordered in the structures of PhATPPase as well two-domain type GMP synthetases could be modelled in MjGMPS/XMP structure for the first time.

**Figure 3:**
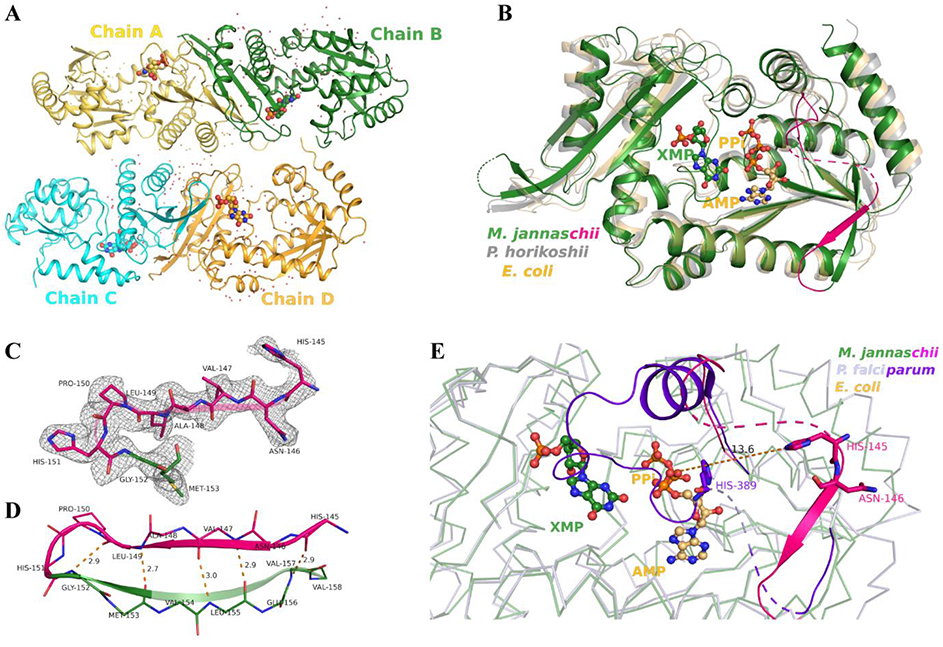
Crystal structure of MjATPPase/XMP complex. (A) The arrangement of the four chains in the asymmetric unit. The backbone is shown in cartoon representation, XMP as spheres and water molecules as dots. (B) Structural similarity among MjATPPase/XMP (chain B), PhATPPase (PDB ID: 2DPL) and the ATPPase domain of two-domain type *E. coli* GMPS (PDB ID: 1GPM). The stretch of residues Gly128-His151 in the MjATPPase/XMP structure is colored magenta to highlight that this segment is missing due to disorder in the other two overlapped structures. The dashed pink line corresponds to residues 136-144 for which electron density is missing in MjATPPase/XMP structure. (C) The fit of the residues 145-153 (chain B) to the electron density. A 2Fo-Fc simulated annealing composite omit map contoured at 1 σ is shown as mesh. (D) H-bonding between the β-strand formed by residues 145-149 (β4a) and the residues on the strand β5. (E) Superposition of the ATPPase domain of PfGMPS/Gln structure (PDB ID: 4WIO) on MjATPPase/XMP structure (chain B). The backbone is in ribbon representation and the lid-loop residues in PfGMPS/Gln (residues 365-401) and MjATPPase/XMP (residues 128-151) are in cartoon representation. The residues of the lid-loop invariant motif namely His389 of PfGMPS and His145 and Asn146 of MjATPPase are shown as sticks.

**Table 1:** Crystallographic data collection and model refinement statistics.

|  |  |
| --- | --- |
| PDB ID | 6JP9 |
| Wavelength (Å) | 1.54179 |
| Temperature (K) | 100 |
| Crystal-to-detector distance (mm) | 175 |
| Rotation range per image (°) | 1 |
| Total rotation range (°) | 190 |
| Exposure time per image (s) | 300 |
| Resolution range (Å)<sup>a</sup> | 52.53-2.10 (2.14-2.10) |
| Space group | $P2_12_12_1$ |
| Unit-cell parameters (Å) | $a = 82.18$ $b = 100.43$ $c = 157.59$ |
| Wilson B factor (Å<sup>2</sup>) | 32.1 |
| Average mosaicity (°) | 0.6 (fixed) |
| Total number of reflections<sup>a</sup> | 586519 (33959) |
| Multiplicity<sup>a</sup> | 7.6 (7.5) |
| Mean I/σ(I)<sup>a</sup> | 8.9 (1.7) |
| Completeness (%)<sup>a</sup> | 100 (100) |
| CC<sub>1/2</sub><sup>a, b</sup> | 0.989 (0.645) |
| $R_{\text{work}}$ | 0.228 |
| $R_{\text{free}}$ | 0.274 |
| RMS bond length (Å) | 0.016 |
| RMS bond angle (°) | 1.75 |
| Average B factor (Å<sup>2</sup>) | 46.0 |
| Ramachandran favoured (%) | 96.5 |
| Ramachandran allowed (%) | 3.3 |
| Ramachandran outliers (%) | 0.3 |

<sup>a</sup> Values in parentheses are for the highest resolution shell. <sup>b</sup> CC<sub>1/2</sub> is the Pearson's correlation coefficient of the intensity estimates between two random halves of a data set. $R_{\text{work}} = \Sigma ||F_{\text{obs}}| - |F_{\text{calc}}|| / \Sigma |F_{\text{obs}}|$ , $R_{\text{free}}$ is $R_{\text{work}}$ calculated for 5% reflections that were randomly chosen and omitted from the use in refinement.

MjATPPase subunit is composed of a N-terminal ATPPase core domain (residues 1-184) which consists a five stranded parallel β-sheet surrounded by seven α-helices (Figure S1B, C). This domain harbours residues involved in binding ATP.Mg^2+^ as well as the catalysis of AMP-XMP intermediate formation. These residues are primarily located on a ∼25 residue long loop, termed the lid-loop, which connects the strands β4 and β5 (Figure S1B). In MjATPPase, the lid-loop spans the residues 128-151. The corresponding stretch of residues is disordered in the structures of PhATPPase while in two-domain type GMP synthetases, except for the N-terminal stretch which is modelled as a helix (MjATPPase residues 131-137), the remaining portion of the lid-loop is disordered (Figure S2A, B). In MjATPPase/XMP structure, the electron density for the lid-loop barring residues 136-144 is evident revealing the conformation of this segment (Figure 3B, C). The conformation of the N-terminal stretch of the lid-loop (residues 128-135) varied among the chains (Figure S3) while the fit of the residues His145-Gly151 to the electron density is good (Figure 3C). The polypeptide stretch His145-Leu149 forms a β-strand (β4a) which is antiparallel to the core β-sheet formed by strands β1-β5 (Figure 3B). The conformation of this antiparallel β-strand is stabilised by five H-bonds with the adjacent strand β5 (Figure 3D). Interestingly, His145, whose counterpart in PfGMPS is crucial for catalysis, is positioned distal from the substrates (Figure 3E).

The dimerization domain (residues 202-310) that is connected to the ATPPase core domain by a 17-residue long linker (residues 185-201) consists of a mixed four-stranded β-sheet (β6-β9) and two α-helices that are placed on one side of the β-sheet (Figure S1D, E). At the dimer interface, residues from the β-sheets of adjacent monomers interact burying an area of ∼1600 Å^2^. This apart, the two monomers in the MjATPPase/XMP structure are cross-linked by a disulphide bond involving Cys239 (Figure 4A). The position corresponding to Cys239 is not conserved barring a few sequences from the genus *Methanocaldococcus* (Figure S4) in which the ATPPase subunits might also be tethered by a disulphide bond. The dimerization domain is also important for ligand binding as residues on a loop at the C-terminal extreme and inter-dimer interactions of a conserved Arg residue are crucial for binding XMP.^28^ In MjATPPase/XMP structure, the interactions of the residues on the C-terminal loop (residues 298-310) with XMP as well as the inter-dimer interactions of Arg294 are conserved (Figure 4B).

**Figure 4:**
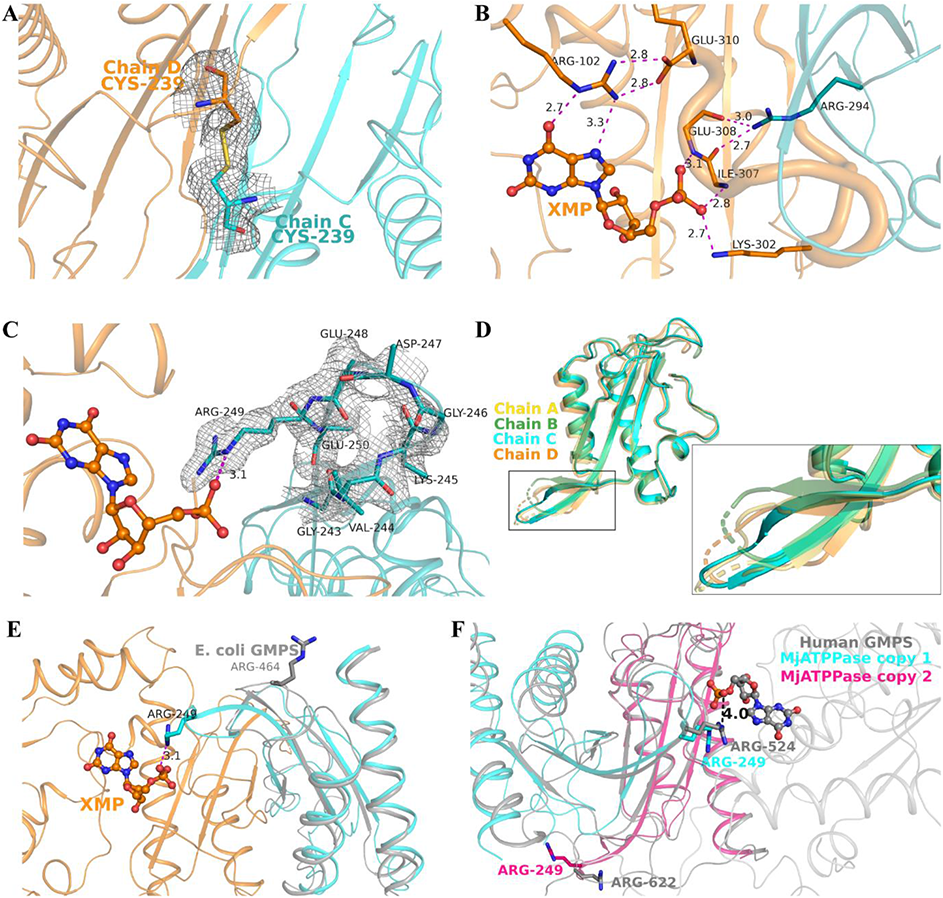
Role of dimerization domain in binding XMP. (A) Inter-dimer disulphide bond between chains C and D. A 2Fo-Fc simulated annealing composite omit map contoured at 1 σ is shown as mesh. The protein backbone is shown as cartoon and residues as sticks. (B) Interactions of residues on the C-terminal loop (tube representation) with XMP and the inter-dimer interactions of Arg294. XMP is shown in ball-and-stick representation. (C) Fit of residues on the loop connecting β7-β8 to the electron density in chain C. The interaction of Arg249 with XMP bound to chain D is also shown. (D) Structural superposition of the dimerization domains in the four chains of MjATPPase. A magnified view of the boxed region is shown in the inset. (E) Conformation of the residue corresponding to Arg249 in *E. coli* GMPS. The ATPPase domain of *E. coli* GMPS was superposed on chain C of MjATPPase/XMP structure. (F) Conformation of the residue corresponding to Arg249 in human GMPS. The dimerization domain of MjATPPase/XMP chain C, was superposed twice, each time on a different dimerization sub-domain of human GMPS (PDB ID: 2VXO). The MjATPPase/XMP structure colored cyan was superposed on the dimerization sub-domain corresponding to the segment 458-563 of human GMPS.

Another novel observation in the MjATPPase/XMP structure is a salt-bridge like interaction between the XMP phosphate and the side chain of Arg249 from the neighbouring chain (Figure 4C). Arg249 is in a loop that connects two antiparallel β-strands (β7-β8) and many residues on this loop including Arg249 are invariant across two-subunit as well as two-domain type GMP synthetases (Figure S4). The loop constituted by residues Gly243-Glu250 is largely disordered and except for chain C, which has strong electron density for these residues, other three chains lack the electron density (Figure 4C, D). The residues on this loop are also disordered in the structures of two-domain type GMPS from *P. falciparum*, *C. burnetii* and *T. thermophilus* suggesting that this loop is conformationally dynamic (Figure S5). Interestingly, in the structures of *E. coli* as well as *N. gonorrhoeae* GMPS which lack a bound XMP, the Arg residue corresponding to Arg249 of MjATPPase takes up a conformation that is far away from the active site of the neighbouring protomer (Figure 4E). Human GMPS is active as a monomer since it harbours an insertion that mimics the arrangement of two dimerization domains in homologs that are dimers.^29^ Owning to the duplication, there are two Arg residues in human GMPS corresponding to Arg249 of MjATPPase. Among the two, the Arg residue in the insertion interacts with XMP in a manner identical to the interaction of Arg249 in MjATPPase/XMP structure. The important difference being that the interaction is with XMP bound to the same chain (Figure 4F).

### Trapping the transient MjGMPS complex using covalent cross-linking

Since the MjGMPS subunits interact transiently, we attempted to trap the complex using covalent cross-linking. We incubated a reaction mix containing equimolar concentrations of MjGMPS subunits and saturating concentrations of ATP.Mg^2+^+XMP at 70 °C for 2 min prior to addition of the homobifunctional cross-linker bis(sulfosuccinimidyl)suberate (BS3), which reacts predominantly with the primary amine groups of Lys residues and the protein N-terminus. MjATPPase alone in the presence/absence of substrates and MjATPPase+MjGATase in the absence of substrates served as controls. The cross-linked species were separated by reducing SDS-PAGE and the results are shown in Figure 5. Interestingly, in the control reaction containing MjATPPase subunit alone (absence/presence of substrates), monomeric species as well as the dimer were observed although the protein is a dimer in its native state.^20^ This suggests that under the experimental conditions only a fraction of the dimeric species is covalently cross-linked. In the cross-linking reaction containing both the subunits, two distinct cross-linked species were observed only in the presence of substrates. Examination of the migration pattern suggested that the band migrating with an apparent molecular mass of ∼100 kDa corresponds to a heterotetramer comprising two subunits each of MjGATase and MjATPPase while the species migrating at ∼60 kDa is a heterodimer. The heterotetramer is the functional form since each chain of dimeric MjATPPase interacts with one subunit of MjGATase.^20^ The heterodimer is observed since a significant fraction of the MjATPPase dimers fail to cross-link as observed in the control reaction. The populations of the hetero oligomers were insignificant in the absence of substrates. These experiments were carried out three times yielding similar results (Figure S6). Thus, covalent cross-linking enabled us capture the functional MjGMPS complex.

**Figure 5:**
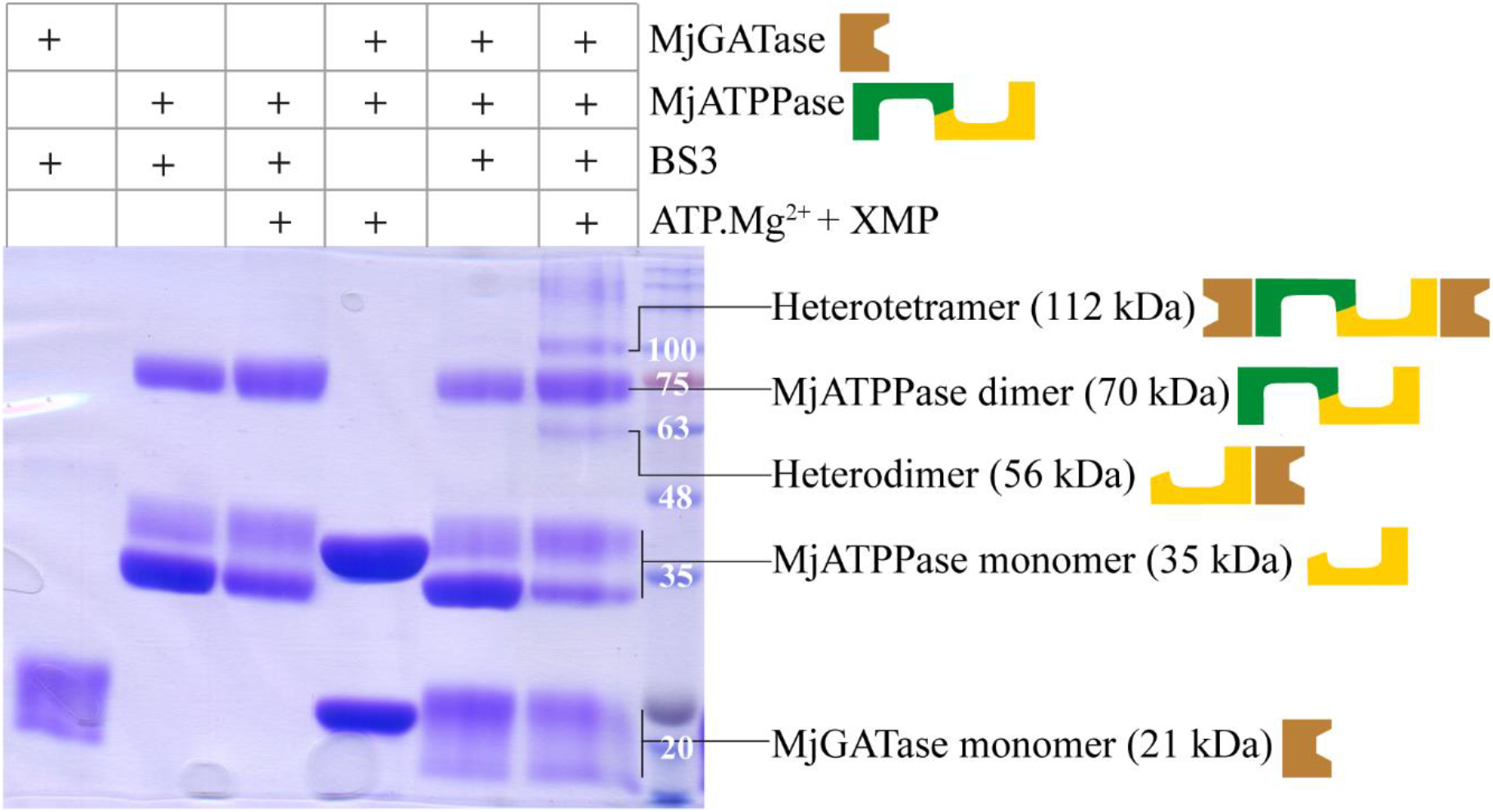
Covalent cross-linking of MjGMPS using BS3. 50 µM each of MjGATase and MjATPPase were incubated with 0.2 mM XMP, 2 mM ATP and 20 mM MgCl_2_ at 70 °C for 2 min before cross-linking with BS3 at 50 °C for 5 min. The reaction was quenched by adding SDS-PAGE loading dye and the bands were resolved by 12 % (w/v) SDS-PAGE. The bands were visualized by Coomassie Brilliant Blue staining. The protein molecular weight standards are in the last lane with the molecular weight in kDa. This is a representative gel image of three independent experiments.

### Identification of cross-linked residues using LC-MS/MS

The bands corresponding to the heterodimer and the heterotetramer were excised, subjected to in-gel tryptic digestion and the tryptic peptides were analysed by LC-MS/MS. The tandem mass spectra were processed using the program pLink2 to identify the cross-linked residues. False positives were omitted by including only the cross-linked peptides that were consistent across replicates (for details see methods section). The cross-links observed in the heterodimer and the heterotetramer samples were classified as intrachain MjGATase or intrachain MjATPPase cross-links (referred to in the following section as intralinks), inter-dimer cross-links across the two chains of the MjATPPase dimer and the inter-subunit cross-links across the MjGATase-MjATPPase subunits. We observed 16 unique MjGATase intralinks in the heterodimer as well as the heterotetramer with 12 common to the two sets (Figure 6A, Table S3 and Supplementary file 2). Structural validation of these intralinks was performed by measuring the Euclidean distances between the Cα atoms of the cross-linked residues in the apo structure of MjGATase (PDB ID: 7D40). The distances of the 16 GATase intralinks were within the cut-off limit of 30 Å imposed by the length of the cross-linker arm (Figure 6B, Table S3).^30^ The absence of intralinks exceeding the distance cut-off validated our experimental approach as well as the analysis workflow and avoided false-positives. With regard to the MjATPPase subunit, we identified 26 unique intralinks in the heterodimer and the heterotetramer, with 11 being common (Figure 6A, Table S4, Supplementary file 2). Among the 26 unique intralinks, the structural validity of the cross-links 142-302 and 103-140 could not be ascertained as they involve residues on the disordered lid-loop while the intralink 1-61 was an over-length cross-link (Figure 6A, C, Table S4). Of the remaining 23 unique intralinks, 21 were valid intralinks while the cross-links 245-302 and 166-245 observed in the heterotetramer samples, were valid only as inter-dimer cross-links connecting the two chains of the MjATPPase dimer.

**Figure 6:**
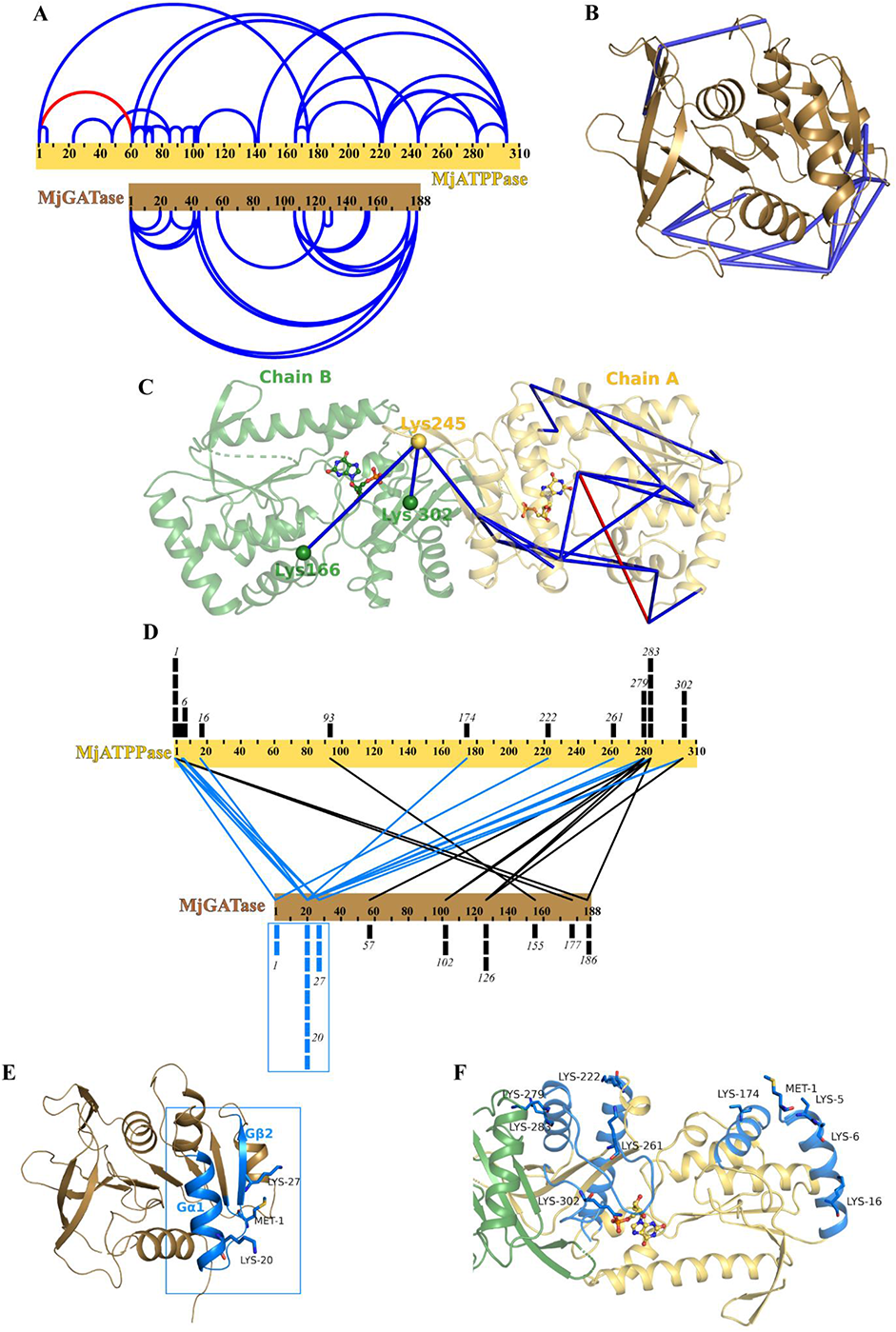
Identification and structural validation of cross-links. The cross-links mapped on (A) the length of the protein sequences, the crystal structures of (B) MjGATase and (C) MjATPPase dimer. In panels B and C, the protein backbone is shown in cartoon representation, the cross-links as lines and the disordered region as dashes. The Euclidean distances between the Cα atoms of cross-linked residues are within the distance threshold (30 Å) except for the cross-link 1-61 (36.5 Å), which is colored red. The Cα atoms of residues involved in MjATPPase inter-dimer cross-links are shown as spheres and the residues are labelled. (D) The MjGATase-MjATPPase inter-subunit cross-links mapped across the lengths of the two protein sequences. Cross-links are shown as rectangles with each rectangle indicating a unique cross-link and the residue is labelled. The cross-links involving the first 27 residues of MjGATase subunit are colored blue and the secondary structural elements that take part in these cross-links are highlighted in the crystal structures of MjGATase (E) and MjATPPase (F) using the same color scheme. The cross-linked residues are shown as sticks and XMP is shown in ball and stick representation (C, F).

A total of 24 unique MjGATase-MjATPPase inter-subunit cross-links were identified in the heterodimer and heterotetramer, with 10 cross-links common across both the samples (Figure 6D, Table 2 and Supplementary file 2). The tandem mass spectra of the cross-linked peptides were manually inspected for the presence of fragment ions that permit unambiguous identification of the cross-linked residues (Supplementary files 3A and 3B). The N-terminal amino group of MjATPPase was cross-linked with the N-terminal amino group as well as 6 different Lys residues of MjGATase (Table 2). In the MjATPPase sequence, the cross-links were distinctly clustered either to the N-terminal region (9 cross-links), or the C-terminus (11 cross-links) (Figure 6D). Whereas in the MjGATase sequence, 14 cross-links clustered to 27 residues at the N-terminus and the remaining were distributed throughout the sequence. In the MjGATase crystal structure, these 14 cross-links map to helix Gα1 and β-strand Gβ2 at one end of the core β-sheet (Figure 6E). The MjATPPase counterparts of these 14 cross-links are located on helices α1 and α6 of the MjATPPase core domain as well as various regions of the dimerization domain (Figure 6F). These observations suggest that the MjGATase subunit interacts primarily through Gα1 and Gβ2 at its N-terminus, and this region interacts with both the core and dimerization domains of MjATPPase.

**Table 2:** List of inter-subunit MjATPPase and MjGATase cross-links.

| Serial Number | Cross-linked residues (MjATPPase-MjGATase) | Present in heterodimer | Present in heterotetramer | Nature of cross-link |
| --- | --- | --- | --- | --- |
| 1 | 1-1 | Yes | Yes | N-terminus – N-terminus |
| 2 | 1-20 | Yes | Yes | N-terminus – Lys |
| 3 | 1-27 | Yes | Yes | N-terminus – Lys |
| 4 | 1-177 | Yes | Yes | N-terminus – Lys |
| 5 | 1-186 | Yes | Yes | N-terminus – Lys |
| 6 | 5-20 | No | Yes | Lys – Lys |
| 7 | 6-20 | Yes | Yes | Lys – Lys |
| 8 | 6-27 | Yes | No | Lys – Lys |
| 9 | 16-20 | Yes | Yes | Lys – Lys |
| 10 | 93-155 | No | Yes | Lys – Lys |
| 11 | 174-20 | Yes | Yes | Lys – Lys |
| 12 | 222-1 | No | Yes | Lys – N-terminus |
| 13 | 261-20 | Yes | No | Lys – Lys |
| 14 | 279-20 | Yes | No | Lys – Lys |
| 15 | 279-102 | Yes | No | Lys – Lys |
| 16 | 279-126 | Yes | Yes | Lys – Lys |
| 17 | 283-20 | Yes | No | Lys – Lys |
| 18 | 283-57 | Yes | No | Lys – Lys |
| 19 | 283-102 | Yes | No | Lys – Lys |
| 20 | 283-126 | Yes | No | Lys – Lys |
| 21 | 283-186 | Yes | No | Lys – Lys |
| 22 | 302-20 | Yes | Yes | Lys – Lys |
| 23 | 302-27 | Yes | No | Lys – Lys |
| 24 | 302-126 | Yes | No | Lys – Lys |

### Integrative modelling of the MjGMPS complex

We used the inter-subunit cross-links as distance restraints to compute and visualize the MjGATase-MjATPPase interaction space using the exploratory modelling tool DisVis.^31^ This tool performs a six-dimensional docking search of the scanning chain (MjGATase) around a fixed chain (MjATPPase dimer) and reports the number of conformations of the scanning chain that is consistent with increasing number of restraints, as well as the number of times a restraint is violated. DisVis analysis using 24 cross-links as restraints flagged 12 cross-links as outliers based on high Z-scores (Table S5) while 99.6% of the complexes were consistent with the remaining 12 cross-links (Figure 7A and Table S6) suggesting that a subset of 12 cross-links captures a distinct conformation of MjGATase-MjATPPase complex. To validate the same, we divided the 24 cross-links into two subsets, Subset 1 and Subset 2, with the latter containing the 12 cross-links that were flagged for high Z-scores in the first analysis. While DisVis analysis using Subset 1 identified complexes consistent with all 12 cross-links, complexes consistent with only 9 cross-links were identified using Subset 2 (Figure 7B and Tables, S5, S6). The interaction space captured by Subset 1 maps to an area primarily surrounding the MjATPPase core domain (Figure 7C). This is distinct compared to the area traversed by the MjGATase subunit when the 9 valid cross-links of Subset 2 are considered. Thus, exploratory modelling revealed the possibility of two modes to the interaction of MjGATase-MjATPPase.

**Figure 7:**
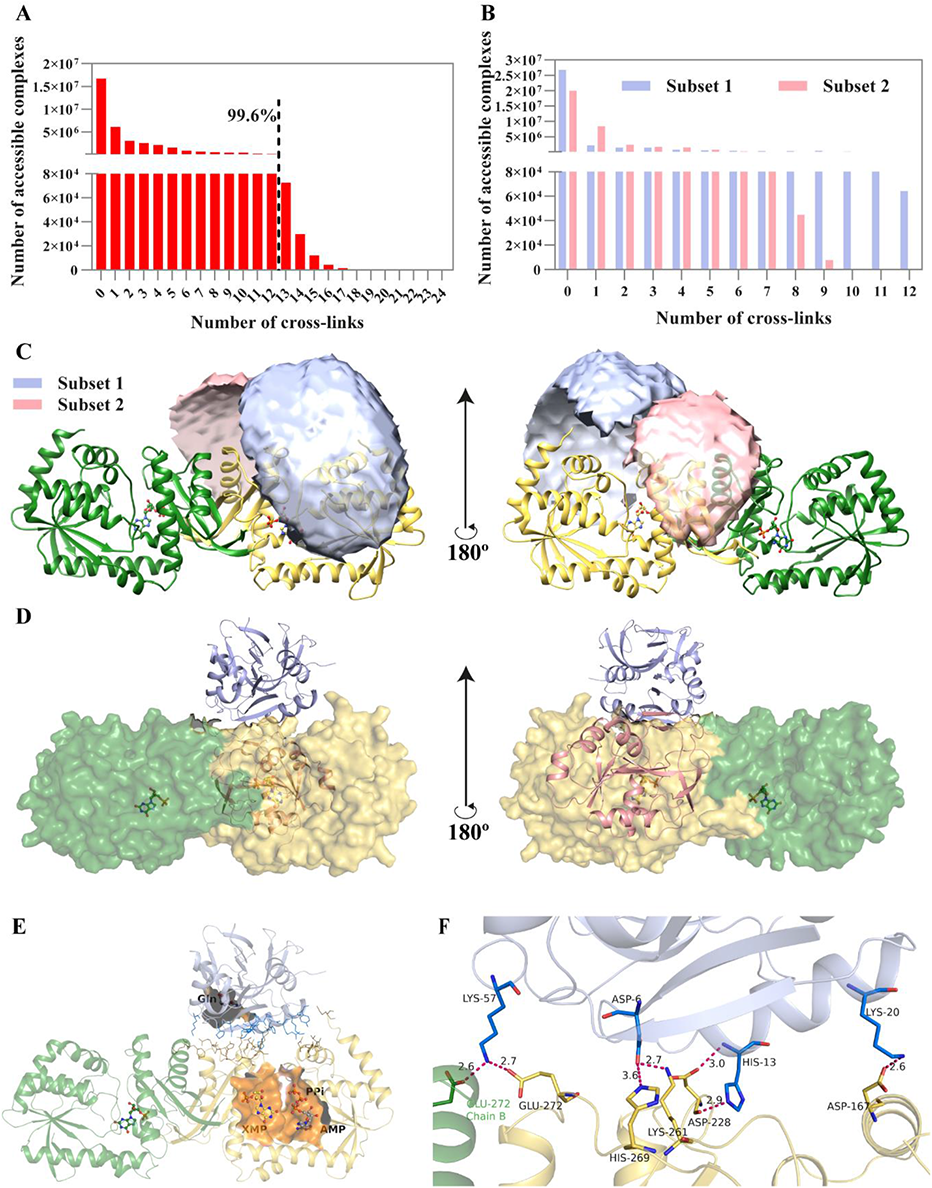
XL-MS guided modelling of the MjGMPS complex. Exploratory modelling of the MjGMPS complex using (A) 24 cross-links and (B) cross-links of Subsets 1 and 2, as distance restraints. (C) Visualization of the MjGATase-MjATPPase interaction space calculated using the cross-links of Subsets 1 and 2. MjATPPase backbone is shown as cartoon and XMP is in ball and stick representation. (D) HADDOCK modelled structures of the MjGMPS complex generated using the cross-links of Subsets 1 and 2 as distance restraints. The ATPPase subunits from the highest scoring models were superposed on each other. (E) The MjGATase-MjATPPase interaction space in the model generated using Subset 1. The residues within 4 Å of each subunit are shown as lines. The ATPPase ligands AMP, PPi and the GATase ligand Gln are superposed from the crystal structures of *E. coli* GMPS (PDB ID: 1GPM) and *P. falciparum* GMPS (PDB ID: 4WIO), respectively. The active sites of the two subunits are shown in surface representation. (F) A zoomed-in view of the inter-subunit salt bridges. Color coding of the GATase subunit is uniform across the panels B-F.

To analyze the molecular details of subunit crosstalk, an experimentally driven model of the MjGATase-MjATPPase complex was generated using HADDOCK 2.4 webserver.^32^ The structures of MjATPPase (dimer) and MjGATase subunits were docked against each other guided by the set of 24 cross-links, as well as the two subsets, Subset 1 and Subset 2, that were grouped based on DisVis analysis. HADDOCK generates 200 water-refined models and clusters them based on interface RMSD, a measure that captures the conformational changes about the interface. The top cluster of models generated using the cross-links of subset 1 had the lowest HADDOCK score (−124.3 ± 2.5) compared to the top cluster generated with all 24 cross-links (−60.2 ± 0.8) or subset 2 (−82.4 ± 2.2) (Table S7). The best models (ones with the lowest HADDOCK score) in the top clusters of the three docking runs, were then evaluated to determine how well they satisfy the XL-MS data. The model generated using Subset 1 which was the best in terms of HADDOCK score, also had the distances between the Cα atoms for the 12 cross-links within the cut-off limit (Table S8). Unlike this, many cross-links exceeded the limits in both the other cases.

In the model generated using Subset 1, the MjGATase subunit is positioned atop the dimerization domain while it is positioned on the rear side of the MjATPPase active site in the model generated using Subset 2 (Figure 7D). Analysis of the model from Subset 1 revealed that the MjGATase residues on the helix Gα1 and strand Gβ2 at one extreme of the seven stranded β-sheet interact with residues on the MjATPPase core as well as dimerization domains (Figure 7E). Further examination revealed the presence of several inter-subunit salt bridges with a majority of them involving the MjATPPase core domain (Figure 7F). Although the model (Figure S7) generated with all 24 cross-links has the poorest HADDOCK score, the MjGATase subunit is positioned atop the MjATPPase dimerization domain but in a position distinct from that of Subset 1.

### Fusing the two subunits of MjGMPS results in an active, two-domain type enzyme

Alignment of the sequences of the two types of GMP synthetases showed the absence of additional linker sequence in the two-domain type enzyme (Figure 8A) suggesting that a fused MjGMPS may be functional. To probe this possibility, we fused the MjATPPase subunit to the C-terminus of MjGATase rendering it a two-domain type GMP synthetase and used the fused enzyme for biochemical studies. Fused MjGMPS is a dimer in solution similar to MjATPPase subunit, which is not surprising considering that dimerization is mediated by the ATPPase subunit (Figure 8B, S8). Tethering the two subunits did not activate the GATase domain and hydrolysis of Gln was observed only in presence of ATP.Mg^2+^ and XMP (Figure 8C). The pH and temperature dependence of Gln and NH_4_Cl dependent GMP formation in fused MjGMPS were largely similar to MjGMPS (Figure 1B, C). The *k*_cat_ for Gln dependent GMP formation was estimated to be 0.6 ± 0.03 s^-1^, a value that is 3-fold lower compared to MjGMPS. The *K*_m_ for Gln (1.4 ± 0.3 mM) was 2.7-fold higher while the *K*_m_ for ATP.Mg^2+^ and XMP remained unaltered compared to MjGMPS (Table S1). Thus, fusing the two subunits of MjGMPS without any intervening linker does not impede the function. Our earlier studies on PfGMPS have established the occurrence of large-scale domain reorganization arising from a 85° rotation of the GATase domain that plays a role in catalysis.^33^ Hence, it is likely that the functional mechanism of two-domain type and two-subunit type GMP synthetases is conserved.

**Figure 8:**
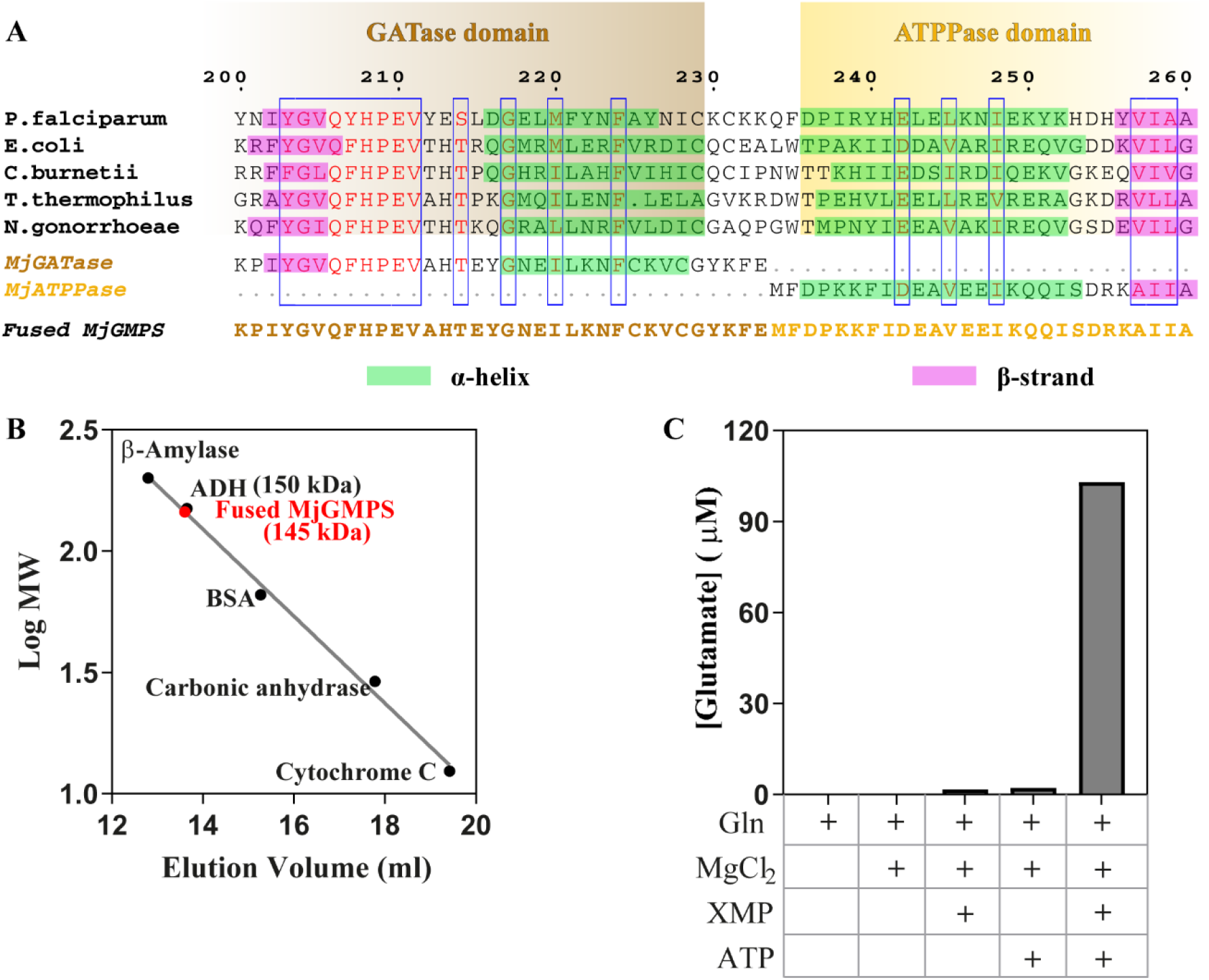
Sequence alignment and biochemical characterization of fused MjGMPS. (A) Multiple sequence alignment of MjGATase, MjATPPase and sequences of two-domain type GMP synthetases whose structures are available. The numbering is for PfGMPS sequence. Conserved residues are colored red and boxed in blue. The sequence of fused MjGMPS is also shown. (B) Analytical size-exclusion chromatography experiment to determine the oligomeric state of fused MjGMPS. The elution volume of fused MjGMPS corresponds to a dimer. (C) Substrate screen for GATase activation using an end point assay that involved measuring the Glu formed using glutamate dehydrogenase as the coupling enzyme.

## Discussion

### High affinity for ammonia suggests that MjGMPS is at the crossroads of evolution

Considering that Gln amidotransferases can only utilize ammonia (NH_3_) and not ammonium (NH_4_ ^+^) and the p*K*_a_ for NH_4_ ^+^ — NH_3_ + H^+^ is ∼9.2, the NH_4_Cl dependent activity increases with increase in pH values above 7.0 with an optimum activity around pH of 8-8.5. However, pH optimum of 7.0 for NH_4_Cl dependent GMP formation is a distinct feature of MjATPPase, which is significantly lower in comparison to 8.3 for *E. coli* GMPS^15^ or 9.2 for PfGMPS^19^. Apart from the low pH optimum, even the magnitude of the *K*_m_ value for NH_4_Cl (4.1 ± 0.2 mM) is the least for MjATPPase compared to the values of 103 mM for *E. coli* GMPS^26^, 174 mM for human GMPS^16^, 26 mM for *M. tuberculosis* GMPS^18^ and 10.8 mM for PfGMPS^17^ when estimated at the corresponding pH optima. The pH optimum of 7 and the smaller *K*_m_ value suggests that MjATPPase has higher affinity for ammonia compared to its homologs, a feature that has relevance to the evolution of Gln amidotransferases. In the case of NAD synthetase, a Gln amidotransferase, the genome of *M. jannaschii* lacks a cognitive GATase subunit that can associate with the synthetase subunit. However, the synthetase subunit is capable of utilizing ammonia *in vitro*. Based on phylogenetic analysis, it was established that *M. jannaschii* NAD synthetase represents an ancestral Gln amidotransferase that utilizes ammonia directly from the cellular milieu^34^. Hence, the observations with MjGMPS raise the intriguing possibility that this enzyme is at an intermediate state of evolution where it has not lost its ability to efficiently utilize ammonia, but has evolved far enough to utilize Gln by associating with a GATase subunit.

### Complex formation underlies a unique allosteric mechanism in MjGMPS

Since extensive domain/subunit crosstalk underlies the function of Gln amidotransferases, the GATase and the acceptor subunits form a complex even in the absence of substrates.^9, 21–25^ In contrast, the MjGATase and MjATPPase subunits even in the presence of the ATP.Mg^2+^+XMP, form only a transient complex, a departure from Gln amidotransferases studied thus far. In addition to size-exclusion chromatography and pull-down experiments, the transient interaction of the MjGMPS subunits has also been validated by NMR experiments which places the interaction in the intermediate to fast exchange regimes of NMR time scales.^20^ The complex remains intact over a catalytic cycle with the upper limit for the time-scales of association being 0.5 s, as determined from the steady state *k*_cat_ value. This observation of a strict ligand-dependent association is indicative of extremely tight regulation of Gln hydrolysis. While conformational changes induced by substrate binding underlies allosteric activation in Gln amidotransferases that pre-exist as a complex, in MjGMPS however, formation of a complex is conditional to ligand binding, that in-turn leads to allostery. Further, despite the transient nature of subunit interaction, biochemical experiments reported here provide unequivocal evidence for ammonia tunnelling.

### Crystal structure

Observations in the MjATPPase/XMP structure reported here reveal novel insights into the role of conformationally dynamic elements in the ATPPase core and dimerization domains. In two-domain type GMP synthetases, the ATPPase core domain is involved in binding the substrates, catalysing the formation of AMP-XMP intermediate as well as interacting with and activating the GATase domain. An earlier study using site-directed mutants of PfGMPS has shown that residues on the lid-loop and the subsequent helix (α6 of MjATPPase, residues 166-176) are crucial for the various functions of the ATPPase core domain.^28^ Following substrate binding, the lid-loop closes on the active site and positions the residues of the lid-loop invariant motif ^141^IK(T/S)HHN^1^^46^ (MjATPPase numbering) in proximity with the substrates.^28, 33^ Since the lid-loop is conformationally dynamic, it is disordered in a majority of the crystal structures (Figure S2); however, in the MjATPPase/XMP structure, we could model the residues His145 and Asn146 of the invariant motif (Figure 3C, D and E). The corresponding residues in PfGMPS play roles in catalysing the synthesis of AMP-XMP intermediate and GMP, respectively.^28^ Considering that His145 and Asn146 are positioned > 13 Å away from the bound XMP, the conformation of the lid-loop in MjATPPase must correspond to an inactive or resting state of the enzyme. Thus, subsequent to the binding of ATP.Mg^2+^+XMP, conformational changes in the lid-loop must reposition His145 and Asn146 in proximity to the substrates. It is possible that these changes render the MjATPPase subunit competent to associate with the MjGATase subunit.

The mode of dimerization in MjATPPase/XMP structure, as well as in PhGMPS is similar to two-domain type GMP synthetases which dimerize through their ATPPase domains.^14, 33, 35^ The role of the dimerization domain was probed in PfGMPS where the interactions of the residues on the C-terminal loop with XMP were deemed crucial for substrate binding.^28^ Moreover, the conformation of the 13-residue long C-terminal loop was stabilized by H-bonds with the side chain of Arg539 (PfGMPS numbering) located in the neighbouring chain. Mutation of Arg539 rendered the enzyme inactive on account of impaired substrate binding underscoring the importance of inter-dimer interactions for enzyme function.^28^ In MjATPPase/XMP structure, interactions of the residues on the C-terminal loop with XMP as well as the inter-dimer interactions involving Arg294 (corresponding to Arg539 of PfGMPS) are retained (Figure 4B) suggesting a conserved mechanism of binding XMP. Apart from this indirect interaction, the MjATPPase/XMP structure enables the visualization of a direct salt-bridge of Arg249 with the phosphate of XMP bound to the neighbouring chain (Figure 4C). Arg249 is invariant in the sequences of two-subunit and two-domain type enzymes suggesting that the inter-dimer interactions involving this residue are conserved (Figure S4). Moreover, in GMPS structures from *E. coli* and *N. gonorrhoeae*, which lack a bound XMP, the side chain of the residue corresponding to Arg249 is oriented towards the same chain rather than the neighbouring chain (Figure 4E). Thus, the binding of XMP triggers a conformational change in the loop bearing Arg249 by more than 9 Å leading to stronger substrate binding.

The observations in the MjATPPase/XMP structure also provide rationale for how human GMPS is active as a monomer.^16^ The latter enzyme has an insertion of 130 residues that essentially duplicates the dimerization domain and this additional copy mimics the arrangement of two dimerization domains in dimeric GMP synthetases.^29^ Structural superposition of the dimerization domain of MjATPPase/XMP on the insertion in human GMPS reveals that Arg524 in the latter structure occupies the position corresponding to Arg249 of MjATPPase and interacts with the XMP bound to the same chain (Figure 4F). Thus, the insertion in human GMPS substitutes for the interactions of the dimerization domain from the neighbouring chain thereby enabling the human enzyme to function as a monomer.

### Mechanism of allostery and catalysis in MjGMPS

In 9 out of the 10 crystal structures of the well-studied two-domain type GMP synthetases (PDB IDs: 1GPM, 3TQI, 2YWB, 2YWC, 5TW7, 2VXO, 3UOW, 4WIM, 7SBC), the GATase domain is oriented in the 0° rotated state and the inter-domain interface is constituted of α1, β2 of the GATase domain and the helices α11 and α12 of the ATPPase domain (Figure 9A). The amino acid sequences of the interface helices are highly conserved, and the conserved sequence motifs are enriched with charged residues (Figure S9A).^28^ Further, residues of the conserved motif of GATase helix α1 form inter-domain salt-bridge interactions with residues on the ATPPase helices α11 and α12 motifs (Figure S9B). Perturbing these inter-domain interactions in PfGMPS by mutagenesis revealed that the α1-α12 inter-domain salt-bridge interactions are conduits for information exchange pertaining to allostery through which the catalytic events in the ATPPase domain are communicated to the GATase domain.^28^

**Figure 9:**
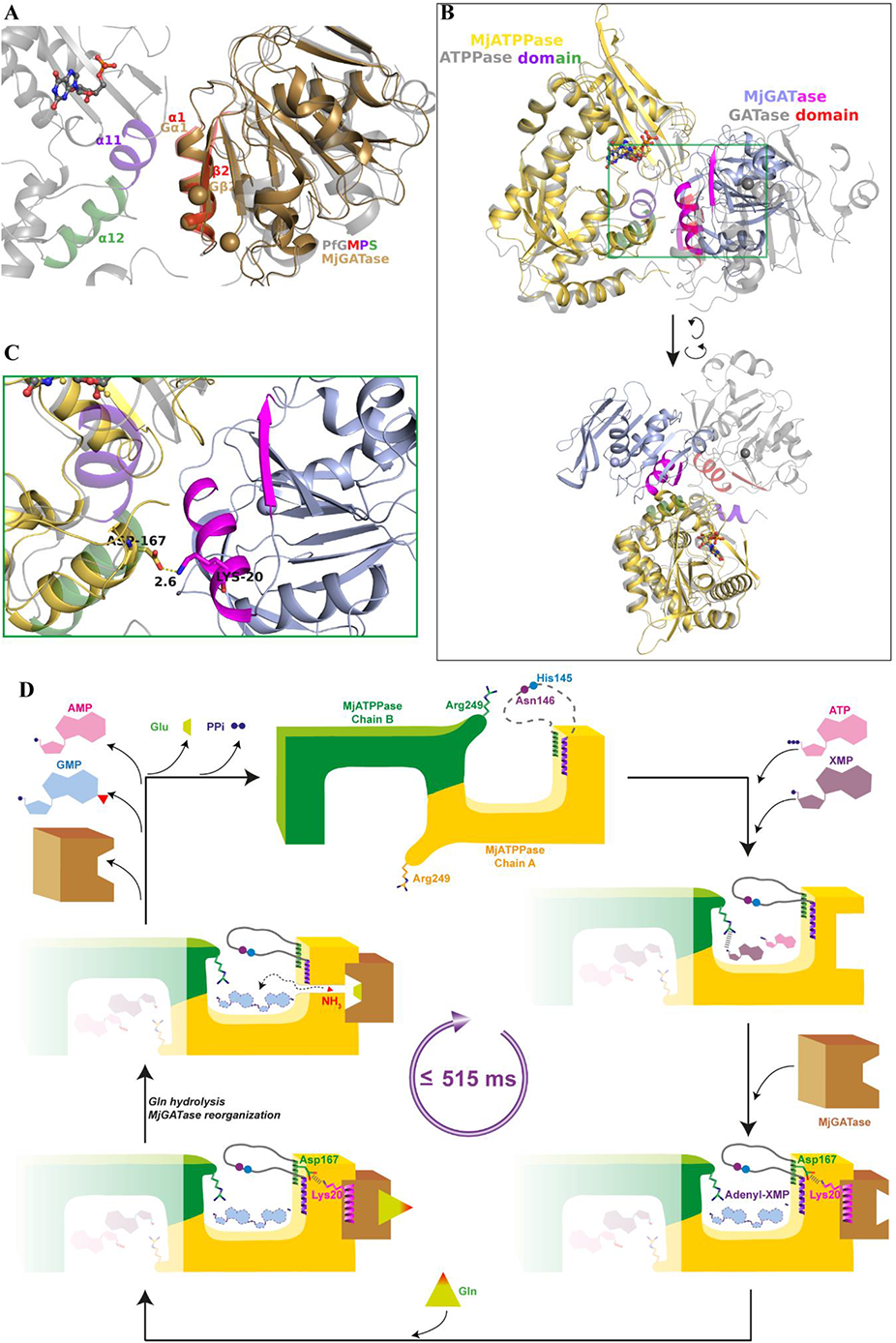
Structural basis for allosteric activation of MjGATase subunit. (A) Interdomain interface in two-domain type GMP synthetases with GATase in the 0° rotated state as exemplified by the PfGMPS/XMP structure. The MjGATase structure (PDB ID: 7D40) is superposed on the GATase domain of PfGMPS/XMP. The N-terminal residues of MjGATase namely 1, 20 and 27 which are involved in 14 inter-subunit cross-links are shown as spheres. (B) Structural superposition of the HADDOCK model of MjGMPS (Subset 1) with PfGMPS/XMP structure. The catalytic Cys residues of the GATase domain and subunit are shown as spheres to indicate the orientation of the active site. The portion of the image within the green box is blown-up and shown in (C). The inter-subunit interaction is shown and the GATase domain of PfGMPS/XMP is removed to declutter the image. (D) Schematic explaining the functional scheme of MjGMPS. It should be noted that MjGATase binding is not conditional to the formation of AMP-XMP.

Owning to the transient nature of the MjGMPS complex, we resorted to XL-MS and integrative modelling to uncover the structural basis of allostery. Among the 24 inter-subunit cross-links, 14 of them involved residues on the helix Gα1 and the strand Gβ2 of the MjGATase subunit. These elements, which correspond to GATase helix α1 and strand β2 of two domain type GMP synthetases, are also the components of the 0° inter-domain interface (Figure 9A) suggesting that the interactions of the MjGATase subunit with the MjATPPase subunit bears features of the 0° rotated state of the GATase domain. Superposing the HADDOCK model of the MjGMPS complex generated using Subset 1, with the PfGMPS/XMP structure (PDB ID: 3UOW) further validated that the mode of interaction of the MjGATase subunit has features similar to the interaction of the GATase domain in the 0° rotated state (Figure 9B). Moreover, identical to the α1-α12 inter-domain salt-bridge interactions in structures of two-domain type GMP synthetases, the MjGMPS model revealed an inter-subunit salt bridge between Lys20 on helix Gα1 of MjGATase and Asp167 on helix α6 of MjATPPase (Fig. 9C and 7F). Since these inter-domain salt bridge interactions are conserved in the structures of two-domain type GMP synthetases (Figure S9), and moreover proven to be critical for GATase activation, XL-MS guided integrative modelling has successfully revealed a high-resolution structural snapshot of the MjGMPS complex in which the ATP.Mg^2+^+XMP bound MjATPPase subunit interacts with and allosterically activates the MjGATase subunit.

Interestingly, in the 0° structures of two-domain type GMP synthetases, the GATase active site is exposed to the solvent and a tunnel linking the two active sites is not apparent in the apo as well as the ligand-bound structures (Figure S10). Hence, the structural basis of the tight regulation of GATase activity and ammonia tunnelling were unresolved despite the biochemical evidence for these features. This conundrum was resolved to an extent by the crystal structure of Gln-complexed PfGMPS which displayed an 85° rotation of the GATase domain (PDB ID: 4WIO) and brought forth the possibility of active site coupling mediated by a large-scale domain rearrangement (Figure S10).^33^ Further, molecular dynamics simulations suggested that ammonia is channelled at an intermediate rotated state of the GATase domain. The retention of all catalytic properties in fused MjGMPS, including the dimeric state, allostery and ability to utilize Gln validates that the functional mechanism of MjGMPS is similar to two-domain type enzymes including the reorganization of the MjGATase subunit.

The results of the biochemical experiments, crystal structure and XL-MS guided integrative modelling reported here, along with our earlier understanding of PfGMPS^19, 28, 33^ permit us to converge on a functional model for MjGMPS (Figure 9D). In the apo-state, the lid-loop of MjATPPase is disordered and the side chain of Arg249 is oriented towards the chain that harbours the residue. Binding of ATP.Mg^2+^ and XMP triggers a conformational change in Arg249 leading to the formation of a salt bridge with the phosphate of XMP bound to the neighbouring chain. The lid-loop closes on the active site, repositioning the catalytic residues His145 and Asn146 by more than 15 Å. Importantly, the structural changes brought about by ligand binding render MjATPPase competent to interact with MjGATase. The inter-subunit interactions between the residues on helix α1 of MjGATase and α6 of MjATPPase result in allosteric activation of the MjGATase subunit which binds and hydrolyzes Gln. The MjATPPase subunit catalyzes the formation of AMP-XMP intermediate and ammonia tunnelling is facilitated by the reorganization of the MjGATase subunit. Following this, ammonia reacts with the AMP-XMP intermediate generating the products GMP and AMP, which are released along with PPi and Glu, and the MjGATase subunit dissociates. This study lays a strong foundation to probe the reorganization of the MjGATase subunit and ammonia tunnelling in MjGMPS.

## Materials and Methods

### Expression and purification of MjGATase and MjATPPase subunits

The MjATPPase subunit was expressed and purified as described previously.^20^ A glycerol stock of *E. coli* Rosetta (DE3) cells transformed with pST39 vector carrying the gene for the subunit was used to inoculate 5 ml of Luria-Bertani broth supplemented with 100 μg ml^-1^ ampicillin and 34 μg ml^-1^ chloramphenicol and grown overnight. The entire 5 ml was used to inoculate 1 l of Terrific Broth supplemented with antibiotics and grown at 37 °C, till the OD_600_ reached 0.6 to 0.8. Thereafter, the culture was induced with 0.3 mM isopropyl-β-D-thiogalactoside and incubated overnight at 37 °C. The cells were harvested by centrifugation and resuspended in a lysis buffer containing 20 mM Tris-HCl, pH 8.0, 10 % (v/v) glycerol, 2 mM DTT, 0.1 mM PMSF and 0.1 mM EDTA. The cells were lysed by passing through four cycles of French press (Thermo) and the lysate was clarified by centrifugation at 30500 g, 4 °C for 45 min. The supernatant was incubated at 70 °C for 30 min to precipitate the host *E. coli* proteins. The precipitated proteins were pelleted by centrifugation and the nucleic acids in the supernatant were precipitated by the addition of polyethyleneimine to a final concentration of 0.01 % (w/v). The precipitated nucleic acids were pelleted by centrifugation and the supernatant was filtered using 0.45 µm pore size syringe filters. The filtrate was loaded using an ÄKTA Basic HPLC (GE Healthcare) system onto an XK 16/20 column packed with Q-Sepharose HP ion-exchange media (GE Healthcare), which was pre-equilibrated with the lysis buffer. The column was then washed with two column volumes of lysis buffer till the absorbance at 280 nm reduced to a basal value. Bound proteins were eluted using a linear gradient of NaCl from 0 M to 0.3 M over a volume of 450 ml at a flow rate of 3 ml min^-1^. The fractions were examined on 12 % (w/v) SDS-PAGE gels and fractions with pure protein were pooled and dialyzed against buffer containing 20 mM Tris-HCl, pH 7.4, 10 % (v/v) glycerol and 2 mM DTT. The protein was concentrated using centrifugal concentrators, flash-frozen in liquid nitrogen and stored in −80 °C freezer. The concentration of the protein was estimated using Bradford’s assay.^36^ In experiments where the protein was used for crystallization, glycerol was omitted from the dialysis buffer. MjGATase subunit was expressed and purified in an identical manner except that the protein was induced at 37 °C for 6 h and the pH of the lysis buffer was 7.4.^37^

### Expression and purification of MjGMPS fusion protein

The hexa-histidine tagged MjGATase–MjATPPase fused gene was cloned in pET21b expression vector. A BstBI restriction enzyme recognition site was introduced into MjGATase gene by replacing the codon for the penultimate amino acid Phe from TTT to TTC. MjGATase without a stop codon was cloned into pET21b vector using BamHI and SacI restriction sites, which is upstream of the BstBI site. MjATPPase was then cloned downstream of MjGATase using BstBI and XhoI restriction sites. The reading frame was verified by sequencing. *E. coli* Rosetta (DE3) cells were transformed with this construct which expresses the fused protein with an N-terminal hexa-histidine tag. Cells were grown in Terrific Broth supplemented with 100 μg ml^-1^ ampicillin and 34 μg ml^-1^ chloramphenicol at 37 °C and the protein expression was induced at OD_600_ of 0.6 by addition of isopropyl-β-D-thiogalactoside to a final concentration of 0.3 mM. The cells were incubated at 37 °C for 6 h following which they were harvested by centrifugation and the cell pellet was resuspended in lysis buffer containing 20 mM Tris-HCl, pH 7.4, 10 % (v/v) glycerol, 2 mM DTT, and 0.1 mM PMSF. The cells were lysed by passing through four cycles of French press (Thermo) and the lysate was clarified by centrifugation at 30500 g, 4 °C for 45 min. The supernatant was incubated at 70 °C for 30 min to precipitate the host *E. coli* proteins. The precipitated proteins were pelleted by centrifugation and the supernatant was mixed with Ni-NTA agarose beads (Novagen) and tumbled for four hours at 4 °C. This was then transferred to a column and the beads were washed with lysis buffer followed by lysis buffer containing increasing concentrations of imidazole of 10 mM, 25 mM and 50 mM. The protein was eluted with lysis buffer containing 250 mM imidazole and the eluates were run on 12 % (w/v) SDS-PAGE gels. Fractions containing pure protein were pooled, concentrated using centrifugal concentrators and injected into a size-exclusion chromatography column packed with Sephacryl S200 media (GE healthcare) which was pre-equilibrated with the lysis buffer. The flow rate used was 1 ml min^-1^ and the eluates were examined on 12 % (w/v) SDS-PAGE gels. Fractions containing pure protein were pooled, concentrated using centrifugal concentrators, flash-frozen in liquid nitrogen and stored in −80 °C freezer.

### Enzyme assays

#### Gln and NH_4_Cl dependent GMP formation

NH_4_Cl and Gln dependent GMP synthesis by MjATPPase and MjGMPS, respectively, were monitored at 70 °C using a continuous spectrophotometric assay as reported previously.^17^ The reduction in absorbance due to the conversion of XMP to GMP was monitored at 290 nm using a Hitachi U2010 spectrophotometer. A Δε value of 1500 M^-1^ cm^−1^ was used to calculate the concentration of the product formed.^12^ The standard assay mixture consisted of 90 mM HEPES-Na, pH 7.0, 0.2 mM XMP, 3 mM ATP, 20 mM MgCl_2_. The reaction mix was supplemented with either 50 mM NH_4_Cl and 0.57 µM of MjATPPase to measure the NH_4_Cl dependent GMP formation, or 5 mM Gln and 0.57 µM each of MjGATase and MjATPPase to measure Gln dependent GMP formation.

### Measurement of GATase activation

MjGATase activity was monitored using a discontinuous assay where a reaction mixture consisting of 100 mM Tris-HCl, pH 7.4, 0.2 mM XMP, 3 mM ATP, 5 mM Gln, 20 mM MgCl_2_ and 2 μM each of MjGATase and MjATPPase were incubated at 70 °C for 15 minutes. The reaction was quenched by boiling for 5 min, chilling on ice for 5 min followed by centrifugation at 16500 x g. 100 µl of the supernatant was added to a reaction mix containing 100 mM Tris-HCl, pH 7.4, 0.5 mM NAD^+^, 50 mM KCl, 1 mM EDTA and 1.8 units of glutamate dehydrogenase (Sigma-Aldrich). Glutamate dehydrogenase catalyzes the conversion of Glu, that is generated by MjGATase, to α-ketoglutarate with a concomitant reduction of NAD^+^ to NADH. This reaction mix was incubated for an hour at 37 °C and the NADH produced was estimated by measuring the absorbance at 340 nm. The concentration of Glu produced was estimated using a Δε value of 6220 M^-1^ cm^-1^. Reaction where MjGATase was omitted served as a control to measure the background levels of Glu.

### pH and temperature dependence of the activity of MjATPPase, MjGMPS and fused MjGMPS

pH dependence of enzyme activity was monitored by replacing the HEPES-Na buffer in the standard assay with a mixed buffer consisting of 100 mM each of MES, Tris-HCl, and glycine with the pH adjusted at 70 °C. Temperature dependence of activity was monitored under the standard assay conditions. Temperature of the reactions was maintained by water circulated cell holder attached to the Hitachi U-2010 spectrophotometer.

### Estimation of Michaelis-Menten parameters for MjATPPase, MjGMPS and fused MjGMPS

The steady-state kinetic parameters were determined by measuring the initial velocities over a range of concentrations of a substrate (Gln/ATP/XMP), at fixed saturating concentrations of the other two substrates. The pH of the reaction mix was 7.0 and the temperature was 70 °C. The initial velocity data were fit to the Michaelis–Menten equation *v*=(*v*_max_×[S])/(*K*_m_+[S]) by nonlinear regression using the software Prism 5 (GraphPad Software). For all calculations, the concentration of only the MjATPPase subunit was considered to enable comparison of activity across MjATPPase, MjGMPS and fused MjGMPS. The experiments were performed in duplicates and the results are expressed as mean and standard error of the mean (SEM).

### Dependence of MjATPPase and MjGMPS activity on Mg^2+^ concentration

The effect of Mg^2+^ on the activity of MjATPPase subunit and MjGMPS was investigated by monitoring the enzyme activity at varying concentrations of ATP or MgCl_2_ keeping other substrates at saturated concentrations. To investigate whether ATP.Mg^2+^ competes with ATP or the enzyme requires free Mg^2+^ for optimal activity, initial velocity measurements were done at different fixed concentrations of Mg^2+^.ATP. Concentrations of free Mg^2+^ and ATP.Mg^2+^ complex were calculated by using the equations 1 – 2.^38, 39^

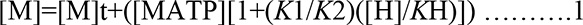

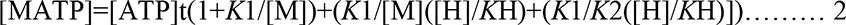

[M]_t_, [ATP]_t_, [MATP], [M] and [H] represent the total Mg^2+^, total ATP, total Mg.ATP^2-^, free Mg^2+^ and free hydrogen ion concentrations, respectively. *K*_1_, *K*_2_ and *K*_H_ represent the dissociation constants for Mg.ATP^2-^, Mg.HATP^-^ and HATP^3-^ complexes, respectively.^16^

### pH dependence of the Michaelis-Menten parameters for Gln and NH_4_Cl dependent GMP formation by MjGMPS

The kinetic parameters *k*_cat_ and *K*_m_ for Gln and NH_4_Cl dependent GMP formation by MjGMPS were determined at various pH using the standard assay conditions at a temperature of 70 °C. A mixed buffer consisting of MES, Tris-HCl and glycine at a concentration of 100 mM each was used. The concentrations of either NH_4_Cl or Gln were varied at a fixed saturating concentration of XMP, ATP and MgCl_2_. The initial velocity data at each pH was fit to the Michaelis-Menten equation by non-linear regression using Prism 5 (GraphPad Software) to derive the kinetic parameters.

### Crystallization and structure solution of MjATPPase

MjATPPase was dialyzed into a buffer containing 20 mM Tris-HCl, pH 7.4 and 2 mM DTT and mixed with a stock solution of the ligands namely PPi.Mg^2+^, ATP.Mg^2+^, AMP-PNP.Mg^2+^, XMP or a combination of these. The pH of the stocks was neutralized using Tris base and the ligands were added in small increments to prevent the precipitation of protein. Following this, the protein was centrifuged at 9600 g, 4 °C for 10 min and the supernatant was filtered using a 0.2 μm pore size syringe filter.^40^ Crystallization screens were set-up using the microbatch under-oil method at 21 °C.^41^ To a 72 well Terasaki plate (Greiner Bio), 8 ml of 1:1 paraffin oil: silicone oil (Hampton research) was poured before adding 2 μl of protein and 2 μl of the crystallization condition to each well. A total of seven commercial crystallization screens comprising 624 unique conditions were screened for either MjATPPase alone or the protein in combination with the ligands. For setting up grid screens, malonic acid/ imidazole/ boric acid (MIB) buffer at various pH were prepared as described before.^42^

A crystal that had grown by mixing 2 μl of 18 mg ml^-1^ MjATPPase supplemented with 1 mM XMP, 1 mM PPi and 20 mM MgCl_2_ with an equal volume of a solution containing 20 % (w/v) PEG 1500 and 0.1 M malonic acid/ imidazole/ boric acid buffer, pH 5.5 was cryoprotected by the addition of small increments of a cryoprotectant solution containing 7.8 mM XMP, 40 % (w/v) PEG 1500 and 0.1 M MIB buffer, pH 5.5. The crystal was flash frozen using a nitrogen cryo-stream at 100 K and a diffraction data set was collected using a Bruker Microstar Ultra II copper rotating anode X-ray generator equipped with a MarResearch MAR345 image plate detector. The data consisted of 190 images, each collected with a 5 min exposure at an increment of 1 degree in φ.

The structure was solved using the CCP4 (Collaborative Computational Project No. 4) suite of programs unless mentioned otherwise. The intensity data were indexed and integrated using iMOSFLM.^43, 44^ The data were scaled using AIMLESS and the intensities were converted to structure factor amplitudes using Ctruncate.^45–47^ The program Matthews_coef was used to determine the number of molecules in the asymmetric unit. Phase information was obtained using PhaserMR^48^ using the crystal structure of PhATPPase (PDB ID: 2DPL, sequence identity of 65.9%) as the search model. The search model was prepared using the CHAINSAW^49^ by pruning the non-conserved residues to the last common atom. The resulting model was then subjected to a round of automated model building using the Phenix program Autobuild.^50^ This model was then subjected to many rounds of manual refinement using the program COOT.^51^ After each round of manual refinement, the resulting model was subjected to restrained refinement using Refmac5.^52^ The Rfactor and Rfree were monitored after each cycle of refinement. After reaching a point where the residuals no longer converged, the ligand was placed into the electron density. This was followed by placing waters into spherical electron density of 1.3 σ in a 2Fo-Fc map. The final model was validated using RAMPAGE^53^ and PDB validation servers.^54, 55^

### Trapping the MjGATase-MjATPPase complex by covalent cross-linking

The MjGATase and MjATPPase subunits were first buffer exchanged into 20 mM HEPES-Na, pH 7.0 and 2 mM DTT. A 50 µl reaction mix consisting of 20 mM HEPES-Na, pH 7.0, 50 µM each of MjGATase and MjATPPase, 0.2 mM XMP, 3 mM ATP and 20 mM MgCl_2_ was incubated at 70 °C for 2 min to promote the formation of complex. Following this, the amine reactive cross-linker bis(sulfosuccinimidyl)suberate (BS3) was added to a final concentration of 5 mM from a stock solution prepared in 20 mM HEPES-Na, pH 7.0. The contents of the tube were quickly mixed and incubated at 50 °C for 5 min. The cross-linking reaction was quenched by adding 4X SDS-PAGE loading dye. Experimental controls included a reaction where both the subunits were cross-linked in absence of substrates and reactions containing only one of the subunits that were cross-linked in the presence and absence of substrates. The cross-linked products were separated using 12 % (w/v) SDS-PAGE and the bands were visualized by staining with Coomassie Brilliant Blue. The molecular masses of the protein bands were assessed by comparing with the mass of the bands in the lane containing the standards (Abcam). The experiment was performed in triplicate and the bands corresponding to the heterodimer and heterotetramer were visualized in all the three experiments.

### Sample preparation and LC-MS/MS data acquisition

The SDS-PAGE bands corresponding to the heterodimer (56 kDa) and heterotetramer (112 kDa) from duplicate experiments were excised and the proteins trypsinised in-gel following a published protocol.^56^ Briefly, the gels were minced into small pieces (∼ 1 mm^3^), destained by incubating in 1:1 (v/v) 100 mM ammonium bicarbonate: acetonitrile (15 min, two washes), dehydrated in acetonitrile and dried in a SpeedVac (Thermo Fisher Scientific). The gel pieces were then treated with 10 mM DTT in 100 mM ammonium bicarbonate to reduce the cystines, washed with acetonitrile, and the cysteines were alkylated using 55 mM iodoacetamide in 100 mM ammonium bicarbonate. The gels were dehydrated with acetonitrile and rehydrated with 10 mM ammonium bicarbonate containing 13 ng µl^-1^ trypsin (Promega). The protease digestion was carried out for 12 h at 37 °C following which the peptides were extracted from the gels using 1:2 (v/v) 5% formic acid: acetonitrile.

The peptides were dried using a SpeedVac, re-dissolved in 0.1% formic acid and injected into an Acclaim PepMap 100 trap column (C18, 2 cm, 75 µm inner diameter, 3 µm particle size, 100 Å pore size) using an Easy-nLC 1200 chromatography system (Thermo Fisher Scientific). The mobile phase consisted of buffer A (0.1% (v/v) formic acid) and buffer B (80% (v/v) acetonitrile and 0.1% (v/v) formic acid). The peptides were separated using an Easy-Spray column (C18, 25 cm, 75 μm inner diameter, 2 μm particle size, 100 Å pore size) maintained at 40 °C and a flow rate of 300 nl min^-1^. The peptides were eluted using a gradient, with the concentration of buffer B ramped up from 5% to 35% in 45 min and held constant at 35% for 3 min. The concentration was increased from 35% to 95% in 2 min and held constant at 95% for 10 min to wash the column. The eluted peptides were infused into a Q Exactive HF mass spectrometer (Thermo Fisher Scientific) operating in positive ion mode. The orbitrap detector was calibrated using LTQ Velos ESI positive ion calibration mixture (Thermo Fisher Scientific). The mass spectrometric data were acquired using the data dependent acquisition mode with the ten most abundant precursor ions selected for fragmentation using a higher-energy C-trap dissociation (HCD) cell. To maximize the identification of cross-linked peptides, upto six LC-MS/MS data sets were acquired for each sample by varying the MS1 and MS2 scanning parameters, as well as the HCD fragmentation settings (Table S10).

### Identification of cross-linked residues

The cross-linked residues in the biological replicates of the heterodimer and the heterotetramer were identified using the program pLink2 version 2.3.9^57^ against a database consisting of the MjGATase (Uniprot ID: Q58970) and MjATPPase (Uniprot ID: Q58531) protein sequences. The six RAW data files (4 files for one among the two heterodimer samples) for each in-gel digest were together input into pLink2 and analysed collectively. The settings used were, cross-linker-BS3 (reactivity only to N-terminus and Lys residues), proteolytic enzyme-trypsin, number of missed cleavages-3, peptide mass range 400 to 6000 Da, peptide length 4 to 60 residues, precursor mass tolerance 10 ppm, fragment mass tolerance of 20 ppm, fixed modifications-carbamidomethylation of Cys, variable modifications-deamidation of Asn and Gln, oxidation of Met, modification of the protein amino terminus and residues Lys, Ser, Thr, Tyr by BS3_OH (156.078644 Da), modification of residues Lys, Ser, Thr, Tyr by BS3_NH2 (155.09463 Da) and modification of residues Lys, Ser, Thr, Tyr by BS3_Tris (259.141973 Da). An additional filter tolerance of 10 ppm and false discovery rate of 5 % at the peptide spectrum matches level was applied. The MjGATase intralinks, MjATPPase intralinks and MjATPPase-MjGATase inter-subunit cross-links that were reproducibly identified in the two biological replicate experiments involving the heterodimer and heterotetramer were listed (Supplementary file 2). The tandem mass spectra of the MjGATase-MjATPPase inter-subunit cross-links were visualized using the program pLabel.^58^ The 21×2 tandem mass spectra of the inter-subunit cross-links in the heterodimer samples and 13×2 spectra from the heterotetramer were manually inspected for the presence of fragment ions that permit unambiguous identification of cross-linked residues (Supplementary files 3A and 3B). The distribution of the cross-links on the MjGATase and MjATPPase protein sequences was visualized using xiVIEW web server.^59^ The cross-links were visualized in the MjGATase and MjATPPase/XMP crystal structures and the Euclidean distances between the Cα atoms of the cross-linked residues were measured using the PyXlinkViewer plugin^60^ for PyMOL.^61^

### Integrative modelling of the MjGMPS complex

The 24 MjGATase-MjATPPase cross-links identified by XL-MS were used as restraints to visualize the accessible interaction space of the two subunits using DisVis web server.^31^ The crystal structures of the MjATPPase/XMP dimer and MjGATase, which were stripped of water molecules, were used as the fixed and scanning chains, respectively. The chain B of MjATPPase/XMP dimer was renamed and the residues renumbered to make it amenable for analysis. The maximum distances between the Cα atoms of cross-linked residues were set to 20.4, 25.3 and 30 Å for the N-terminus‒N-terminus, Lys‒N-terminus and Lys‒Lys cross-links, respectively.^30^ DisVis was executed with the default parameters of the complete scanning mode. The DisVis input as well as the output files are have been deposited to the ProteomeXchange Consortium *via* the PRIDE^62^ partner repository with the dataset identifier PXD031519. The interaction space was visualized using PyMOL and ChimeraX.^63^

The HADDOCK 2.4 webserver was used for the integrative modelling of the MjGATase-MjATPPase complex.^32^ The dimer of MjATPPase/XMP and MjGATase structures were prepared as described above for DisVis analysis. Working in the expert access level of HADDOCK, a TBL file containing distance restraints was uploaded specifying the 24 cross-links as unambiguous restraints. The distance allowances between the Cα atoms of the cross-linked residues were as described for DisVis analysis. Note that none of the Cα atoms of the cross-linked residues were disordered in the two structures. All the parameters were set to default except that the centre-of-mass restraints was turned on. The analysis was repeated using the 12 cross-links of Subset 1 and Subset 2. The structural models with the best HADDOCK scores were then analysed using PyMOL. The input TBL and other files as well as the output files are have been deposited to the ProteomeXchange Consortium *via* the PRIDE^62^ partner repository with the dataset identifier PXD031519.

### Analytical size-exclusion chromatography

Analytical size-exclusion chromatography was performed using an HR 10/30 column packed with Superdex 200 matrix (GE Healthcare) attached to an ÄKTA basic HPLC system equipped with a UV-900 detector (GE Healthcare). The column was equilibrated with 50 mM Tris HCl, pH 7.4, 100 mM KCl and calibrated with β-amylase (200 kDa), alcohol dehydrogenase (150 kDa), bovine serum albumin (66 kDa), carbonic anhydrase (29 kDa) and cytochrome C (12.4 kDa) as molecular weight markers (Sigma Aldrich). A sample volume of 100 µl was injected into the column individually and eluted at a flow rate of 0.5 ml min^-1^. The eluted proteins were detected by monitoring absorbance at 220 nm.

### Multiple sequence alignment

Protein sequences were retrieved from the GenBank and UniProt sequence databases. The sequence alignment was performed using the MAFFT webserver.^64^ The alignment picture was generated using the tool ESPript.^65^

## Abbreviations

GMP: guanosine 5’-monophosphate
XMP: xanthosine 5’-monophosphate
Gln: glutamine
GMPS: GMP synthetase
GATase: glutamine amidotransferase
ATPPase: ATP pyrophosphatase
PfGMPS: *Plasmodium falciparum* GMP synthetase
MjGMPS: *Methanocaldococcus jannaschii* GMP synthetase
MjGATase: GATase subunit of MjGMPS
MjATPPase: ATPPase subunit of MjGMPS
PhGMPS: *Pyrococcus horikoshii* GMP synthetase
XL-MS: cross-linking mass spectrometry
BS3: bis(sulfosuccinimidyl)suberate
PPi: pyrophosphate
HCD: higher-energy C-trap dissociation

## Accession numbers

Coordinates and structure factors of the crystal structure of MjATPPase have been deposited in the Protein Data Bank with the accession code 6JP9. The mass spectrometry data and analysis to identify the cross-linked residues, input as well as output files pertaining to DisVis and HADDOCK analysis have been deposited to the ProteomeXchange Consortium *via* the PRIDE^62^ partner repository with the dataset identifier PXD031519. The data can be accessed using the below credentials:

Username

Password: kaAuU02S

## Author contributions

SS: Crystal structure of MjATPPase, cross-linking mass spectrometry and data analysis, integrative modelling using Dis-Vis and HADDOCK, interpretation and analysis of crystal structure and XL-MS guided modelling, writing the manuscript; SK: cloning and expression of MjATPPase, MjGATase and fused MjGMPS, all enzymatic assays and size-exclusion chromatography of MjATPPase, MjGMPS and fused MjGMPS, interpretation of kinetic data, initial crystallization and cross-linking experiments, reading of manuscript; AB: Crystal structure of MjATPPase, reading of manuscript; DPVSS : crystallization and initial structure solution, HB: conceptualization, interpretation and analysis of all data, funding and writing the manuscript.

## Acknowledgements

The diffraction data was collected at the X-ray facility at Molecular Biophysics Unit, Indian Institute of Science, Bengaluru, India. We thank Prof. Udupi A. Ramagopal from the Poornaprajna Institute of Scientific Research, Bangalore, India; Prof. B. Gopal, and Dr. K. V. Abhinav from the Molecular Biophysics Unit, Indian Institute of Science, Bengaluru, India for help in solving the crystal structure. The mass spectrometry data was collected at the mass spectrometry facility at the Molecular Biology and Genetics Unit, Jawaharlal Nehru Centre of Advanced Scientific Research (JNCASR), Bengaluru, India. For this research HB acknowledges funding by the Department of Biotechnology, Ministry of Science and Technology, Government of India Grants BT/PR11294/BRB/10/1291/2014, BT/PR13760/COE/34/42/2015, and BT/INF/22/SP27679/ 2018; Science and Engineering Research Board (SERB) CRG/2019/004150/IBS, Department of Science and Technology, Government of India Grant EMR/2014/001276; JC Bose Fellowship, SERB and institutional funding from JNCASR. SS was supported by UGC junior and senior research fellowship and a postdoctoral fellowship by JNCASR. SK thanks the Council of Scientific & Industrial Research (CSIR), Government of India for junior and senior research fellowships and JNCASR for a postdoctoral fellowship. AB was supported by CSIR for junior and senior research fellowships, and from JC Bose fellowship awarded to HB.

## Footnotes

^#^These authors contributed equally.

## References

1. Wheeldon, I., Minteer, S.D., Banta, S., Barton, S.C., Atanassov, P., and Sigman, M. (2016). Substrate channelling as an approach to cascade reactions. Nat. Chem. 8, 299–309.

2. Walsh, C.T., and Moore, B.S. (2019). Enzymatic cascade reactions in biosynthesis. Angew. Chem. Int. Ed. Engl. 58, 6846–6879.

3. Raushel, F.M., Thoden, J.B., and Holden, H.M. (2003). Enzymes with molecular tunnels. Acc. Chem. Res. 36, 539–548.

4. Weeks, A., Lund, L., and Raushel, F.M. (2006). Tunneling of intermediates in enzyme-catalyzed reactions. Curr. Opin. Chem. Biol. 10, 465–472.

5. Zalkin, H. (1993). The amidotransferases. Adv. Enzymol. Relat. Areas. Mol. Biol. 66, 203–309.

6. Massière, F., and Badet-Denisot, M.A. (1998). The mechanism of glutamine-dependent amidotransferases. CMLS, Cell. Mol. Life Sci. 54, 205–222.

7. Raushel, F.M., Thoden, J.B., and Holden, H.M. (1999). The amidotransferase family of enzymes: molecular machines for the production and delivery of ammonia. Biochemistry 38, 7891–7899.

8. Zalkin, H., and Smith, J.L. (1998). Enzymes utilizing glutamine as an amide donor. Adv. Enzymol. Relat. Areas Mol. Biol. 72, 87–144.

9. List, F., Vega, M.C., Razeto, A., Häger, M.C., Sterner, R., and Wilmanns, M. (2012). Catalysis uncoupling in a glutamine amidotransferase bienzyme by unblocking the glutaminase active site. Chem. Biol. 19, 1589–1599.

10. Mouilleron, S., and Golinelli-Pimpaneau, B. (2007). Conformational changes in ammonia-channeling glutamine amidotransferases. Curr. Opin. Struct. Biol. 17, 653–664.

11. Mouilleron, S., Badet-Denisot, M.-A., Badet, B., and Golinelli-Pimpaneau, B. (2011). Dynamics of glucosamine-6-phosphate synthase catalysis. Arch. Biochem. Biophys. 505, 1–12.

12. Moyed, H.S., and Magasanik, B. (1957). Enzymes essential for the biosynthesis of nucleic acid guanine; xanthosine 5’-phosphate aminase of Aerobacter aerogenes. J. Biol. Chem. 226, 351–363.

13. von der Saal, W., Crysler, C.S., and Villafranca, J.J. (1985). Positional isotope exchange and kinetic experiments with Escherichia coli guanosine-5’-monophosphate synthetase. Biochemistry 24, 5343– 5350.

14. Maruoka, S., Horita, S., Lee, W.C., Nagata, K., and Tanokura, M. (2010). Crystal structure of the ATPPase subunit and its substrate-dependent association with the GATase subunit: a novel regulatory mechanism for a two-subunit-type GMP synthetase from Pyrococcus horikoshii OT3. J. Mol. Biol. 395, 417–429.

15. Zalkin, H., and Truitt, C.D. (1977). Characterization of the glutamine site of Escherichia coli guanosine 5’-monophosphate synthetase. J. Biol. Chem. 252, 5431–5436.

16. Nakamura, J., and Lou, L. (1995). Biochemical characterization of human GMP synthetase. J. Biol. Chem. 270, 7347–7353.

17. Bhat, J.Y., Shastri, B.G., and Balaram, H. (2008). Kinetic and biochemical characterization of Plasmodium falciparum GMP synthetase. Biochem. J. 409, 263–273.

18. Franco, T.M.A., Rostirolla, D.C., Ducati, R.G., Lorenzini, D.M., Basso, L.A., and Santos, D.S. (2012). Biochemical characterization of recombinant guaA-encoded guanosine monophosphate synthetase (EC 6.3.5.2) from Mycobacterium tuberculosis H37Rv strain. Arch. Biochem. Biophys. 517, 1–11.

19. Bhat, J.Y., Venkatachala, R., Singh, K., Gupta, K., Sarma, S.P., and Balaram, H. (2011). Ammonia channeling in Plasmodium falciparum GMP synthetase: investigation by NMR spectroscopy and biochemical assays. Biochemistry 50, 3346–3356.

20. Ali, R., Kumar, S., Balaram, H., and Sarma, S.P. (2013). Solution nuclear magnetic resonance structure of the GATase subunit and structural basis of the interaction between GATase and ATPPase subunits in a two-subunit-type GMPS from Methanocaldococcus jannaschii. Biochemistry 52, 4308– 4323.

21. Thoden, J.B., Raushel, F.M., Benning, M.M., Rayment, I., and Holden, H.M. (1999). The structure of carbamoyl phosphate synthetase determined to 2.1 A resolution. Acta Crystallogr. Sect. D, Biol. Crystallogr. 55, 8–24.

22. Morar, M., Hoskins, A.A., Stubbe, J., and Ealick, S.E. (2008). Formylglycinamide ribonucleotide amidotransferase from Thermotoga maritima: structural insights into complex formation. Biochemistry 47, 7816–7830.

23. Smith, A.M., Brown, W.C., Harms, E., and Smith, J.L. (2015). Crystal structures capture three states in the catalytic cycle of a pyridoxal phosphate (PLP) synthase. J. Biol. Chem. 290, 5226–5239.

24. Spraggon, G., Kim, C., Nguyen-Huu, X., Yee, M.C., Yanofsky, C., and Mills, S.E. (2001). The structures of anthranilate synthase of Serratia marcescens crystallized in the presence of (i) its substrates, chorismate and glutamine, and a product, glutamate, and (ii) its end-product inhibitor, L-tryptophan. Proc. Natl. Acad. Sci. USA 98, 6021–6026.

25. Semmelmann, F., Hupfeld, E., Heizinger, L., Merkl, R., and Sterner, R. (2019). A Fold-Independent Interface Residue Is Crucial for Complex Formation and Allosteric Signaling in Class I Glutamine Amidotransferases. Biochemistry 58, 2584–2588.

26. Abbott, J.L., Newell, J.M., Lightcap, C.M., Olanich, M.E., Loughlin, D.T., Weller, M.A., Lam, G., Pollack, S., and Patton, W.A. (2006). The effects of removing the GAT domain from E. coli GMP synthetase. Protein J. 25, 483–491.

27. Dongre, A.V., Das, S., Bellur, A., Kumar, S., Chandrashekarmath, A., Karmakar, T., Balaram, P., Balasubramanian, S., and Balaram, H. (2021). Structural basis for the hyperthermostability of an archaeal enzyme induced by succinimide formation. Biophys. J. 120, 3732–3746.

28. Shivakumaraswamy, S., Pandey, N., Ballut, L., Violot, S., Aghajari, N., and Balaram, H. (2020). Helices on Interdomain Interface Couple Catalysis in the ATPPase Domain with Allostery in Plasmodium falciparum GMP Synthetase. Chembiochem 21, 2805–2817.

29. Welin, M., Lehtiö, L., Johansson, A., Flodin, S., Nyman, T., Trésaugues, L., Hammarström, M., Gräslund, S., and Nordlund, P. (2013). Substrate specificity and oligomerization of human GMP synthetase. J. Mol. Biol. 425, 4323–4333.

30. Merkley, E.D., Rysavy, S., Kahraman, A., Hafen, R.P., Daggett, V., and Adkins, J.N. (2014). Distance restraints from crosslinking mass spectrometry: mining a molecular dynamics simulation database to evaluate lysine-lysine distances. Protein Sci. 23, 747–759.

31. van Zundert, G.C.P., Trellet, M., Schaarschmidt, J., Kurkcuoglu, Z., David, M., Verlato, M., Rosato, A., and Bonvin, A.M.J.J. (2017). The disvis and powerfit web servers: explorative and integrative modeling of biomolecular complexes. J. Mol. Biol. 429, 399–407.

32. van Zundert, G.C.P., Rodrigues, J.P.G.L.M., Trellet, M., Schmitz, C., Kastritis, P.L., Karaca, E., Melquiond, A.S.J., van Dijk, M., de Vries, S.J., and Bonvin, A.M.J.J. (2016). The HADDOCK2.2 Web Server: User-Friendly Integrative Modeling of Biomolecular Complexes. J. Mol. Biol. 428, 720–725.

33. Ballut, L., Violot, S., Shivakumaraswamy, S., Thota, L.P., Sathya, M., Kunala, J., Dijkstra, B.W., Terreux, R., Haser, R., Balaram, H., et al. (2015). Active site coupling in Plasmodium falciparum GMP synthetase is triggered by domain rotation. Nat. Commun. 6, 8930.

34. De Ingeniis, J., Kazanov, M.D., Shatalin, K., Gelfand, M.S., Osterman, A.L., and Sorci, L. (2012). Glutamine versus ammonia utilization in the NAD synthetase family. PLoS One 7, e39115.

35. Tesmer, J.J., Klem, T.J., Deras, M.L., Davisson, V.J., and Smith, J.L. (1996). The crystal structure of GMP synthetase reveals a novel catalytic triad and is a structural paradigm for two enzyme families. Nat. Struct. Biol. 3, 74–86.

36. Bradford, M.M. (1976). A rapid and sensitive method for the quantitation of microgram quantities of protein utilizing the principle of protein-dye binding. Anal. Biochem. 72, 248–254.

37. Kumar, S., Prakash, S., Gupta, K., Dongre, A., Balaram, P., and Balaram, H. (2016). Unexpected functional implication of a stable succinimide in the structural stability of Methanocaldococcus jannaschii glutaminase. Nat. Commun. 7, 12798.

38. Morrison, J.F. (1979). Approaches to kinetic studies on metal-activated enzymes. Meth. Enzymol. 63, 257–294.

39. Robertson, J.G., and Villafranca, J.J. (1993). Characterization of metal ion activation and inhibition of CTP synthetase. Biochemistry 32, 3769–3777.

40. Chayen, N.E. (2009). Rigorous filtration for protein crystallization. J. Appl. Crystallogr. 42, 743– 744.

41. Chayen, N.E., Shaw Stewart, P.D., and Blow, D.M. (1992). Microbatch crystallization under oil — a new technique allowing many small-volume crystallization trials. J. Cryst. Growth 122, 176–180.

42. Newman, J. (2004). Novel buffer systems for macromolecular crystallization. Acta Crystallogr. Sect. D, Biol. Crystallogr. 60, 610–612.

43. Powell, H.R., Battye, T.G.G., Kontogiannis, L., Johnson, O., and Leslie, A.G.W. (2017). Integrating macromolecular X-ray diffraction data with the graphical user interface iMosflm. Nat. Protoc. 12, 1310–1325.

44. Powell, H.R., Johnson, O., and Leslie, A.G.W. (2013). Autoindexing diffraction images with iMosflm. Acta Crystallogr. Sect. D, Biol. Crystallogr. 69, 1195–1203.

45. Evans, P. (2006). Scaling and assessment of data quality. Acta Crystallogr. Sect. D, Biol. Crystallogr. 62, 72–82.

46. Evans, P.R. (2011). An introduction to data reduction: space-group determination, scaling and intensity statistics. Acta Crystallogr. Sect. D, Biol. Crystallogr. 67, 282–292.

47. Evans, P.R., and Murshudov, G.N. (2013). How good are my data and what is the resolution? Acta Crystallogr. Sect. D, Biol. Crystallogr. 69, 1204–1214.

48. McCoy, A.J., Grosse-Kunstleve, R.W., Adams, P.D., Winn, M.D., Storoni, L.C., and Read, R.J. (2007). Phaser crystallographic software. J Appl. Crystallogr. 40, 658–674.

49. Stein, N. (2008). CHAINSAW : a program for mutating pdb files used as templates in molecular replacement. J Appl. Crystallogr. 41, 641–643.

50. Adams, P.D., Afonine, P.V., Bunkóczi, G., Chen, V.B., Davis, I.W., Echols, N., Headd, J.J., Hung, L.-W., Kapral, G.J., Grosse-Kunstleve, R.W., et al. (2010). PHENIX: a comprehensive Python-based system for macromolecular structure solution. Acta Crystallogr. Sect. D, Biol. Crystallogr. 66, 213– 221.

51. Emsley, P., Lohkamp, B., Scott, W.G., and Cowtan, K. (2010). Features and development of Coot. Acta Crystallogr. Sect. D, Biol. Crystallogr. 66, 486–501.

52. Murshudov, G.N., Skubák, P., Lebedev, A.A., Pannu, N.S., Steiner, R.A., Nicholls, R.A., Winn, M.D., Long, F., and Vagin, A.A. (2011). REFMAC5 for the refinement of macromolecular crystal structures. Acta Crystallogr. Sect. D, Biol. Crystallogr. 67, 355–367.

53. Lovell, S.C., Davis, I.W., Arendall, W.B., de Bakker, P.I.W., Word, J.M., Prisant, M.G., Richardson, J.S., and Richardson, D.C. (2003). Structure validation by Calpha geometry: phi,psi and Cbeta deviation. Proteins 50, 437–450.

54. Gore, S., Sanz García, E., Hendrickx, P.M.S., Gutmanas, A., Westbrook, J.D., Yang, H., Feng, Z., Baskaran, K., Berrisford, J.M., Hudson, B.P., et al. (2017). Validation of structures in the protein data bank. Structure 25, 1916–1927.

55. Smart, O.S., Horský, V., Gore, S., Svobodová Vařeková, R., Bendová, V., Kleywegt, G.J., and Velankar, S. (2018). Validation of ligands in macromolecular structures determined by X-ray crystallography. Acta Crystallogr. D Struct. Biol. 74, 228–236.

56. Shevchenko, A., Tomas, H., Havlis, J., Olsen, J.V., and Mann, M. (2006). In-gel digestion for mass spectrometric characterization of proteins and proteomes. Nat. Protoc. 1, 2856–2860.

57. Chen, Z.-L., Meng, J.-M., Cao, Y., Yin, J.-L., Fang, R.-Q., Fan, S.-B., Liu, C., Zeng, W.-F., Ding, Y.-H., Tan, D., et al. (2019). A high-speed search engine pLink 2 with systematic evaluation for proteome-scale identification of cross-linked peptides. Nat. Commun. 10, 3404.

58. Wang, L.-H., Li, D.-Q., Fu, Y., Wang, H.-P., Zhang, J.-F., Yuan, Z.-F., Sun, R.-X., Zeng, R., He, S.-M., and Gao, W. (2007). pFind 2.0: a software package for peptide and protein identification via tandem mass spectrometry. Rapid Commun. Mass Spectrom. 21, 2985–2991.

59. Graham, M.J., Combe, C., Kolbowski, L., and Rappsilber, J. (2019). xiView: A common platform for the downstream analysis of Crosslinking Mass Spectrometry data. BioRxiv. doi: 10.1101/561829.

60. Schiffrin, B., Radford, S.E., Brockwell, D.J., and Calabrese, A.N. (2020). PyXlinkViewer: A flexible tool for visualization of protein chemical crosslinking data within the PyMOL molecular graphics system. Protein Sci. 29, 1851–1857.

61. Schrödinger (2010). The Pymol Molecular Graphics System (Schrödinger).

62. Perez-Riverol, Y., Csordas, A., Bai, J., Bernal-Llinares, M., Hewapathirana, S., Kundu, D.J., Inuganti, A., Griss, J., Mayer, G., Eisenacher, M., et al. (2019). The PRIDE database and related tools and resources in 2019: improving support for quantification data. Nucleic Acids Res. 47, D442–D450.

63. Pettersen, E.F., Goddard, T.D., Huang, C.C., Meng, E.C., Couch, G.S., Croll, T.I., Morris, J.H., and Ferrin, T.E. (2021). UCSF ChimeraX: Structure visualization for researchers, educators, and developers. Protein Sci. 30, 70–82.

64. Katoh, K., Rozewicki, J., and Yamada, K.D. (2017). MAFFT online service: multiple sequence alignment, interactive sequence choice and visualization. Brief. Bioinformatics bbx108.

65. Robert, X., and Gouet, P. (2014). Deciphering key features in protein structures with the new ENDscript server. Nucleic Acids Res. 42, W320–W324.

