## Supplementary file 1 for "Mechanistic insights into the functioning of GMP synthetase: a two-subunit, allosterically regulated, ammonia tunnelling enzyme"

**Title:** Mechanistic insights into the functioning of GMP synthetase: a two-subunit, allosterically regulated, ammonia tunnelling enzyme

**Authors:** Santosh Shivakumaraswamy#, Sanjeev Kumar#, Asutosh Bellur, Satya Dev Polisetty and Hemalatha Balaram*

#These authors contributed equally, *Corresponding author.

**List of figures:**

Figure S1: Crystal structure of XMP bound MjATPPase.

Figure S2: Conformation of the lid-loop connecting the strands β4-β5.

Figure S3: The fit of the lid-loop residues to electron density in MjATPPase/XMP structure.

Figure S4: Conservation of Cys239 and Arg249 in the sequences of ATPPase subunits.

Figure S5: Conformational flexibility of the residues in the loop connecting the antiparallel β-strands β7-β8.

Figure S6: Replicate of MjGATase-MjATPPase cross-linking experiment described in Figure 5.

Figure S7: XL-MS guided modelling of the MjGMPS complex.

Figure S8: Examining the oligomeric state of fused MjGMPS using analytical size-exclusion chromatography.

Figure S9: Interdomain interactions in GMP synthetases.

Figure S10: Repositioning of active sites in PfGMPS due to domain rotation.

**List of tables:**

Table S1: Steady-state kinetic parameters of MjATPPase, MjGMPS and fused MjGMPS.

Table S2: RMSD values for different chains in MjATPPase/XMP structure, the ATPPase subunit of PhGMPS and PfGMPS.

Table S3: List of MjGATase intralinks and structural validation.

Table S4: List of MjATPPase intralinks and structural validation.

Table S5: Exploratory modelling using DisVis.

Table S6: Exploratory modelling using DisVis.

Table S7: Results of HADDOCK modelling.

Table S8: Structural validation of HADDOCK models.

Table S9: Details of the sequences of the ATPPase subunit used in the alignment shown in Figure S4.

Table S10: Data acquisition settings for LC-MS/MS.

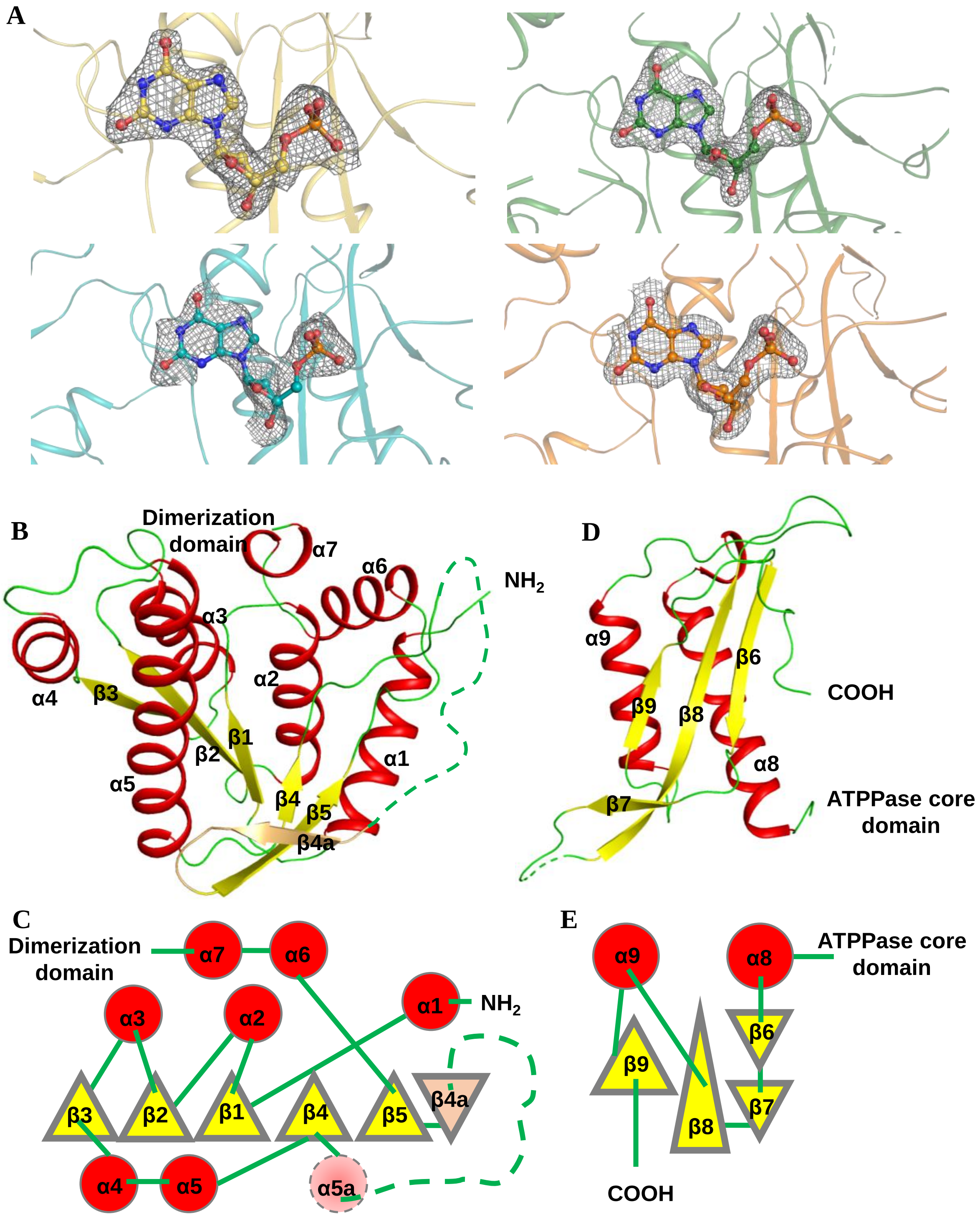

**Figure S1: Crystal structure of XMP bound MjATPPase.** (A) The fit of XMP to electron density in the four chains of the asymmetric unit of MjATPPase/XMP structure. XMP is shown in ball and stick representation and a 2Fo-Fc simulated annealing composite omit map contoured at 1 σ is shown as mesh. Protein backbone is shown in cartoon representation. (B) The structure of the MjATPPase core domain and the corresponding topology diagram (C). (D) The structure of the dimerization domain and its topology diagram (E). The protein backbone is shown in cartoon representation with the β-strands and α-helices colored yellow and red, respectively. The topology diagram is drawn with the β-sheet parallel to the plane of the paper. Triangles, circles and lines represent β-strands, α-helices and loops, respectively. A line drawn to the center of the triangle or circle indicates a loop connecting to the top of the β-strand or α-helix whereas a line drawn to the edge of a triangle or circle indicates a bottom connection. The residues corresponding to α5a are modelled as a helix in the structures of two-domain type GMP synthetases.

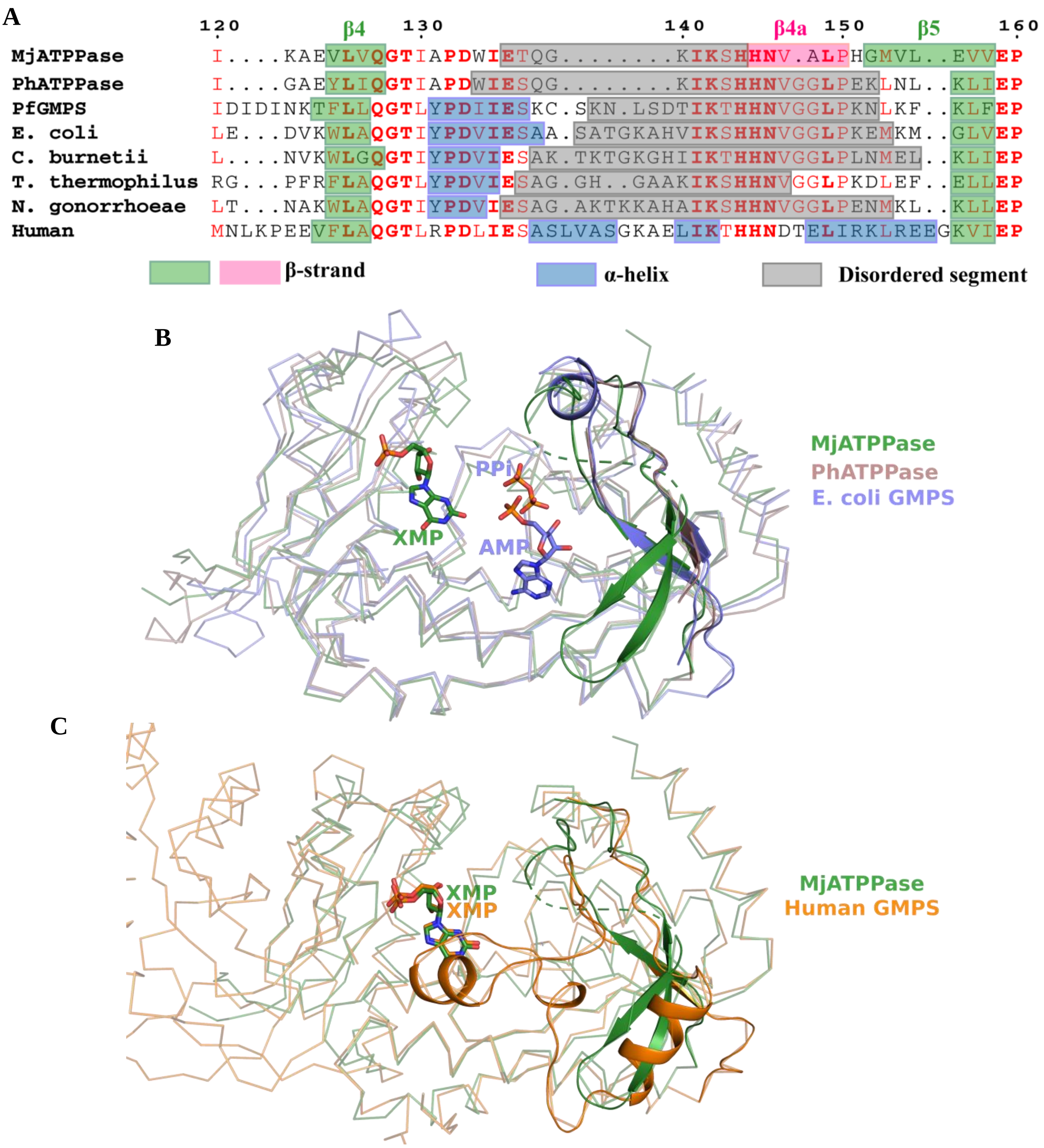

**Figure S2: Conformation of the lid-loop connecting the strands β4-β5.** (A) Sequence alignment of the ATPPase subunits (MjATPPase and PhATPPase) and two-domain type GMP synthetases for which a crystal structure is available. Sequence numbering and the identity of the secondary structural elements on top of the alignment is for MjATPPase. Conserved residues are coloured red and invariant residues are highlighted with a bold font. Secondary structures and disordered regions are indicated using rectangles of different colours. (B) Structural superposition of MjATPPase/XMP on PhATPPase and the ATPPase domain of *E. coli* GMPS. (C) Structural superposition of the ATPPase domain of human GMPS on MjATPPase/XMP. In both B and C, the protein backbone is shown in ribbon representation while the stretch of residues in between and inclusive of β4-β5 is shown is cartoon representation. In MjATPPase/XMP, the disordered stretch spanning residues 136-144 is shown as a dashed line. The ligands AMP, PPi and XMP are shown in stick representation.

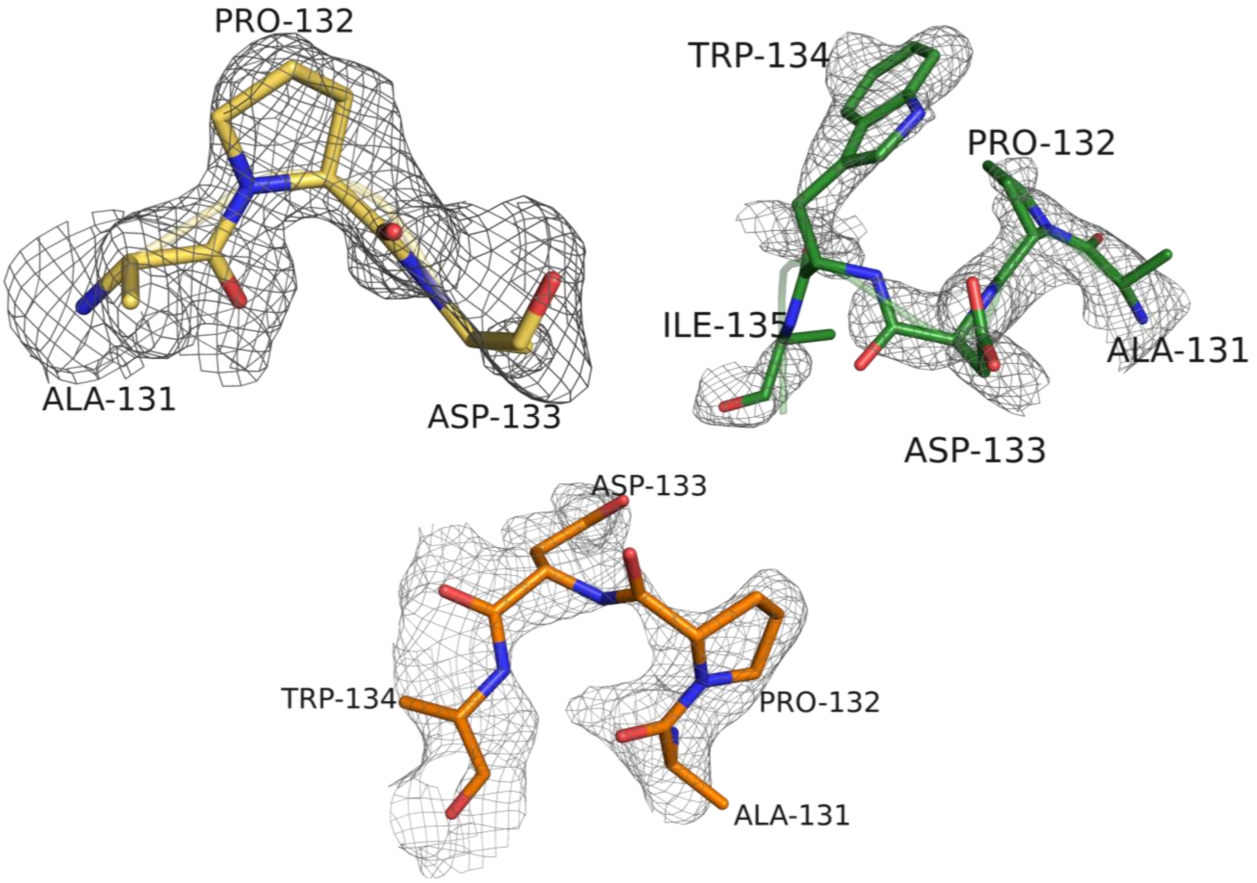

**Figure S3: The fit of the lid-loop residues to electron density in MjATPPase/XMP structure.** The fit of the residues starting from Ala131 in chains A (yellow), B (green) and D (orange) to the electron density. The residues are shown as sticks and a 2Fo-Fc simulated annealing composite omit map contoured at 1 σ is shown as mesh.

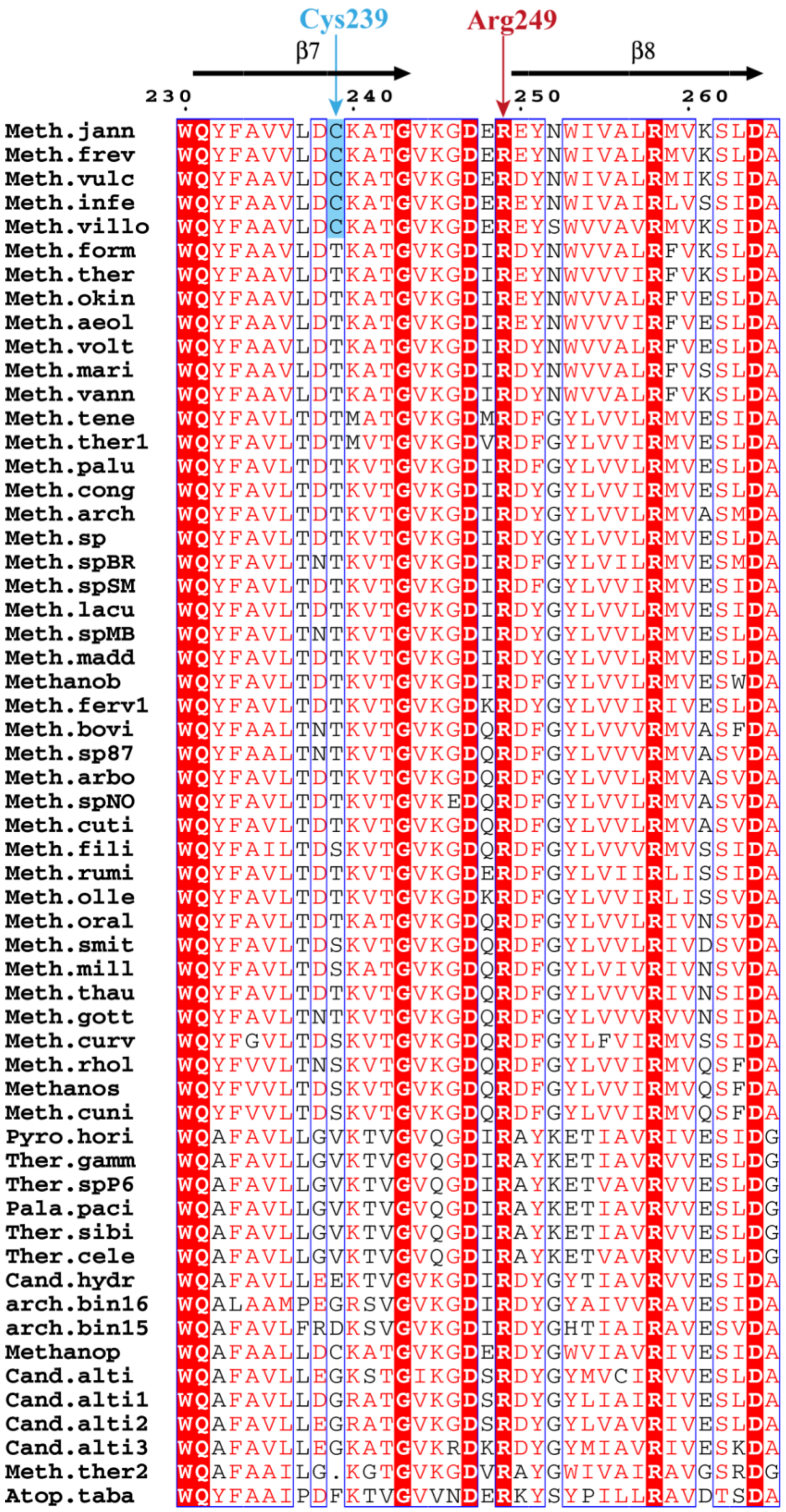

**Figure S4: Conservation of Cys239 and Arg249 in the sequences of ATPPase subunits.** An excerpt of the multiple sequence alignment is shown here. The sequences were retrieved by blasting the MjATPPase sequence using the program BLAST (doi: 10.1016/S0022-2836(05)80360-2). The program CD-Hit (doi: 10.1093/bioinformatics/bts565) was used to cluster the sequences with greater than 90 % identity. The secondary structural elements of chain A in the MjATPPase/XMP structure are shown above the alignment. Numbering is for MjATPPase sequence. Invariant residues are highlighted by red/ white inversion and residues with high sequence similarity are shown in red and boxed in blue. The details of the sequences are in Table S9.

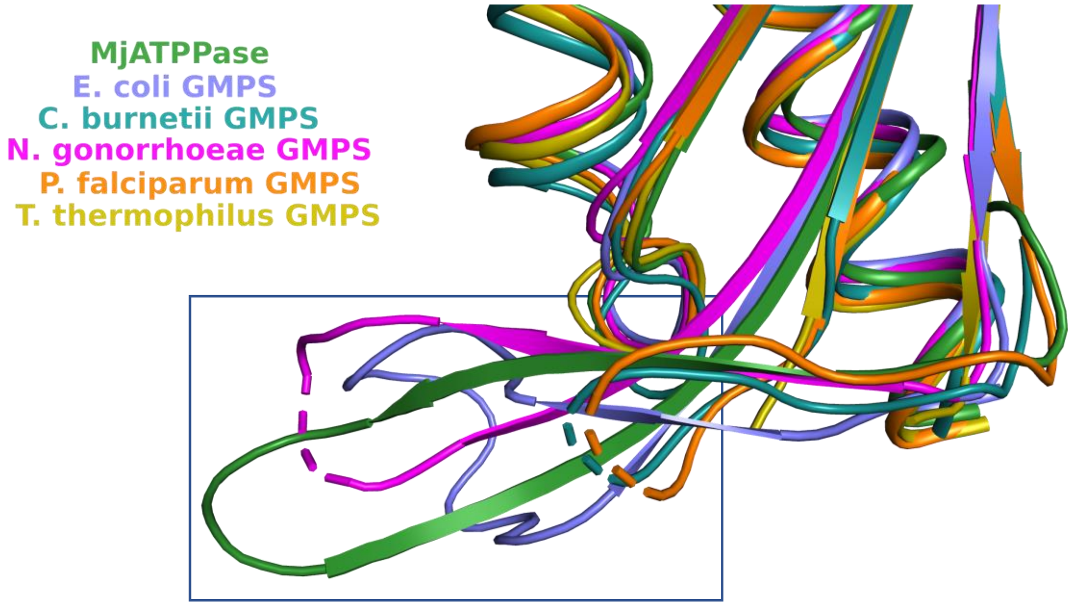

**Figure S5: Conformational flexibility of the residues in the loop connecting the antiparallel β-strands β7-β8.** Structural superposition of the dimerization domains in two-domain type enzymes on the dimerization domain of MjATPPase/XMP structure (chain C). The protein backbone is shown in cartoon representation. The region surrounding the loop connecting the two strands is boxed. The loop residues are modelled only in the structures of MjATPPase and *E.coli* GMPS.

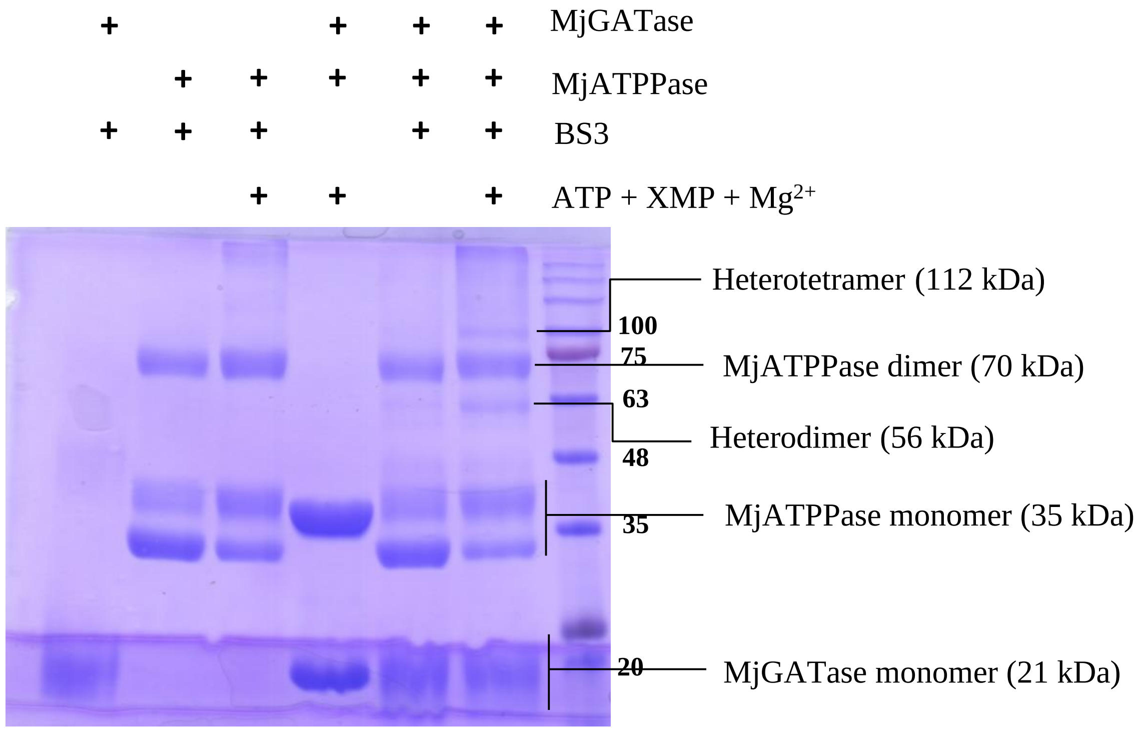

**Figure S6: Replicate of MjGATase-MjATPPase crosslinking experiment described in Figure 5.** 50 µM each of MjGATase and MjATPPase were incubated with 0.2 mM XMP, 2 mM ATP and 20 mM $MgCl_2$ at 70 °C for 2 min before crosslinking with BS3 at 50 °C for 5 min. The reaction was quenched by addition of SDS-PAGE loading dye and the bands were resolved by 12 % (w/v) SDS-PAGE. The bands were visualized by Coomassie Brilliant Blue staining. The protein molecular mass standards are in the last lane with the molecular mass in kDa for specific bands indicated in bold.

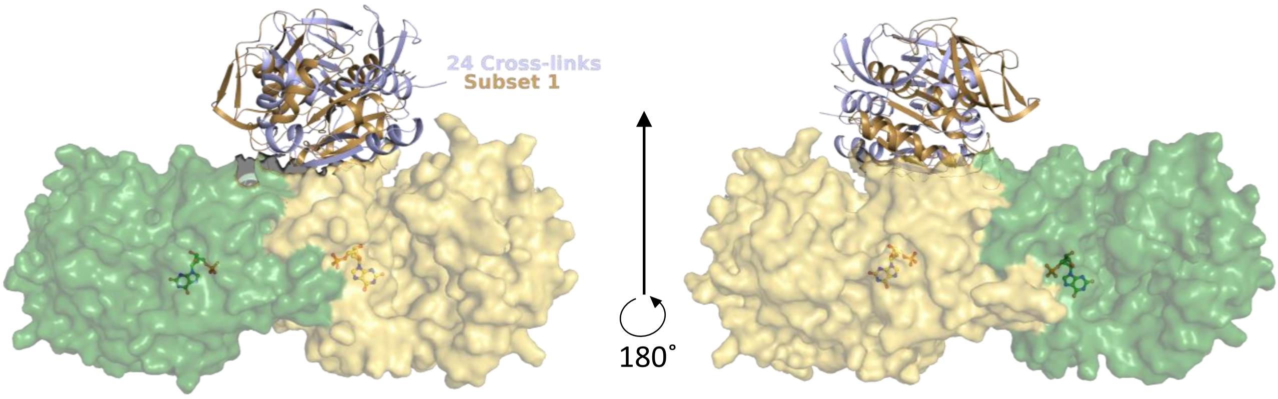

**Figure S7: XL-MS guided modelling of the MjGMPS complex.** HADDOCK modelled structures of the MjGMPS complex generated using the 24 cross-links and the cross-links of Subset 1 are shown. The MjATPPase subunits from the highest scoring models were superposed. The MjATPPase subunits are shown in surface representation and the backbone of the MjGATase subunits is shown as cartoon. XMP is in ball and stick representation.

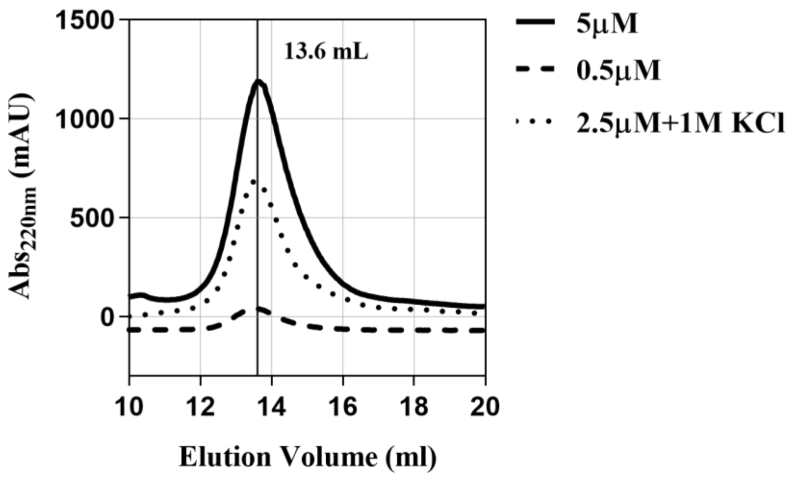

**Figure S8: Examining the oligomeric state of fused MjGMPS using analytical size-exclusion chromatography.** The elution volume of fused MjGMPS did not change upon reducing the protein concentration that was injected nor in presence of 1 M KCl, suggesting that the protein is a strong dimer. Please refer to Figure 8B for more details.

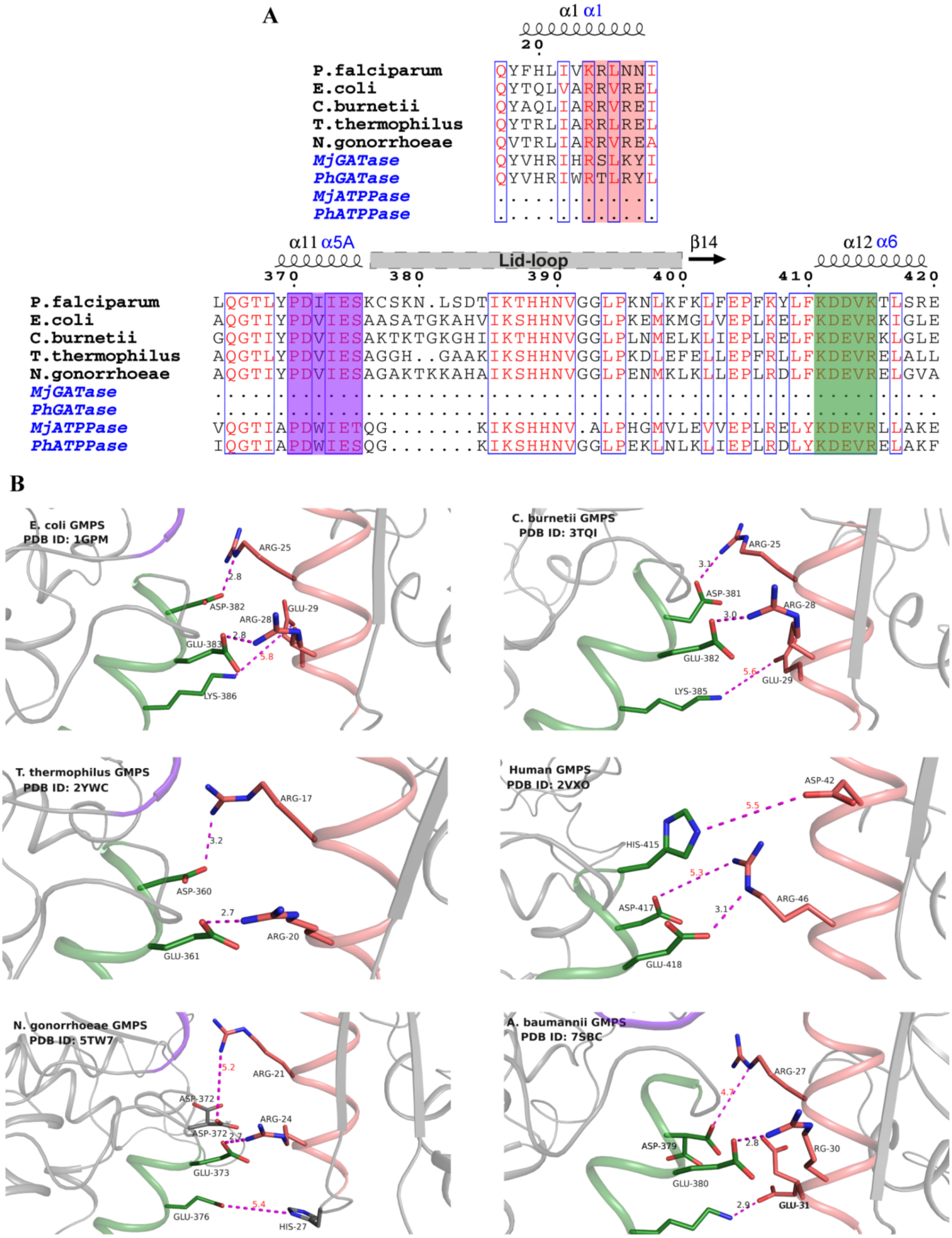

**Figure S9: Interdomain interactions in GMP synthetases.** (A) Multiple sequence alignment of two-domain type and two-subunit type GMP synthetases. The conserved blocks corresponding to interface helices are extracted from the full alignment and shown here. Numbering is for *P. falciparum* GMPS. The conserved motifs corresponding to the three helices are highlighted. (B) Interdomain salt bridges between GATase helix α1 and ATPPase helices α11 and α12. The protein backbone is shown in cartoon representation and the residues are in stick representation. Distances greater than 4 Å are colored red.

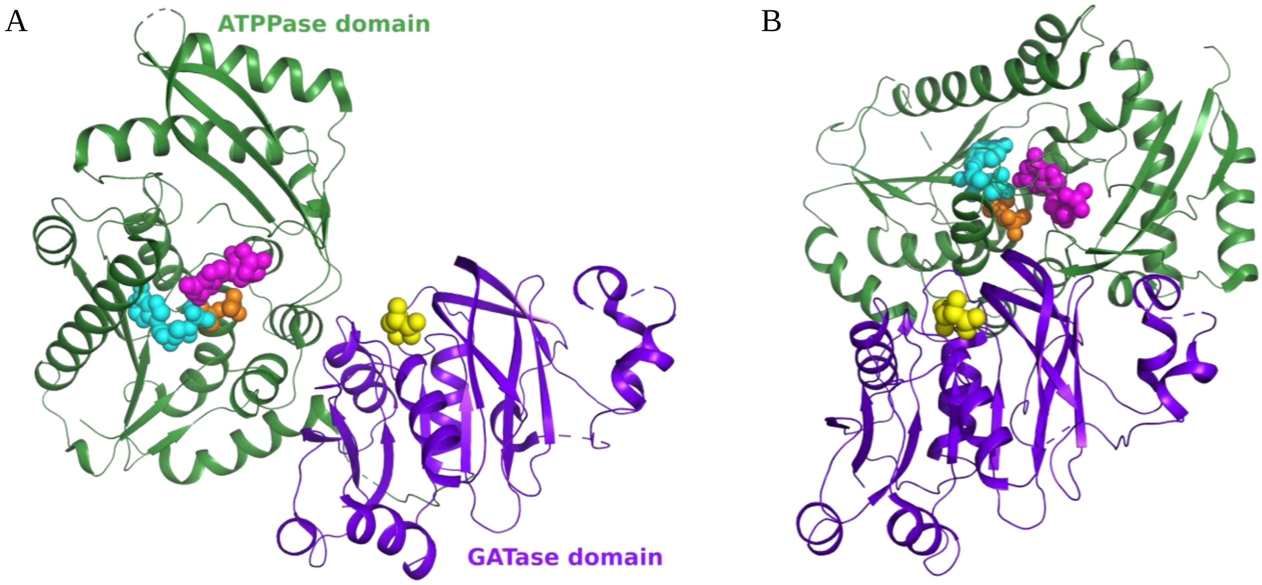

**Figure S10: Repositioning of active sites in PfGMPS due to domain rotation.** (A) Structure of XMP bound PfGMPS (PDB ID: 3UOW) and (B) Gln bound PfGMPS (PDB ID: 4WIO) where the GATase domain is rotated by 85°. Conformation of the GATase domain is the same across A and B, and as a result the ATPPase domain appears to have rotated. Visualizing domain rotation in this manner highlights the changes in the relative orientation of the GATase and ATPPase active sites. The backbone is shown in cartoon representation and the ligands Gln (yellow), XMP (magenta), AMP (cyan) and PPi (orange) are shown as spheres. The ligands provide the approximate location of the active sites. The ligands AMP and PPi are superposed from the structure of *E. coli* GMPS (PDB ID: 1GPM). In (A), Gln is superposed from the PfGMPS/Gln structure.

**Table S1:** Steady-state kinetic parameters of MjATPPase, MjGMPS and fused MjGMPS.

| | Substrate | $K_M$ (μM) | $k_{cat}$ ($s^{-1}$) |
|---|---|---|---|
| **MjATPPase** | ATP.$Mg^{2+}$ | 447 ± 5 | 2.47 ± 0.01 |
| | XMP | 30 ± 2 | 1.89 ± 0.03 |
| | $NH_4Cl$ | 4139 ± 212 | 1.91 ± 0.02 |
| **MjGMPS** | ATP.$Mg^{2+}$ | 452 ± 3 | 2.85 ± 0.02 |
| | XMP | 61 ± 3 | 2.44 ± 0.05 |
| | Glutamine | 520 ± 8 | 1.94 ± 0.02 |
| **Fused MjGMPS** | ATP.$Mg^{2+}$ | 503 ± 6 | 0.995 ± 0.001 |
| | XMP | 44 ± 1 | 1.41 ± 0.03 |
| | Glutamine | 1418 ± 263 | 0.60 ± 0.03 |

All values are mean ± SEM.

**Table S2:** RMSD values for different chains in MjATPPase/XMP structure, the ATPPase subunit of PhGMPS and PfGMPS.

| | **Chain A** | **Chain B** | **Chain C** | **Chain D** | ***P. horikoshii* ATPPase** | | **ATPPase of PfGMPS (PDB ID: 3UOW)** | **ATPPase of PfGMPS (PDB ID: 4WIO)** |
|---|---|---|---|---|---|---|---|---|
| | | | | | **PDB ID: 3A4I** | **PDB ID: 2DPL** | | |
| **Chain A** | | 1.09 (293) | 0.72 (273) | 0.56 (296) | 1.17 (285) | 1.16 (286) | 1.96 (278) | 1.72 (272) |
| **Chain B** | | | 0.96 (270) | 1.34 (295) | | | | |
| **Chain C** | | | | 0.91 (272) | | | | |

The RMSD is in Å. The number of equivalent residues in the structural alignment is in parenthesis. The alignment was performed using the FATCAT web server (doi: 10.1093/nar/gkh430).

**Table S3:** List of MjGATase intralinks and structural validation.

| | Crosslinked residues | Present in heterodimer | Present in heterotetramer | Euclidean distance in Å between the Cα atoms in the apo MjGATase structure | |
|---|---|---|---|---|---|
| | | | | Chain A | Chain B |
| 1 | 107-153 | Yes | Yes | 11.4 | 11.4 |
| 2 | 1-42 | Yes | Yes | 9.2 | 9.2 |
| 3 | 1-45 | Yes | Yes | 5.9 | 5.7 |
| 4 | 45-186 | Yes | Yes | 12.4 | 12.4 |
| 5 | 42-186 | Yes | Yes | 17.6 | 17.6 |
| 6 | 1-186 | Yes | Yes | 12.8 | 12.7 |
| 7 | 113-181 | Yes | Yes | 10.5 | 10.5 |
| 8 | 113-186 | Yes | Yes | 13.9 | 14 |
| 9 | 1-181 | Yes | Yes | 12.2 | 12.1 |
| 10 | 57-126 | Yes | Yes | 16.6 | 16.6 |
| 11 | 107-155 | Yes | Yes | 15.7 | 15.7 |
| 12 | 107-186 | Yes | Yes | 22.3 | 22.3 |
| 13 | 1-27 | Yes | No | 7.2 | 7.4 |
| 14 | 27-45 | Yes | No | 11.5 | 11.4 |
| 15 | 1-20 | No | Yes | 10.6 | 10.7 |
| 16 | 126-131 | No | Yes | 15.6 | 15.7 |

It should be noted that all the distances specified in columns 5 and 6 are within the required 30 Å cut-off. The distances were calculated using the PyMOL plugin PyXlinkViewer.

**Table S4:** List of MjATPPase intralinks and structural validation.

| | Cross-linked residues | Present in heterodimer | Present in heterotetramer | Euclidean distance in Å between the Cα atoms in the MjATPPase/XMP structure | | | |
|---|---|---|---|---|---|---|---|
| | | | | Chain A | Chain B | ChainA_ChainB | ChainB_ChainA |
| 1 | 69-74 | Yes | Yes | 6 | 6 | 77.3 | 77.3 |
| 2 | 85-93 | Yes | Yes | 12.6 | 12.5 | 41.7 | 41.8 |
| 3 | 61-221 | Yes | Yes | 15.2 | 15.3 | 43.7 | 43.8 |
| 4 | 69-221 | Yes | Yes | 21.3 | 20.7 | 55.9 | 55.4 |
| 5 | 1-174 | Yes | Yes | 16.1 | 0 | 70 | 0 |
| 6 | 142-302 | Yes | Yes | 0 | 0 | 0 | 0 |
| 7 | 166-302 | Yes | Yes | 22.1 | 19.5 | 44.2 | 41.5 |
| 8 | 93-103 | Yes | Yes | 13.8 | 13.9 | 35.9 | 36.2 |
| 9 | 166-174 | Yes | Yes | 12.8 | 12.8 | 62.7 | 61.6 |
| 10 | 61-74 | Yes | Yes | 20 | 19.9 | 63.9 | 64.3 |
| 11 | 174-221 | Yes | Yes | 19.3 | 17.2 | 52.2 | 49.9 |
| 12 | 221-283 | Yes | No | 16.9 | 17 | 29 | 28.8 |
| 13 | 1-6 | Yes | No | 11.9 | 0 | 81.3 | 0 |
| 14 | 61-69 | Yes | No | 14.6 | 14.4 | 59.5 | 60 |
| 15 | 103-140 | Yes | No | 0 | 0 | 0 | 0 |
| 16 | 101-103 | Yes | No | 5.5 | 5.6 | 39.1 | 39.1 |
| 17 | 23-48 | Yes | No | 5 | 4.9 | 97 | 96.7 |
| 18 | 222-283 | Yes | No | 15.7 | 15.8 | 30.6 | 30.1 |
| 19 | 245-283 | Yes | No | 15.2 | 0 | 34.8 | 0 |
| 20 | 283-302 | Yes | No | 27.1 | 27.1 | 10.9 | 10.9 |
| 21 | 69-85 | No | Yes | 19.8 | 19.5 | 64.7 | 64.2 |
| 22 | 245-302 | No | Yes | 36.9 | 0 | 10.6 | 0 |
| 23 | 221-302 | No | Yes | 23.9 | 23.7 | 27.2 | 27.2 |
| 24 | 1-61 | No | Yes | 36.5 | 0 | 61 | 0 |
| 25 | 48-85 | No | Yes | 27.4 | 27.7 | 74.4 | 73.7 |
| 26 | 166-245 | No | Yes | 48.3 | 0 | 0 | 26.8 |

A box is colored green if the distance is within the 30 Å cut-off or else is colored pink. If a residue involved in a cross-link is disordered in the structure, the distance is specified as 0 Å and the box is colored yellow. The distances were calculated using the PyMOL plugin PyXlinkViewer.

**Table S5:** Exploratory modelling using DisVis. The table lists the average fraction violated and the standard deviation for a crosslink as well as the Z-score.

| | 24 Cross-links | | | | Subset 1 (12 Cross-links) | | | | Subset 2 (12 Cross-links) | | | |
|---|---|---|---|---|---|---|---|---|---|---|---|---|
| | Crosslink | Average Fraction Violated | Standard deviation | Z-score | Crosslink | Average Fraction Violated | Standard deviation | Z-score | Crosslink | Average Fraction Violated | Standard deviation | Z-score |
| 1 | 1-1 | 0.36 | 0.33 | -0.68 | 1-1 | 0.38 | 0.29 | 0.41 | | | | |
| 2 | 1-20 | 0.25 | 0.31 | -1.16 | 1-20 | 0.22 | 0.27 | -0.67 | | | | |
| 3 | 1-27 | 0.27 | 0.3 | -1.05 | 1-27 | 0.25 | 0.25 | -0.43 | | | | |
| 4 | 1-177 | 0.33 | 0.26 | -0.78 | 1-177 | 0.26 | 0.24 | -0.41 | | | | |
| 5 | 1-186 | 0.33 | 0.29 | -0.82 | 1-186 | 0.31 | 0.24 | -0.08 | | | | |
| 6 | 222-1 | 0.58 | 0.26 | 0.32 | | | | | 222-1 | 0.5 | 0.3 | 0.01 |
| 7 | 5-20 | 0.21 | 0.27 | -1.33 | 5-20 | 0.17 | 0.19 | -1.03 | | | | |
| 8 | 6-20 | 0.23 | 0.26 | -1.25 | 6-20 | 0.17 | 0.19 | -1.01 | | | | |
| 9 | 6-27 | 0.29 | 0.23 | -0.97 | 6-27 | 0.22 | 0.17 | -0.68 | | | | |
| 10 | 16-20 | 0.69 | 0.1 | 0.81 | | | | | 16-20 | 0.97 | 0.07 | 1.9 |
| 11 | 174-20 | 0.2 | 0.28 | -1.36 | 174-20 | 0.22 | 0.21 | -0.67 | | | | |
| 12 | 261-20 | 0.3 | 0.31 | -0.91 | 261-20 | 0.42 | 0.25 | 0.71 | | | | |
| 13 | 279-20 | 0.57 | 0.28 | 0.26 | | | | | 279-20 | 0.3 | 0.3 | -0.79 |
| 14 | 279-102 | 0.6 | 0.3 | 0.4 | | | | | 279-102 | 0.3 | 0.28 | -0.77 |
| 15 | 279-126 | 0.62 | 0.31 | 0.46 | | | | | 279-126 | 0.32 | 0.28 | -0.7 |
| 16 | 283-20 | 0.66 | 0.25 | 0.64 | | | | | 283-20 | 0.29 | 0.29 | -0.83 |
| 17 | 283-57 | 0.84 | 0.08 | 1.45 | | | | | 283-57 | 0.45 | 0.27 | -0.2 |
| 18 | 283-102 | 0.66 | 0.28 | 0.64 | | | | | 283-102 | 0.29 | 0.28 | -0.82 |
| 19 | 283-126 | 0.68 | 0.28 | 0.73 | | | | | 283-126 | 0.31 | 0.28 | -0.75 |
| 20 | 283-186 | 0.87 | 0.06 | 1.56 | | | | | 283-186 | 0.48 | 0.24 | -0.05 |
| 21 | 302-20 | 0.47 | 0.31 | -0.17 | 302-20 | 0.59 | 0.23 | 1.82 | | | | |
| 22 | 302-27 | 0.49 | 0.32 | -0.07 | 302-27 | 0.62 | 0.22 | 2.05 | | | | |
| 23 | 302-126 | 0.79 | 0.14 | 1.21 | | | | | 302-126 | 0.79 | 0.27 | 1.2 |
| 24 | 93-155 | 0.98 | 0.04 | 2.07 | | | | | 93-155 | 0.94 | 0.06 | 1.8 |

The cross-linked residues are specified as MjATPPase residue-MjGATase residue. Cross-links that are outliers are flagged in the DisVis output using various shades of pink.

**Table S6:** Exploratory modelling using DisVis. The table lists the number of complexes consistent with a given number of restraints.

| 24 Cross-links | | | Subset 1 (12 Cross-links) | | | Subset 2 (12 Cross-links) | | |
|---|---|---|---|---|---|---|---|---|
| Number of consistent restraints (N) | Number of complexes consistent with N restraints | Fraction of all complexes consistent with N restraints | Number of consistent restraints (N) | Number of complexes consistent with N restraints | Fraction of all complexes consistent with N restraints | Number of consistent restraints (N) | Number of complexes consistent with N restraints | Fraction of all complexes consistent with N restraints |
| 0 | 16702634 | 0.474253 | 0 | 26693379 | 0.75793 | 0 | 20012364 | 0.568229 |
| 1 | 6074400 | 0.172476 | 1 | 2241688 | 0.06365 | 1 | 8441959 | 0.2397 |
| 2 | 3019579 | 0.085738 | 2 | 1471717 | 0.041788 | 2 | 2343545 | 0.066542 |
| 3 | 2504128 | 0.071102 | 3 | 1448362 | 0.041125 | 3 | 1692074 | 0.048045 |
| 4 | 2085009 | 0.059202 | 4 | 766760 | 0.021771 | 4 | 1530987 | 0.043471 |
| 5 | 1500774 | 0.042613 | 5 | 556555 | 0.015803 | 5 | 759534 | 0.021566 |
| 6 | 848653 | 0.024097 | 6 | 444850 | 0.012631 | 6 | 264038 | 0.007497 |
| 7 | 646708 | 0.018363 | 7 | 386160 | 0.010965 | 7 | 121648 | 0.003454 |
| 8 | 505453 | 0.014352 | 8 | 322860 | 0.009167 | 8 | 44858 | 0.001274 |
| 9 | 406757 | 0.011549 | 9 | 454354 | 0.012901 | 9 | 7697 | 0.000219 |
| 10 | 421102 | 0.011957 | 10 | 221163 | 0.00628 | 10 | 106 | 0.000003 |
| 11 | 231195 | 0.006565 | 11 | 146743 | 0.004167 | 11 | 0 | 0 |
| 12 | 151608 | 0.004305 | 12 | 64217 | 0.001823 | 12 | 0 | 0 |
| 13 | 72661 | 0.002063 | | | | | | |
| 14 | 29728 | 0.000844 | | | | | | |
| 15 | 12089 | 0.000343 | | | | | | |
| 16 | 4360 | 0.000124 | | | | | | |
| 17 | 1468 | 0.000042 | | | | | | |
| 18 | 403 | 0.000011 | | | | | | |
| 19 | 93 | 0.000003 | | | | | | |
| 20 | 3 | 0 | | | | | | |
| 21 | 0 | 0 | | | | | | |
| 22 | 0 | 0 | | | | | | |
| 23 | 0 | 0 | | | | | | |
| 24 | 0 | 0 | | | | | | |

**Table S7:** Results of HADDOCK modelling.

| | 24 cross-links | Subset 1 | Subset 2 |
|---|---|---|---|
| **Number of clusters** | 1 | 7 | 4 |
| **Top Cluster based on HADDOCK score** | Cluster 1 | Cluster 1 | Cluster 2 |
| **HADDOCK score** | -60.2 +/- 0.8 | -124.3 +/- 2.5 | -82.4 +/- 2.2 |
| **Cluster size** | 200 | 140 | 35 |
| **RMSD from the overall lowest-energy structure** | 0.5 +/- 0.3 | 0.4 +/- 0.2 | 0.3 +/- 0.2 |
| **Van der Waals energy** | -44.2 +/- 2.4 | -36.4 +/- 4.3 | -46.6 +/- 0.9 |
| **Electrostatic energy** | -305.6 +/- 33.5 | -435.7 +/- 26.1 | -379.2 +/- 24.5 |
| **Desolvation energy** | -1.2 +/- 3.3 | -0.9 +/- 0.6 | 4.9 +/- 2.1 |
| **Restraints violation energy** | 463.6 +/- 10.9 | 0.3 +/- 0.1 | 351.9 +/- 2.3 |
| **Buried Surface Area** | 1774.7 +/- 61.0 | 1654.4 +/- 29.0 | 1617.6 +/- 65.6 |
| **Z-Score** | 0 | -1.4 | -1.7 |

Table S8: Structural validation of HADDOCK models.

| | | Euclidean distance in Å between the Cα atoms in the MjATPPase/XMP structure | | |
|---|---|---|---|---|
| Cross-link MjATPPase-MjGATase | Distance threshold (Å) | 24 Cross-links | Subset 1 | Subset 2 |
| 1-1 | 20.4 | 15.6 | 19.5 | |
| 1-20 | 25.3 | 22 | 10.1 | |
| 1-27 | 25.3 | 18.6 | 20.6 | |
| 1-177 | 25.3 | 26.7 | 14.6 | |
| 1-186 | 25.3 | 22.2 | 16.3 | |
| 5-20 | 30 | 24.7 | 20.1 | |
| 6-20 | 30 | 27.7 | 20.4 | |
| 6-27 | 30 | 27.3 | 30 | |
| 16-20 | 30 | 38.3 | | 40.9 |
| 93-155 | 30 | 64.9 | | 49.6 |
| 174-20 | 30 | 16.7 | 18.9 | |
| 222-1 | 25.3 | 23.1 | | 25.4 |
| 261-20 | 30 | 18.3 | 17.7 | |
| 279-20 | 30 | 24.5 | | 27.5 |
| 279-102 | 30 | 27.1 | | 24.3 |
| 279-126 | 30 | 22.2 | | 18.6 |
| 283-20 | 30 | 27.7 | | 23.7 |
| 283-57 | 30 | 31.4 | | 26.3 |
| 283-102 | 30 | 31.4 | | 19.7 |
| 283-126 | 30 | 27.6 | | 17.7 |
| 283-186 | 30 | 41.3 | | 34.6 |
| 302-20 | 30 | 28 | 23.8 | |
| 302-27 | 30 | 20.4 | 15.1 | |
| 302-126 | 30 | 30 | | 40.7 |

A box is colored green if the Euclidean distance between the Cα atoms is within the threshold or else is colored pink. The distances were calculated using the PyMOL plugin PyXlinkViewer.

**Table S9:** Details of the sequences of the ATPPase subunit used in the alignment shown in Figure S4.

| Abbreviation | Organism | Genbank accession |
|---|---|---|
| Meth.jann | Methanocaldococcus Jannaschii | WP_010870642.1 |
| Meth.ferv | Methanocaldococcus fervens AG86 | ACV24125.1 |
| Pyro.hori | Pyrococcus horikoshii | O59072 |
| Meth.vulc | Methanocaldococcus vulcanius | WP_015733448.1 |
| Meth.infe | Methanocaldococcus infernus | WP_013100352.1 |
| Meth.villo | Methanocaldococcus villosus | WP_004594597.1 |
| Meth.form | Methanotorris formicicus | WP_007044532.1 |
| Meth.ther | Methanothermococcus thermolithotrophicus | WP_018153929.1 |
| Meth.okin | Methanothermococcus okinawensis | WP_013866455.1 |
| Meth.mari | Methanococcus maripaludis | WP_104837196.1 |
| Meth.vann | Methanococcus vannielii | WP_011971846.1 |
| Meth.aeol | Methanococcus aeolicus | WP_011972897.1 |
| Meth.volt | Methanococcus voltae | WP_013179487.1 |
| Meth.tene | Methanothermobacter tenebrarum | WP_112093976.1 |
| Meth.palu | Methanobacterium paludis | WP_013824844.1 |
| Meth.arch | Methanobacteriales archaeon HGW-Methanobacteriales-1 | PKL67884.1 |
| Meth.bovi | Methanobrevibacter boviskoreani | WP_040681779.1 |
| Meth.sp87 | Methanobrevibacter sp. 87.7 | WP_088539025.1 |
| Meth.spMB | Methanobacterium sp. MB1 | WP_023991489.1 |
| Meth.ther1 | Methanothermobacter thermautotrophicus str. Delta H | AAB85215.1 |
| Meth.spSM | Methanobacterium sp. SMA-27 | WP_048189816.1 |
| Cand.hydr | Candidatus Hydrothermarchaeota archaeon | RLG56306.1 |
| Meth.cong | Methanobacterium congolense | WP_071905963.1 |
| Meth.lacu | Methanobacterium lacus | WP_013645922.1 |
| Meth.ferv1 | Methanothermus fervidus | WP_013413589.1 |
| Meth.sp | Methanobacterium sp | RJS49331.1 |
| Meth.madd | Methanobacterium sp. Maddingley MBC34 | EKQ55647.1 |
| Meth.spBR | Methanobacterium sp. BRmetb2 | AXV36938.1 |
| Meth.arbo | Methanobrevibacter arboriphilus | WP_054835928.1 |
| Pala.paci | Palaeococcus pacificus | WP_084177517.1 |
| Meth.oral | Methanobrevibacter oralis | WP_063720427.1 |
| Ther.gamm | Thermococcus gammatolerans EJ3 | ACS33262.1 |
| Meth.cuti | Methanobrevibacter cuticularis | WP_067259482.1 |
| Methanob | Methanobacterium | WP_069583434.1 |
| Meth.smit | Methanobrevibacter smithii DSM 2375 | EEE41124.1 |
| Meth.rumi | Methanobrevibacter ruminantium | WP_012956219.1 |
| Ther.sibi | Thermococcus sibiricus MM 739 | ACS89118.1 |
| Ther.spP6 | Thermococcus sp. P6 | WP_088882269.1 |
| Meth.rhol | Methanosphaera sp. rholeuAM130 | RAP53184.1 |
| Meth.mill | Methanobrevibacter millerae | ALT68479.1 |
| Meth.spNO | Methanobrevibacter sp. NOE | RBQ24062.1 |
| Meth.thau | Methanobrevibacter thaueri | WP_116592073.1 |
| Ther.cele | Thermococcus celer | WP_088862304.1 |
| Meth.olle | Methanobrevibacter olleyae | WP_067146758.1 |
| Meth.curv | Methanobrevibacter curvatus | WP_067090106.1 |
| Cand.alti | Candidatus Altiarchaeales archaeon WOR_SM1_86-2 | ODS39529.1 |
| Methanos | Methanosphaera | WP_011406888.1 |
| arch.bin16 | archaeon BMS3Bbin16 | GBE56826.1 |
| Meth.gott | Methanobrevibacter gottschalkii DSM 11977 | RPF50723.1 |
| arch.bin15 | archaeon BMS3Bbin15 | GBE55171.1 |
| Meth.fili | Methanobrevibacter filiformis | WP_066972131.1 |
| Meth.cuni | Methanosphaera cuniculi | PWL08512.1 |
| Cand.alti1 | Candidatus Altiarchaeales archaeon HGW-Altiarchaeales-3 | PKP54457.1 |
| Cand.alti2 | Candidatus Altiarchaeales archaeon WOR_SM1_SCG | ODS36727.1 |
| Methanop | Methanopyrus sp. KOL6 | WP_088335536.1 |
| Cand.alti3 | Candidatus Altiarchaeum sp. CG2_30_32_3053 | OIQ05869.1 |
| Atop.taba | Atopococcus tabaci | WP_051258565.1 |
| Meth.ther2 | Methanosarcina thermophila | WP_048167447.1 |

**Table S10:** Data acquisition settings for LC-MS/MS.

| | | Mass spec data file name | | | | | |
|---|---|---|---|---|---|---|---|
| Heterodimer Experiment 1 (AB1) | | AB1_01 | AB1_02 | AB1_04 | AB1_05 | AB1_06 | AB1_07 |
| Heterodimer Experiment 2 (AB2) | | AB2_01 | AB2_02 | AB2_04 | AB2_05 | | |
| Heterotetramer Experiment 1 (ABBA1) | | ABBA1_01 | ABBA1_02 | ABBA1_04 | ABBA1_05 | ABBA1_06 | ABBA1_07 |
| Heterotetramer Experiment 2 (ABBA2) | | ABBA2_01 | ABBA2_02 | ABBA2_04 | ABBA2_05 | ABBA2_06 | ABBA2_07 |
| | | **Data acquisition settings** | | | | | |
| **Full MS** | **Resolution** | 60,000 | 60,000 | 120,000 | 120,000 | 120,000 | 120,000 |
| | **AGC target** | 1.00E+06 | 1.00E+06 | 1.00E+06 | 1.00E+06 | 3.00E+06 | 3.00E+06 |
| | **Maximum IT** | 50 ms | 50 ms | 50 ms | 50 ms | 50 ms | 50 ms |
| | **Scan Range** | 350 to 2000 m/z | 350 to 2000 m/z | 350 to 2000 m/z | 350 to 2000 m/z | 350 to 1600 m/z | 350 to 1600 m/z |
| **dd MS2** | **Resolution** | 15,000 | 15,000 | 30,000 | 30,000 | 60, 000 | 60, 000 |
| | **AGC target** | 1.00E+05 | 1.00E+05 | 1.00E+05 | 1.00E+05 | 5.00E+04 | 5.00E+04 |
| | **Maximum IT** | 100 ms | 100 ms | 100 ms | 100 ms | 120 ms | 120 ms |
| | **Loop count** | 10 | 10 | 10 | 10 | 10 | 10 |
| | **Isolation window** | 2.0 m/z | 2.0 m/z | 1.6 m/z | 1.6 m/z | 1.4 m/z | 1.4 m/z |
| | **Fixed first mass** | 100.0 m/z | 100.0 m/z | 100.0 m/z | 100.0 m/z | 100.0 m/z | 100.0 m/z |
| | **NCE** | 27 | 28, 30, 32 | 30 | 27 | 27 | 23,25,27 |
| | **Spectrum data type** | Centroid | Centroid | Centroid | Centroid | Centroid | Centroid |
| **dd settings** | **Min AGC target** | 5.00E+03 | 5.00E+03 | 5.00E+03 | 5.00E+03 | 1.00E+03 | 1.00E+03 |
| | **Charge state exclusion** | 1, 8, >8 | 1, 8, >8 | 1, >8 | 1, >8 | 1, 2, 8, >8 | 1, 2, 8, >8 |
| | **Exclude isotopes** | on | on | on | on | on | on |
| | **Dynamic exclusion** | 5 s | 5 s | 5 s | 5 s | 5 s | 5 s |
