## Supplementary file 3A for "Mechanistic insights into the functioning of GMP synthetase: a two-subunit, allosterically regulated, ammonia tunnelling enzyme"

**Supplementary file 3A: Annotation of the MS/MS spectra of inter-subunit cross-linked peptides identified in the MjATPPase-MjGATase heterodimer complex (AB).**

The crosslinked peptides that are common across the replicate experiments are listed in the Supplementary excel file. These were examined by visualizing the MS/MS spectra using the program pLabel (doi: 10.1002/rcm.3173), which annotates the b ions and y ions of the cross-linked peptides. The spectra were further analysed to confirm the presence of fragment ions that enable unambiguous identification of the crosslinked Lys residues or N-terminus. The presence of such fragment ions is indicated using the notation given below.

- 1+ charged
- 1+ charged with isotopic distribution
- 1+ charged, with water loss or $NH_3$ loss
- 2+ charged
- 2+ charged with isotopic distribution
- 3+ charged
- 3+ charged with isotopic distribution

In each of the following pages, the identity of the crosslinked peptide is shown at the top. The MS/MS spectra from the replicate 1 and 2 are shown at the top and bottom, respectively.

1 MjATPPase(6)-MjGATase(20) KFIDEAVEEIKQQISDR(1)-SLKYIGVSSK(3)

Serial number of the cross-link

MjATPPase cross-linked residue

Position of the cross-linked residue in the MjATPPase tryptic peptide

MjGATase cross-linked residue

Position of the cross-linked residue in the MjGATase tryptic peptide

### 1 MjATPPase(6)-MjGATase(20) KFIDEAVEEIKQQISDR(1)-SLKYIGVSSK(3)

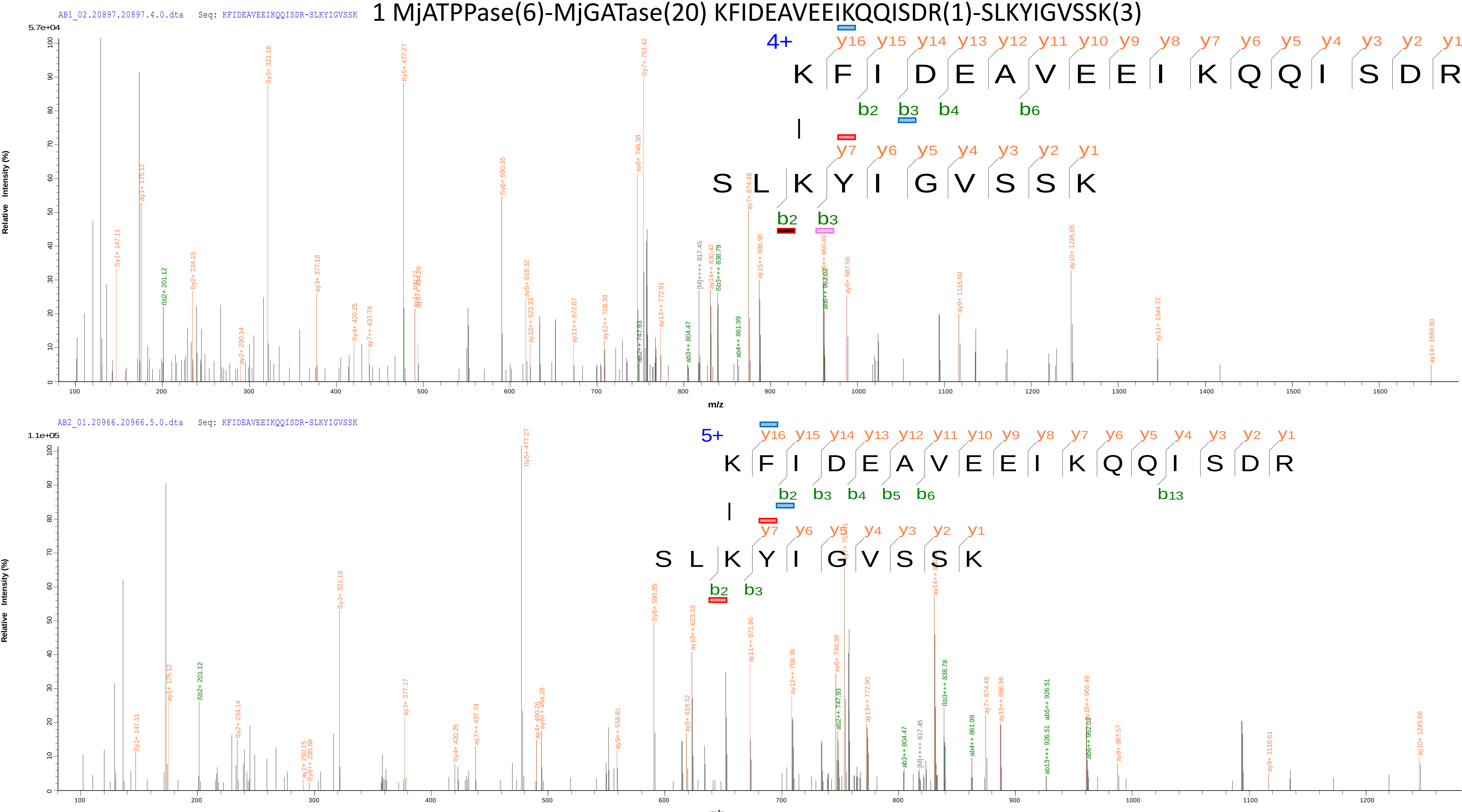

2 MjATPPase(1)-MjGATase(20) MFDPKK(1)-SLKYIGVSSK(3)

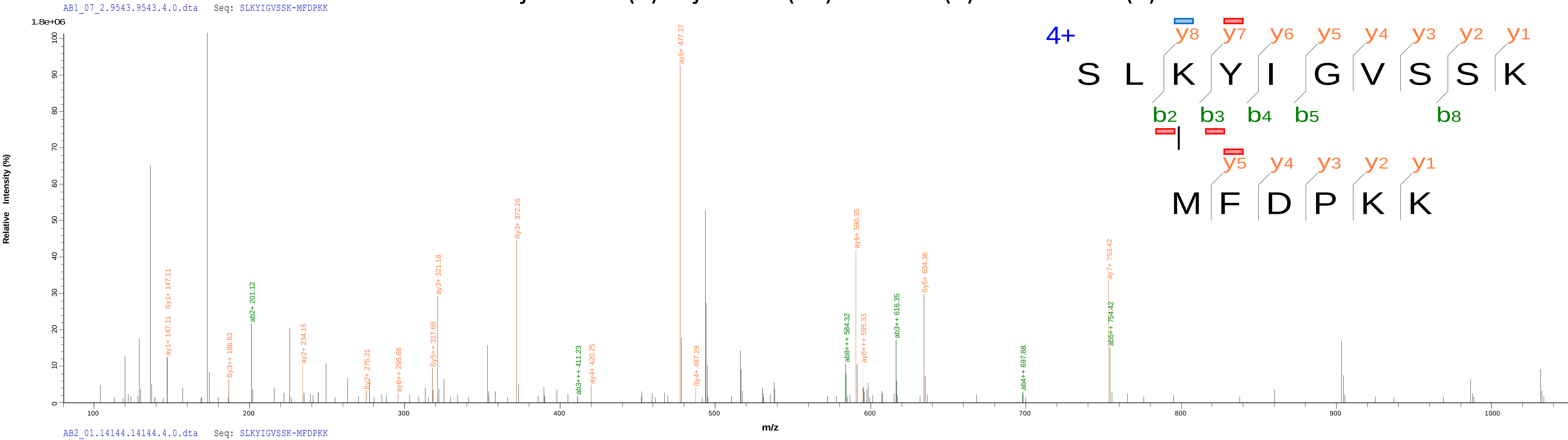


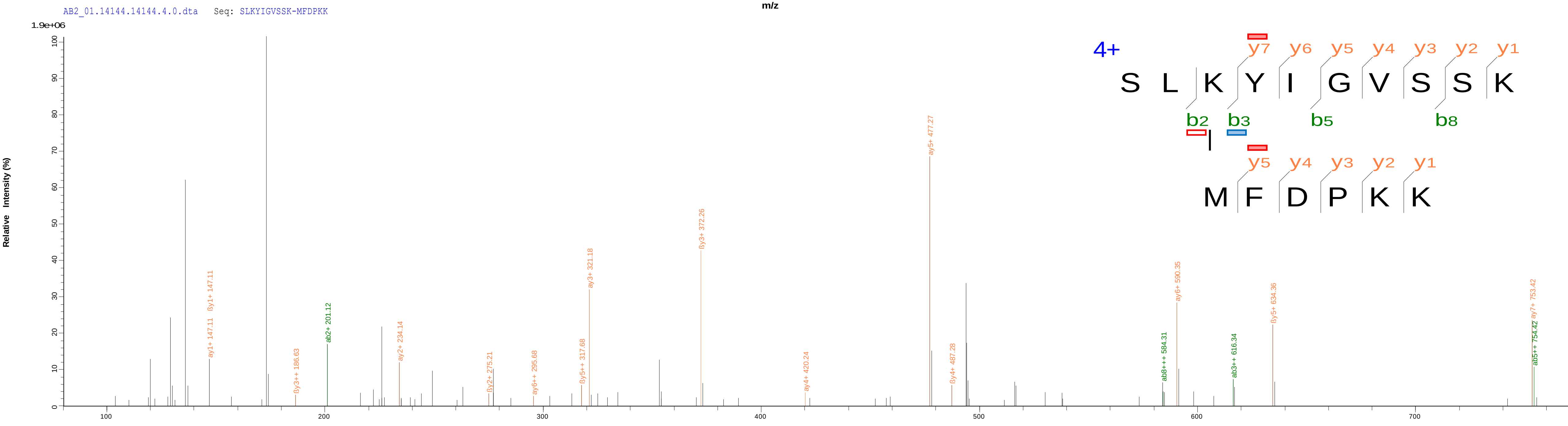

### 3 MjATPPase(302)-MjGATase(20) VVFDITDKPPATIEFE(8)-SLKYIGVSSK(3)

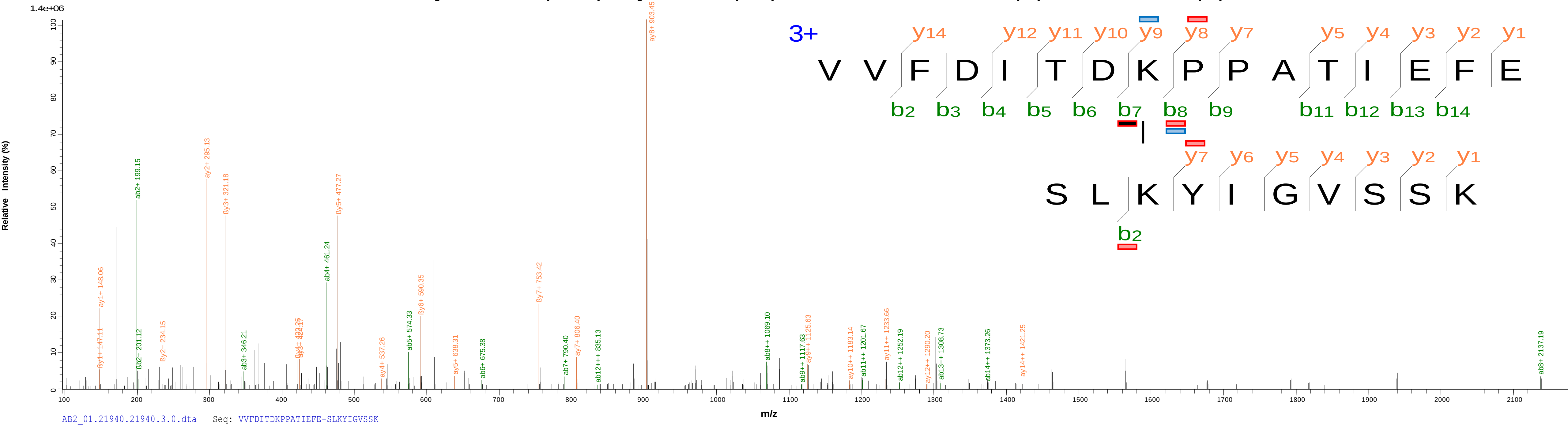


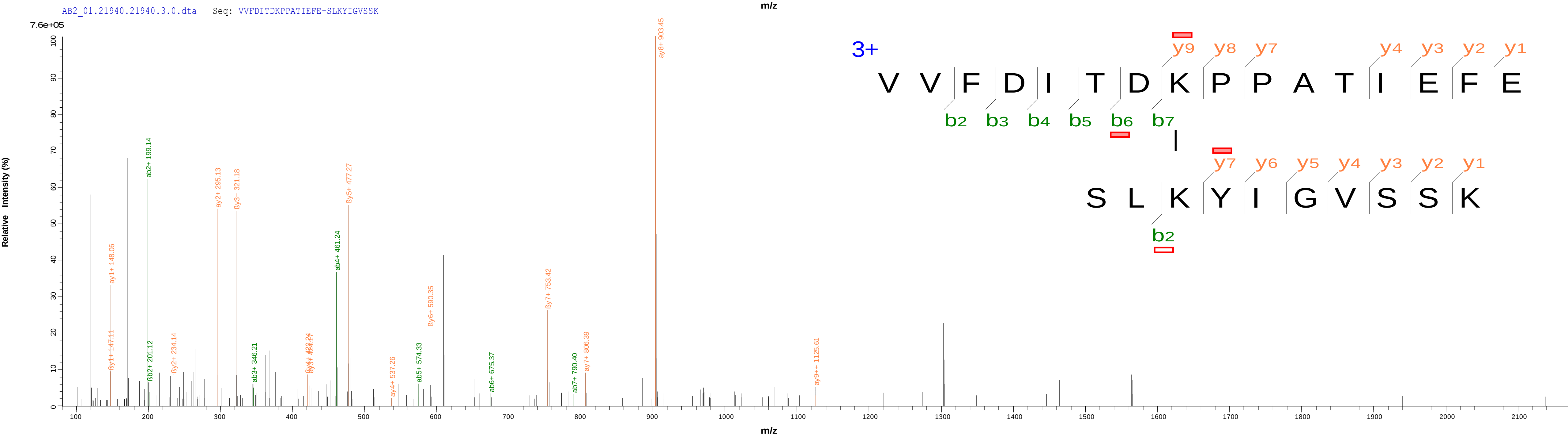

### 4 MjATPPase(16)-MjGATase(20) FIDEAVEEIKQQISDR(10)-SLKYIGVSSK(3)

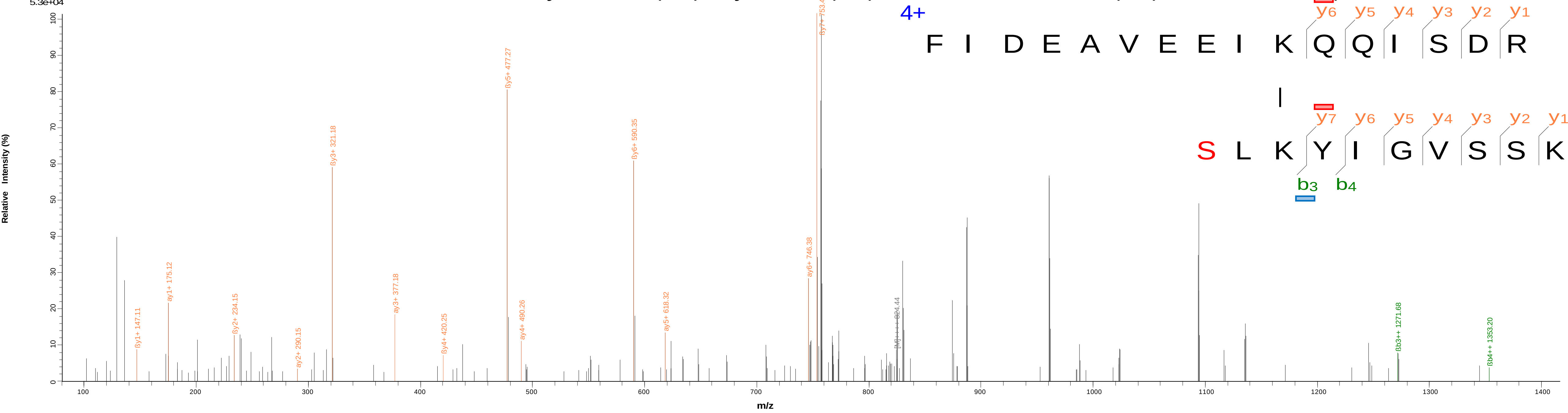


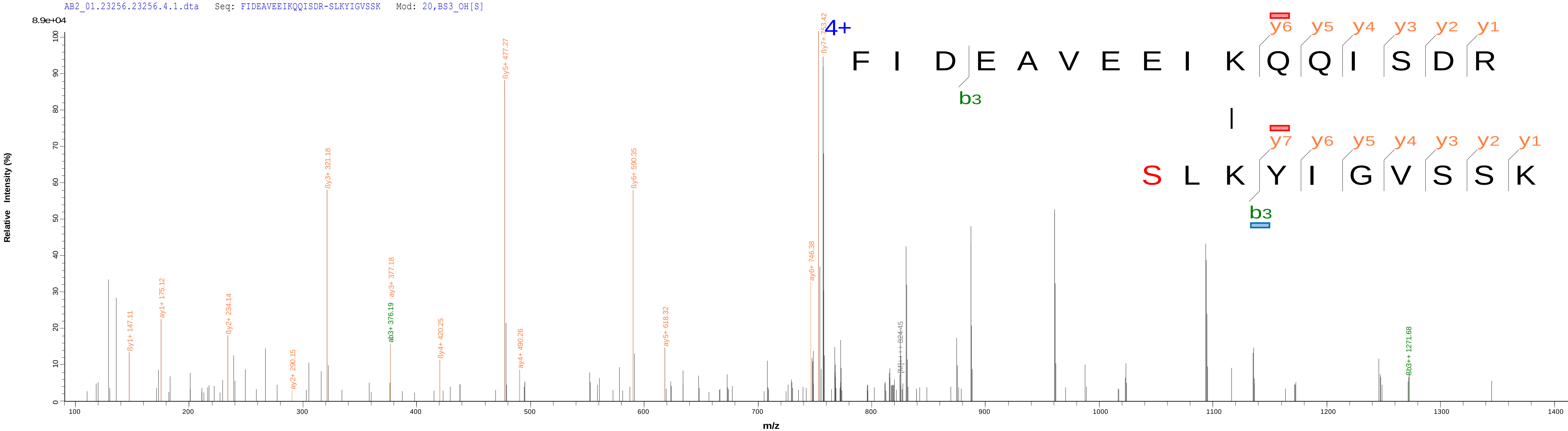

### 5 MjATPPase(283)-MjGATase(20) ISKR(3)-SLKYIGVSSK(3)

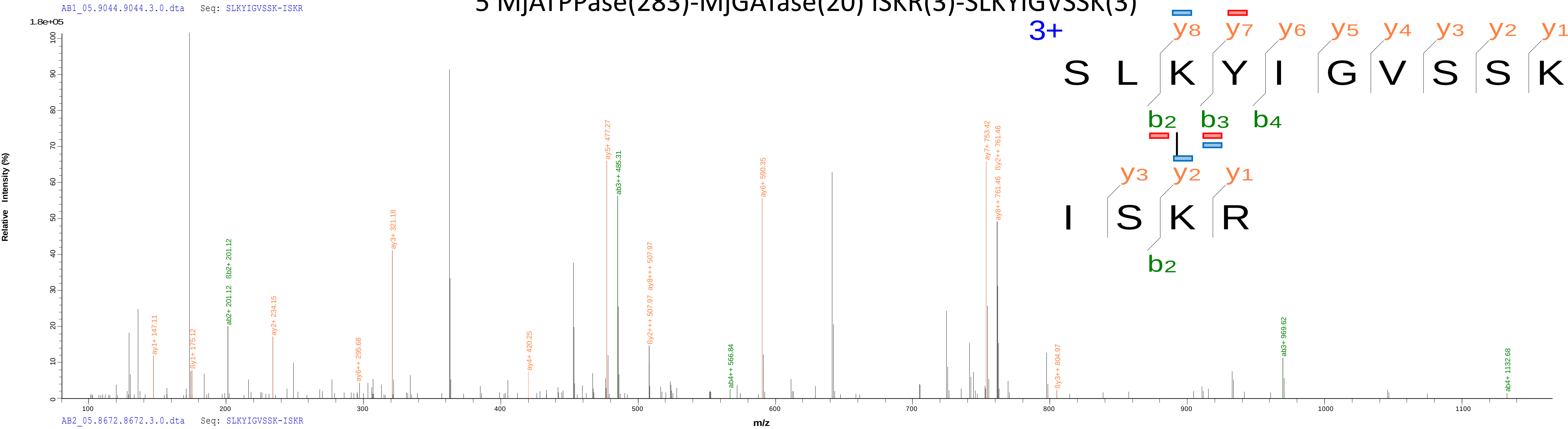


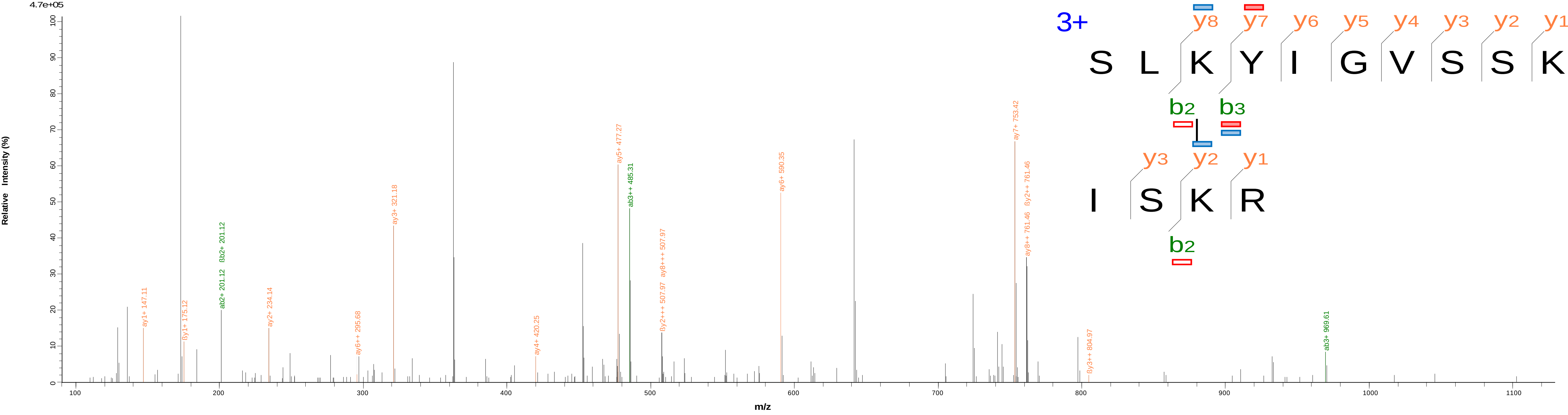

### 6 MjATPPase(279)-MjGATase(20) SLDAMTAHVPEIPFDLLKR(18)-SLKYIGVSSK(3)

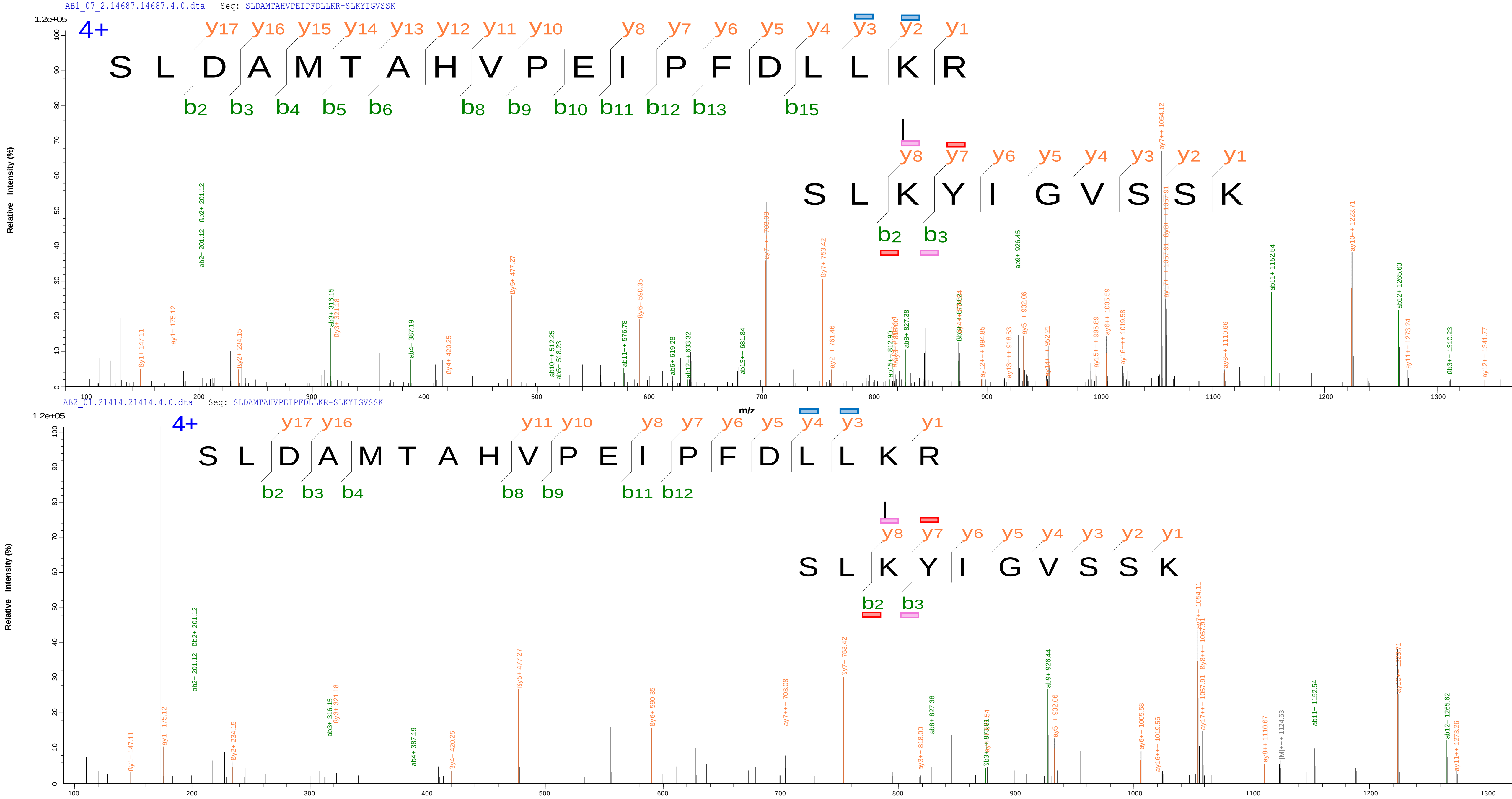

### 7 MjATPPase(174)-MjGATase(20) LLAKELGLPDSIVYR(4)-SLKYIGVSSK(3)

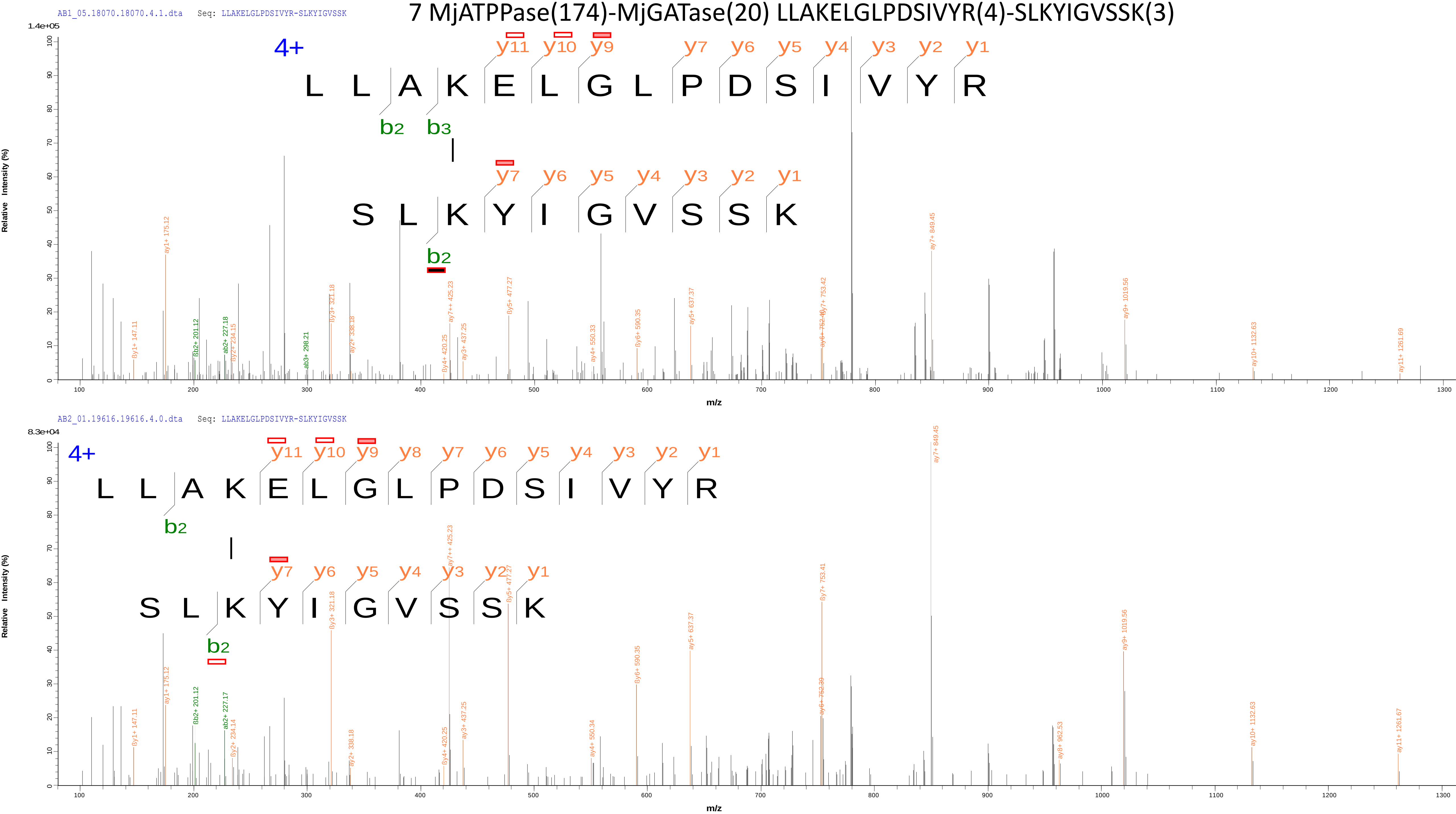

### 8 MjATPPase(261)-MjGATase(20) MVKSLDAMTAHVPEIPFDLLKRISK(3)-SLKYIGVSSK(3)

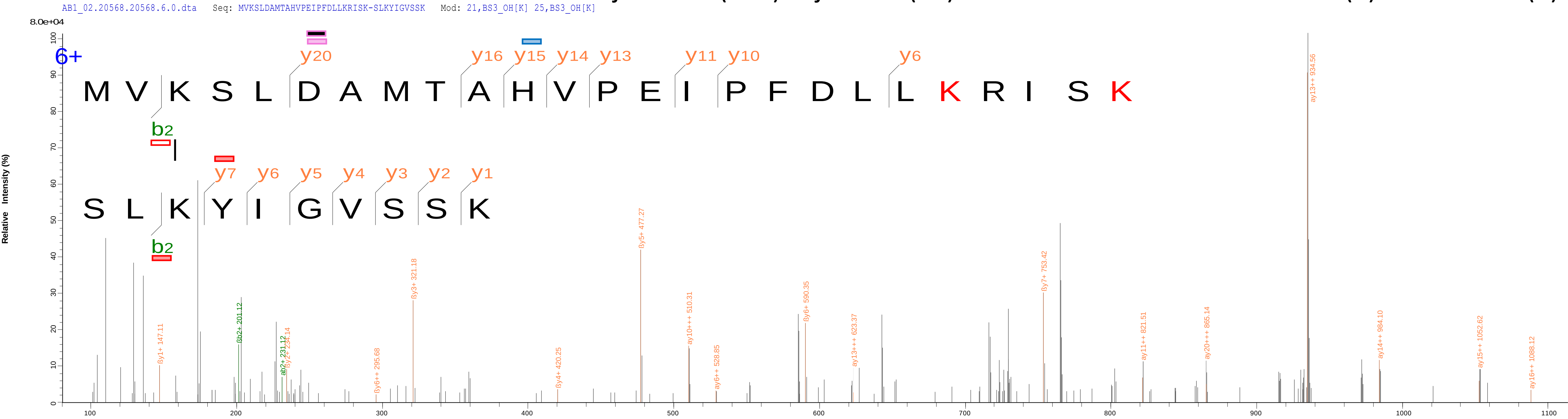


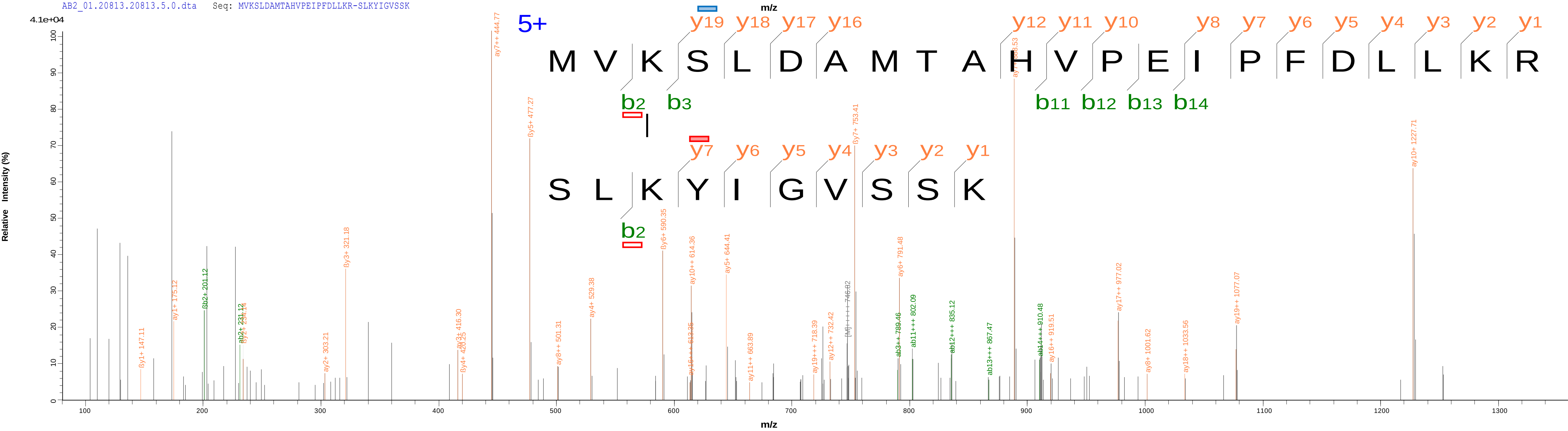

9 MjATPPase(1)-MjGATase(1) MFDPKK(1)-MIVILDNGGQYVHR(1)

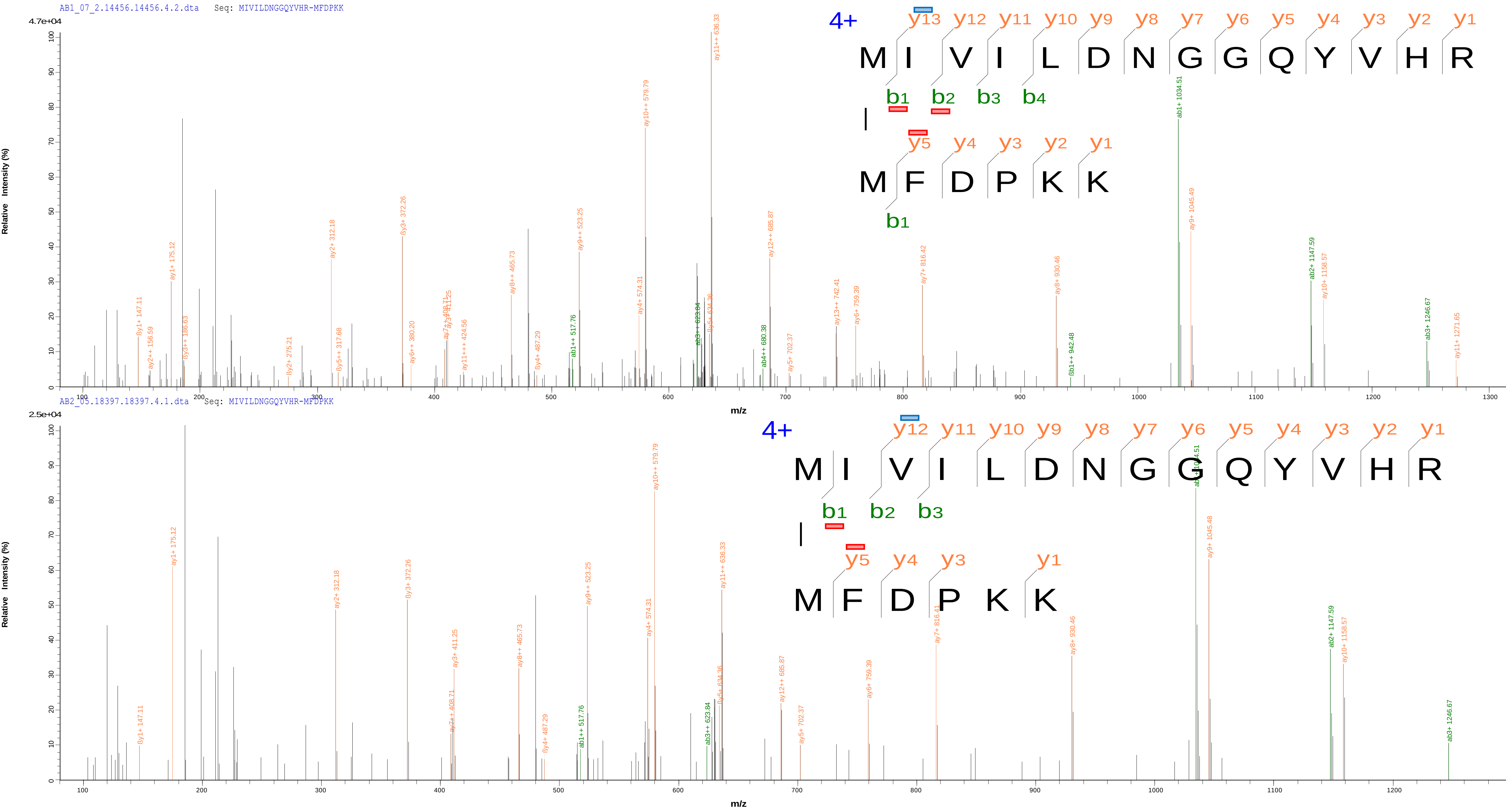

10 MjATPPase(279)-MjGATase(102) SLDAMTAHVPEIPFDLLKR(18)-AEAEEYALTKVYVDKENDLFKNVPR(10)

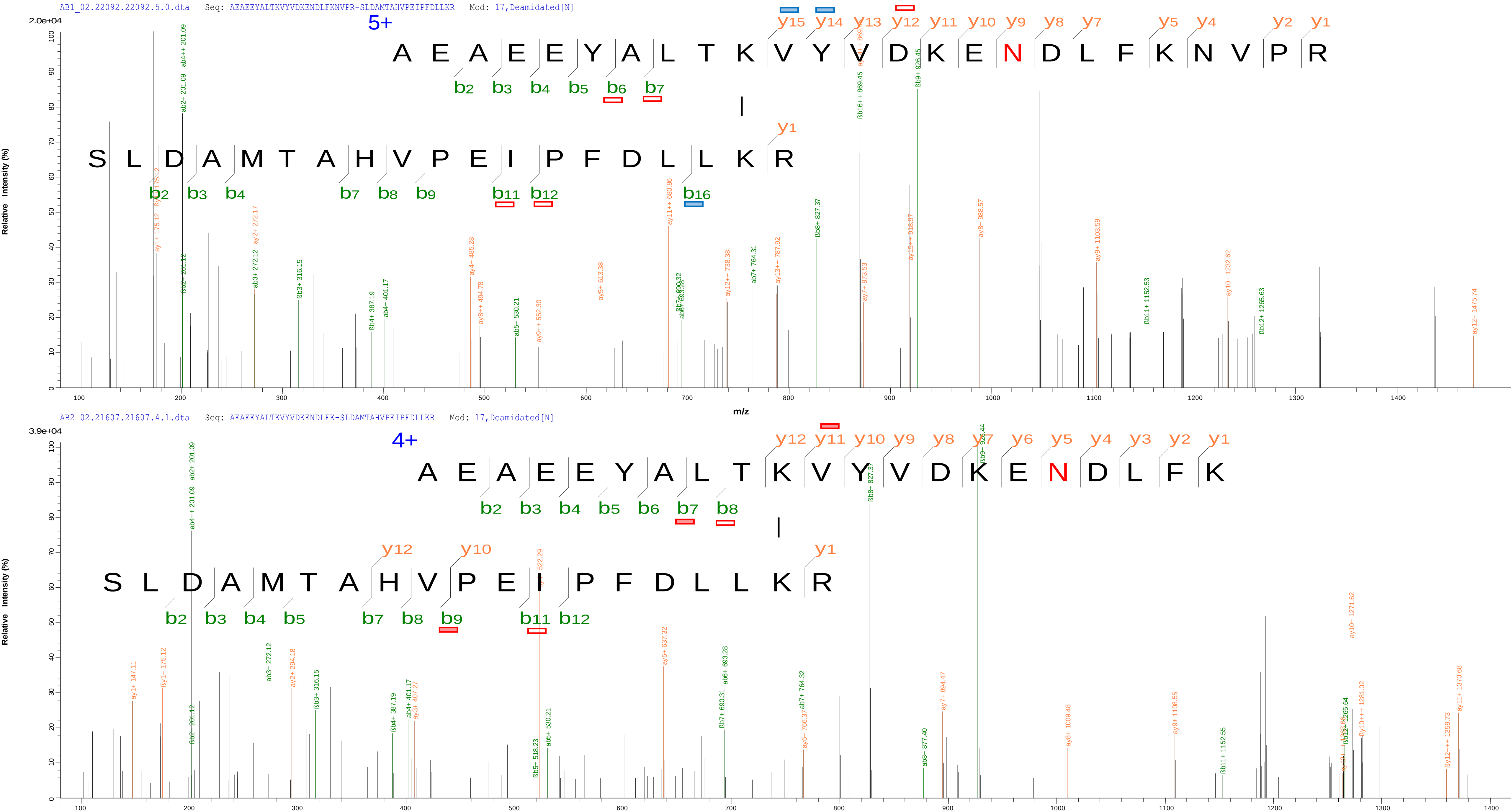

### 11 MjATPPase(6)-MjGATase(27) KFIDEAVEEIKQQISDRK(1)-SLKYIGVSSKIVPNTTPLEEIESNKEVK(10)

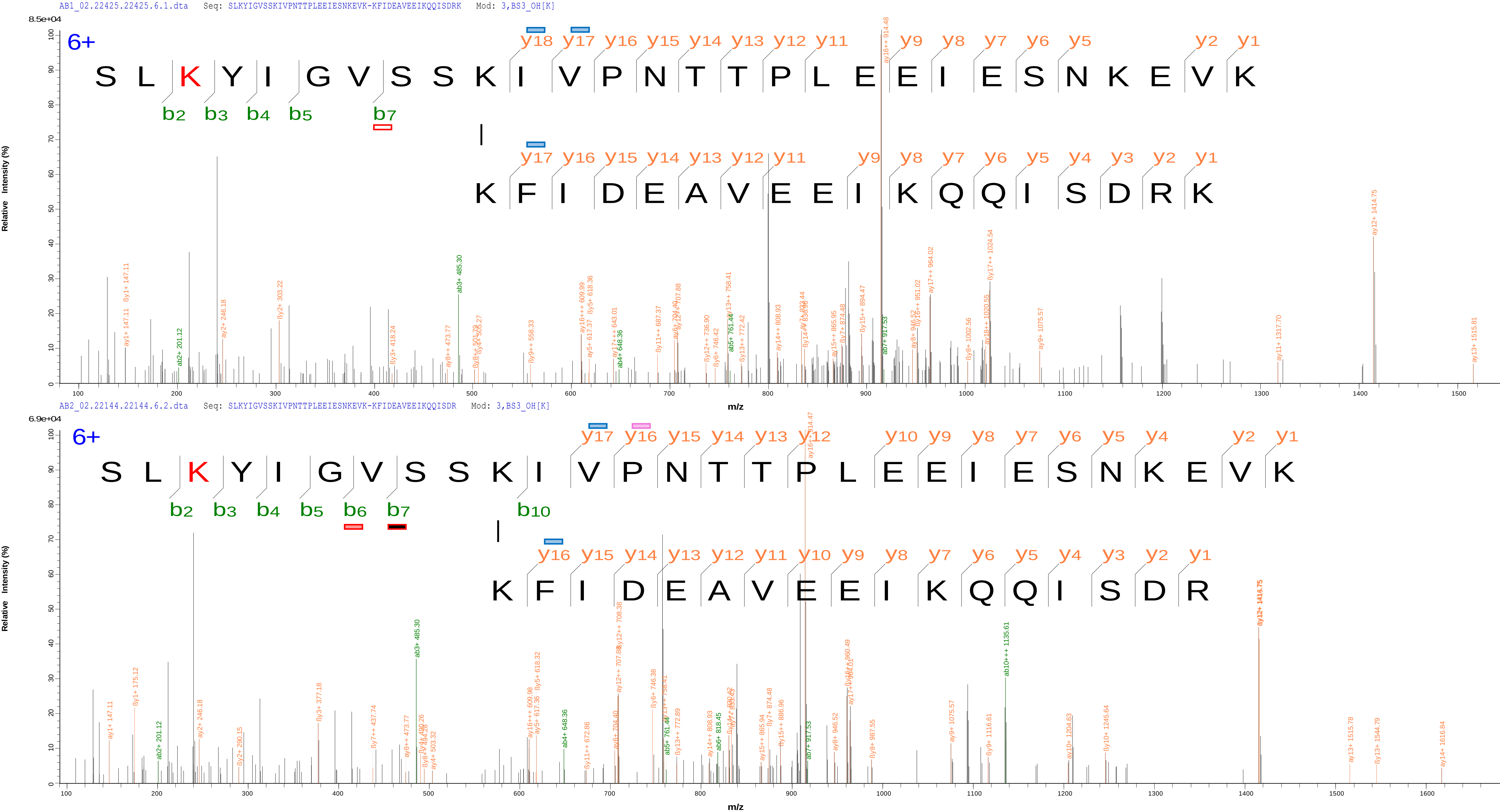

### 12 MjATPPase(1)-MjGATase(177) MFDPKK(1)-TKPIYGVQFHPEVAHTEYGNEILKNFCK(24)

AB1_07_2.11259.11259.6.0.dta Seq: TKPIYGVQFHPEVAHTEYGNEILKNFCK-MFDPKK Mod: 27,Carbamidomethyl[C]

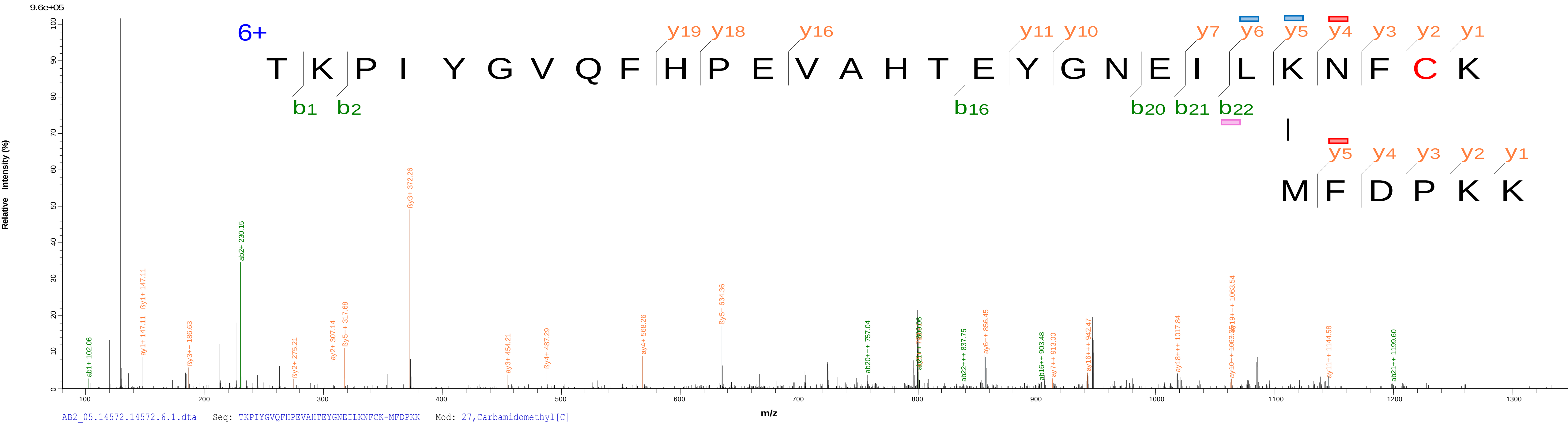


AB2_05.14572.14572.6.1.dta Seq: TKPIYGVQFHPEVAHTEYGNEILKNFCK-MFDPKK Mod: 27,Carbamidomethyl[C]

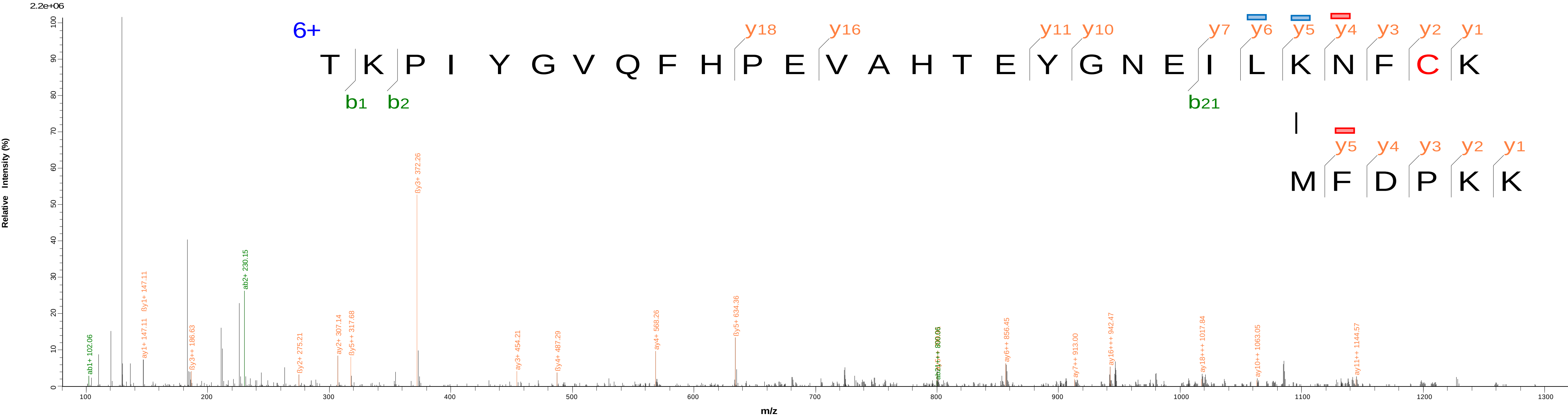

### 13 MjATPPase(1)-MjGATase(186) MFDPKK(1)-VCGYKFE(5)

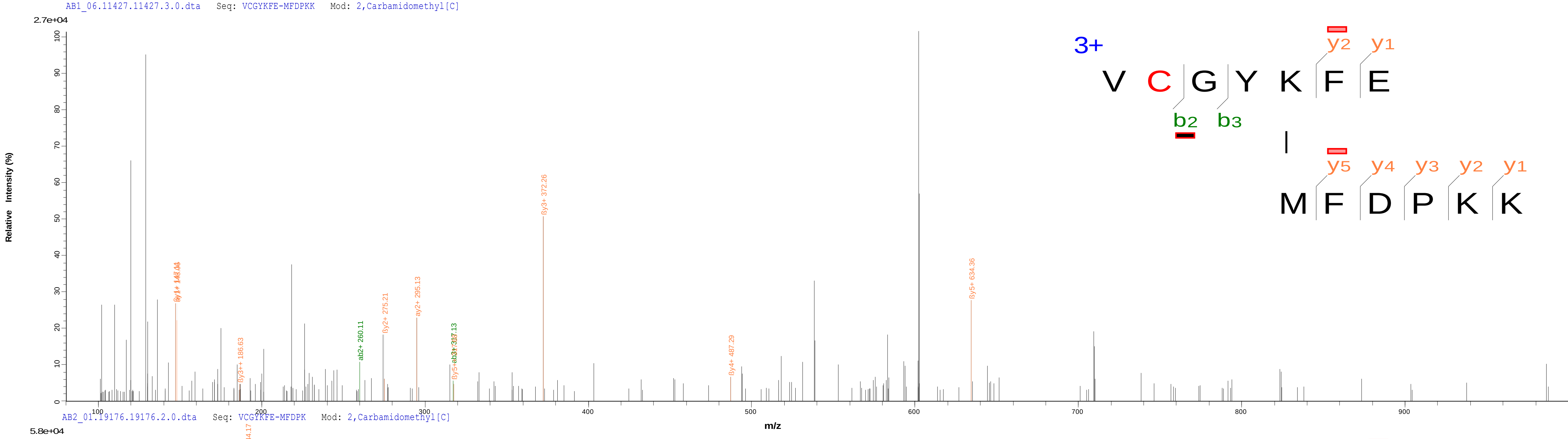


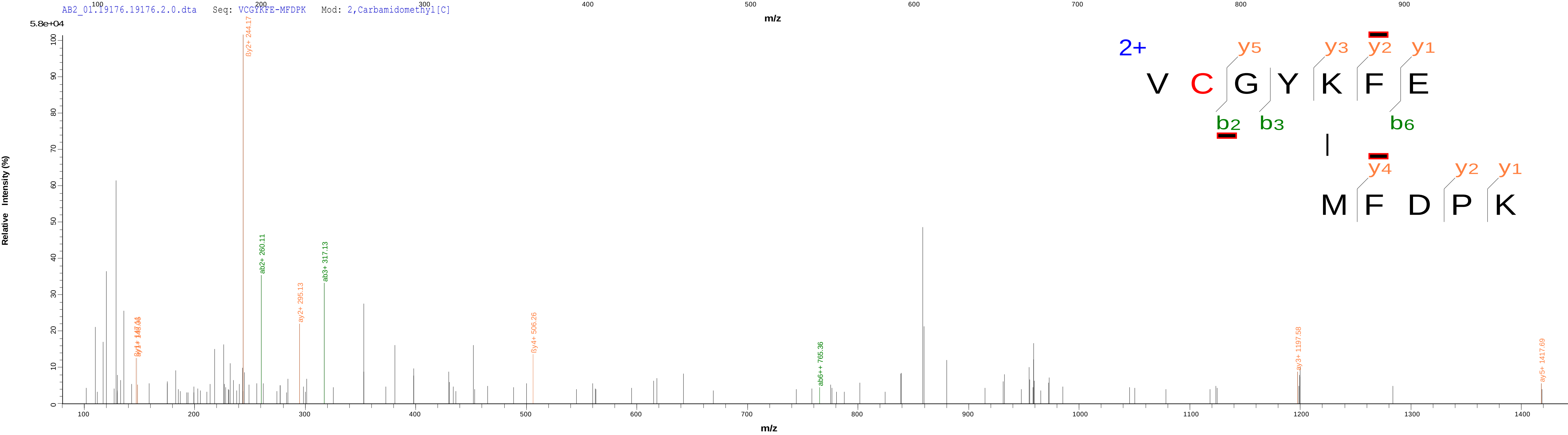

### 14 MjATPPase(279)-MjGATase(126) SLDAMTAHVPEIPFDLLKR(18)-EFNAWASHKDEVK(9)

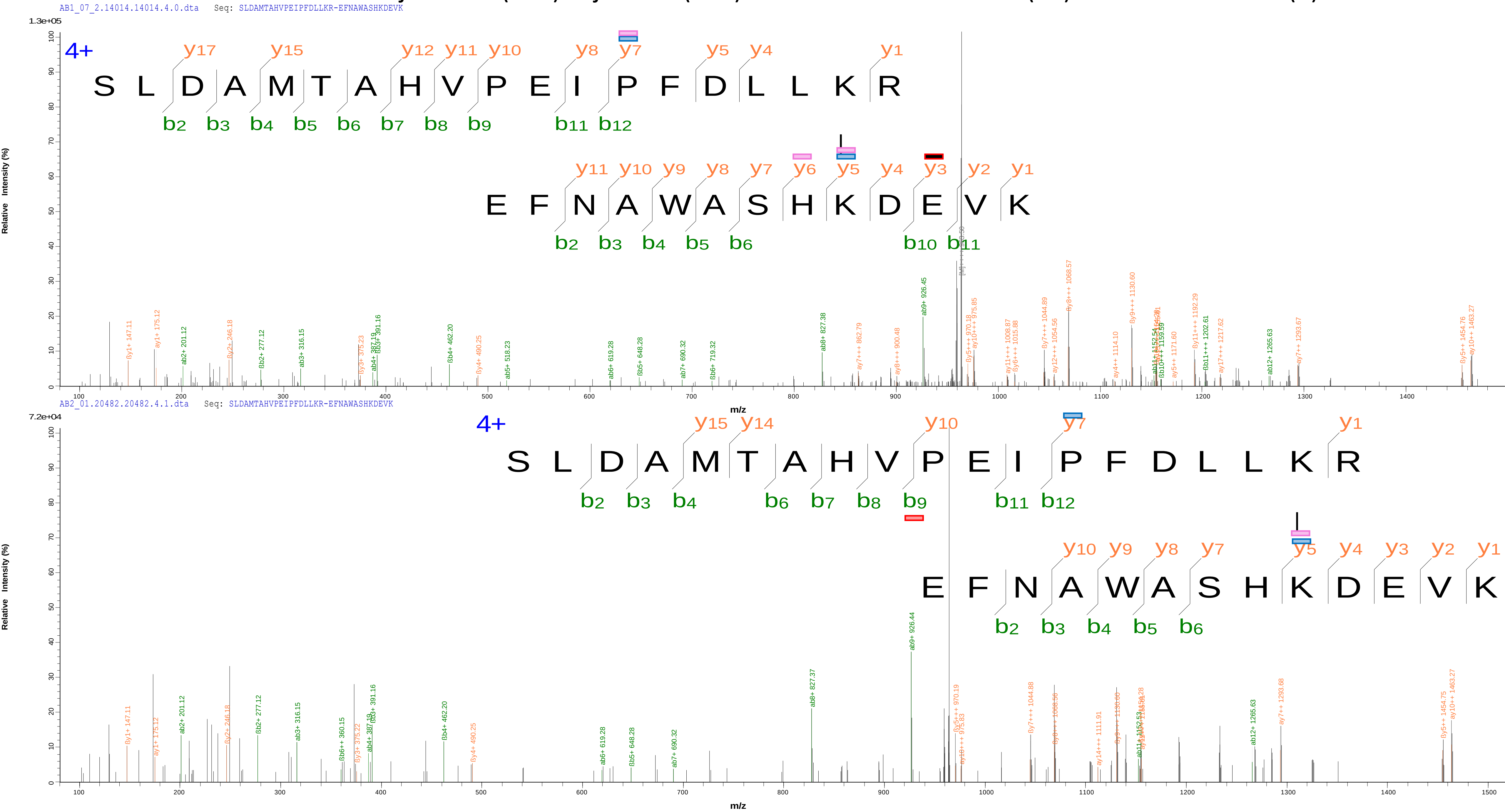

### 15 MjATPPase(1)-MjGATase(27) MFDPK(1)-SLKYIGVSSKIVPNTTPLEEIESNKEVK(10)

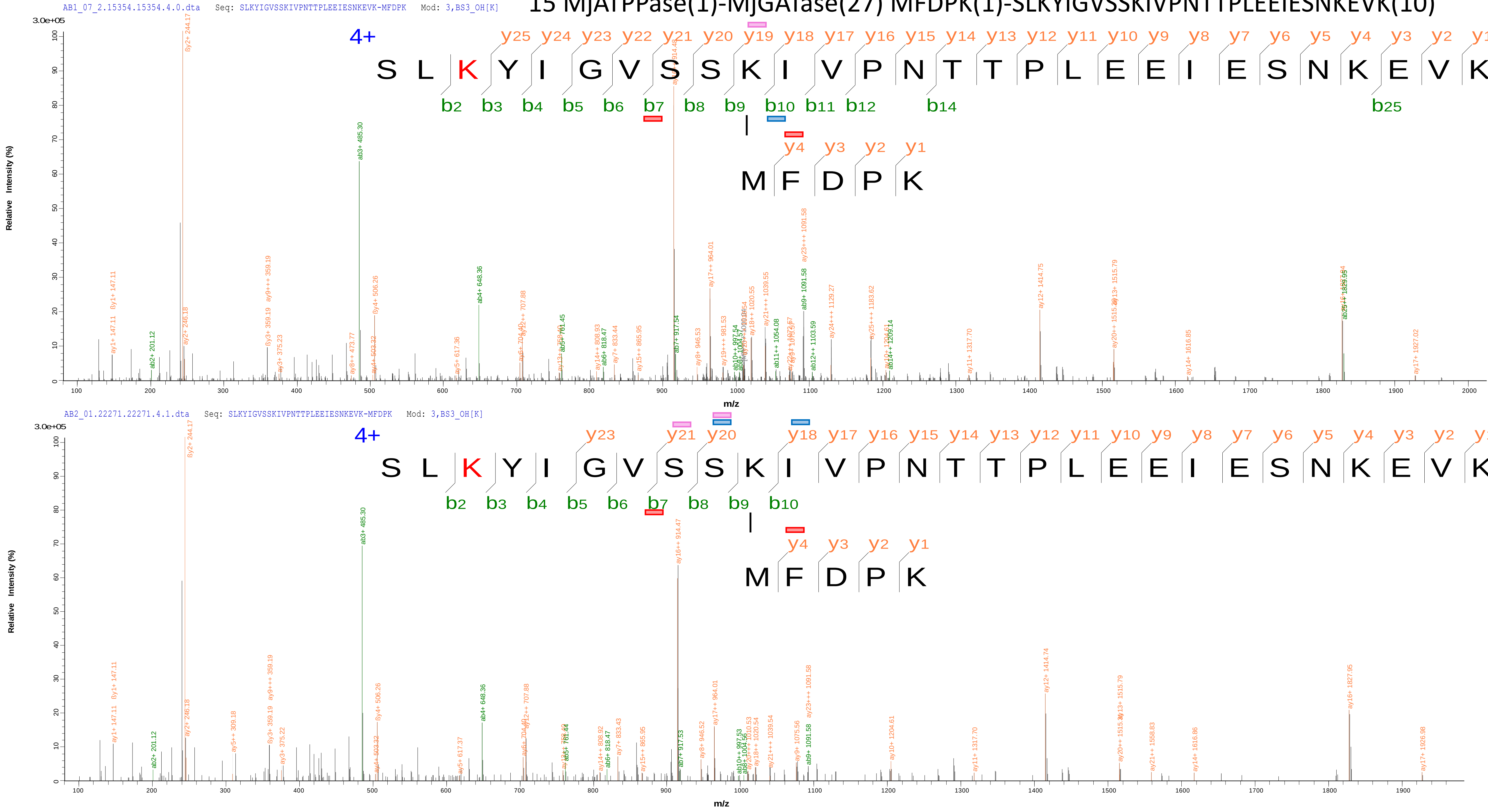

16 MjATPPase(302)-MjGATase(27) VVFDITDKPPATIEFE(8)-SLKYIGVSSKIVPNTTPLEEIESNKEVK(10)

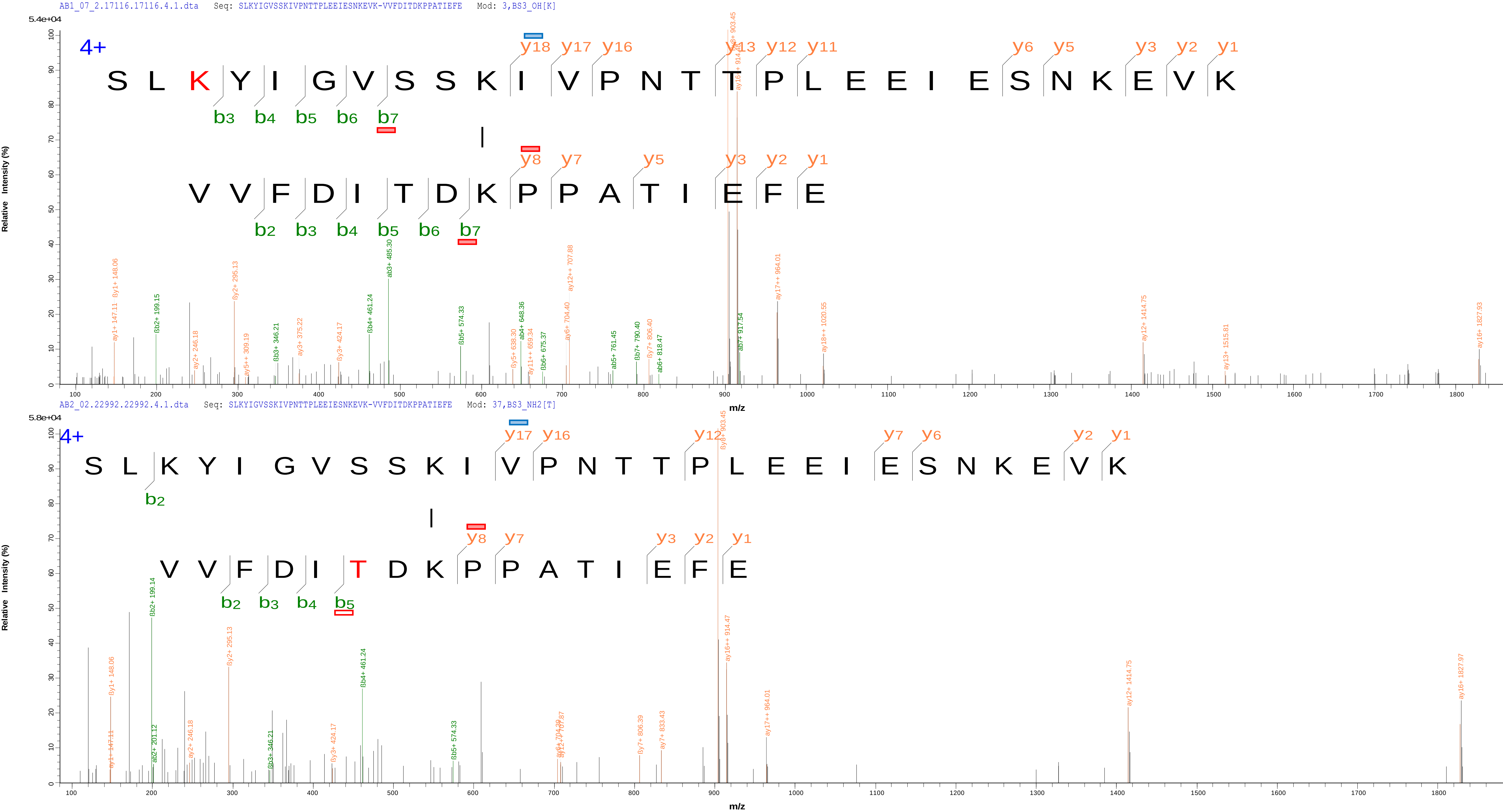

### 17 MjATPPase(283)-MjGATase(57) ISKR(3)-GIILSGGPDIEKAK(12)

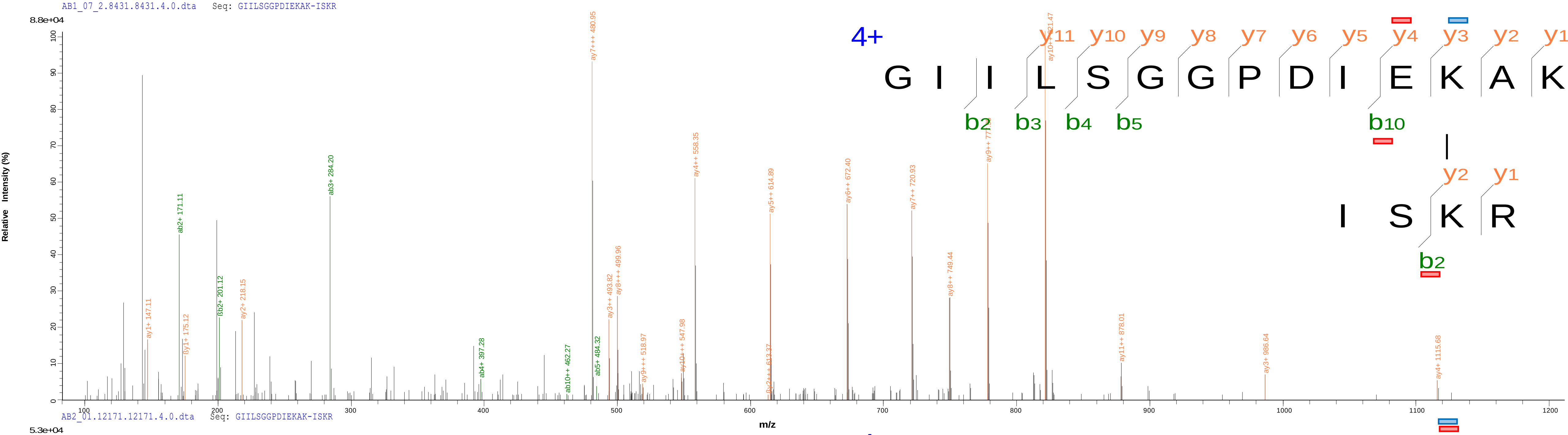


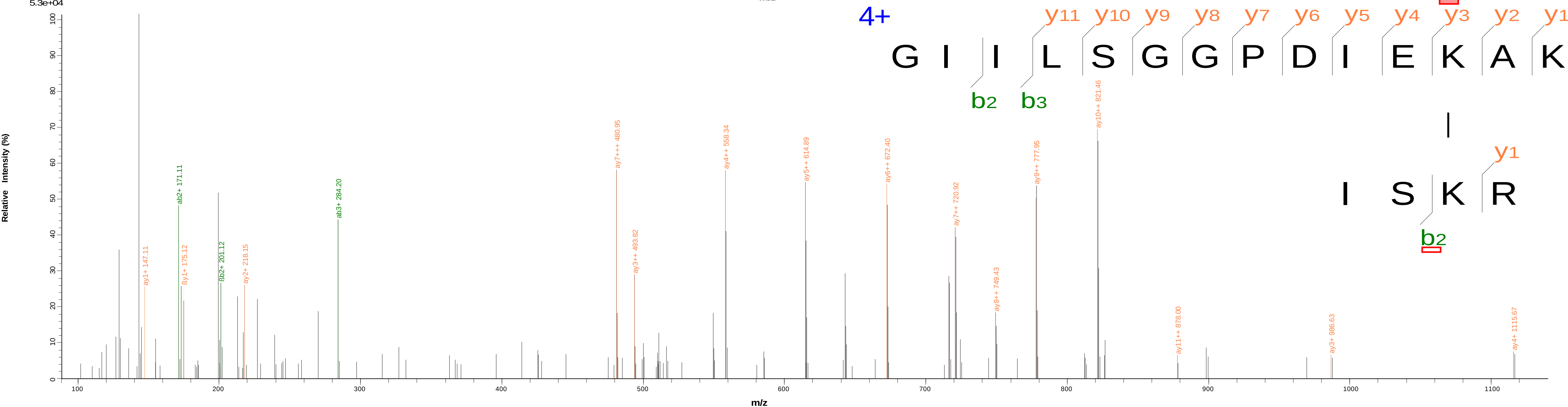

### 18 MjATPPase(302)-MjGATase(126) VVFDITDKPPATIEFE(8)-EFNAWASHKDEVK(9)

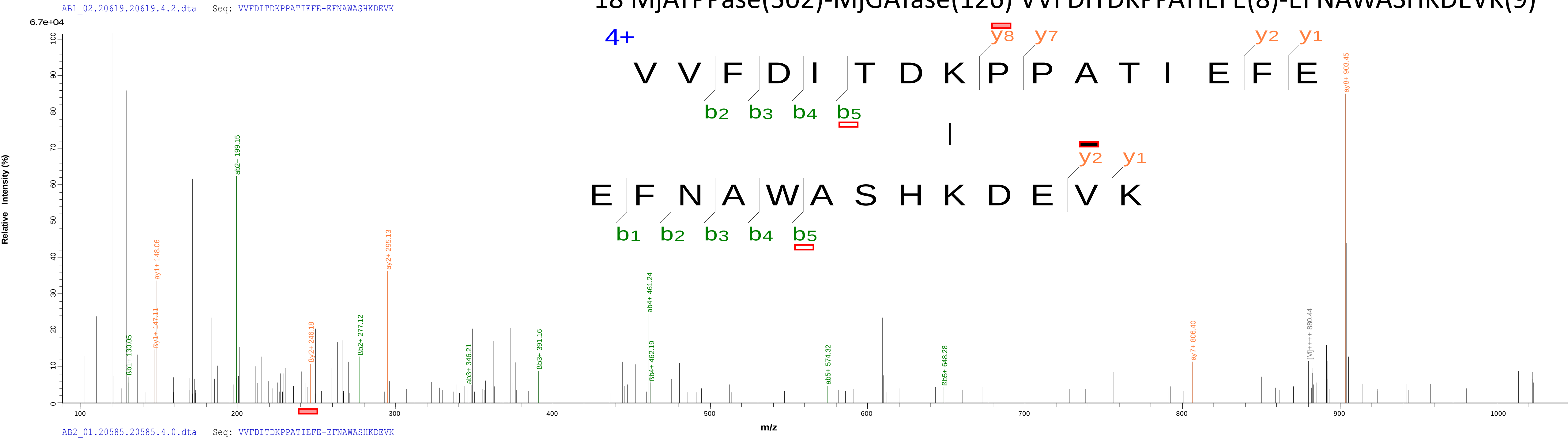


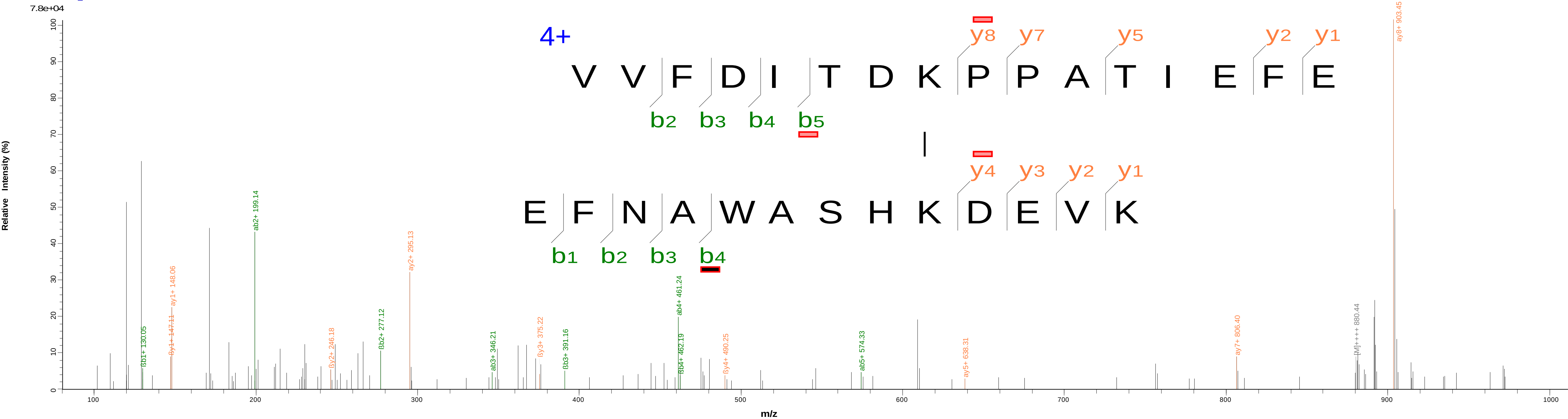

### 19 MjATPPase(283)-MjGATase(186) ISKR(3)- VCGYKFE(5)

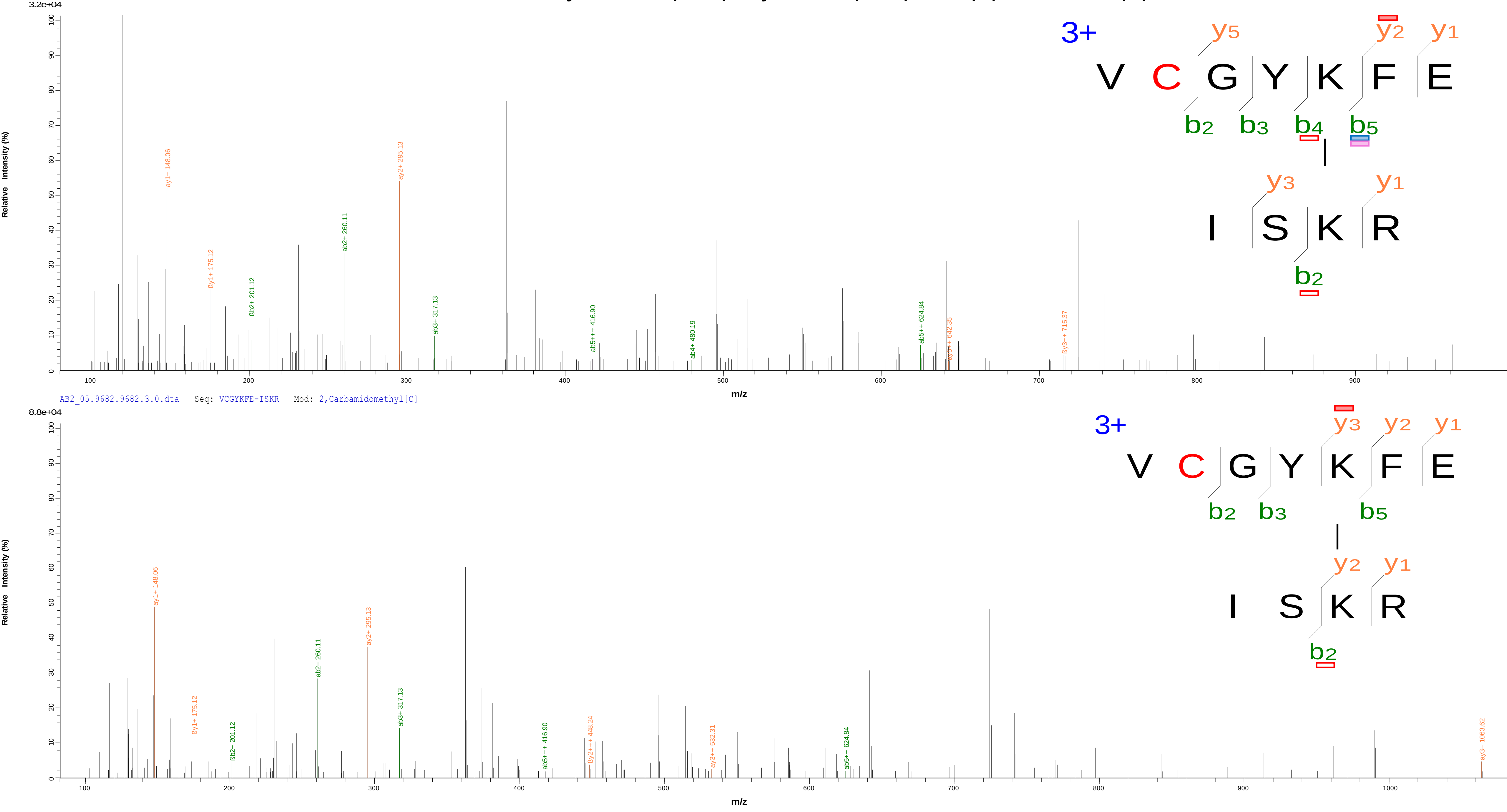

20 MjATPPase(283)-MjGATase(102) ISKR(3)-AEAEEYALTKVYVDKENDLFKNVPR(10)

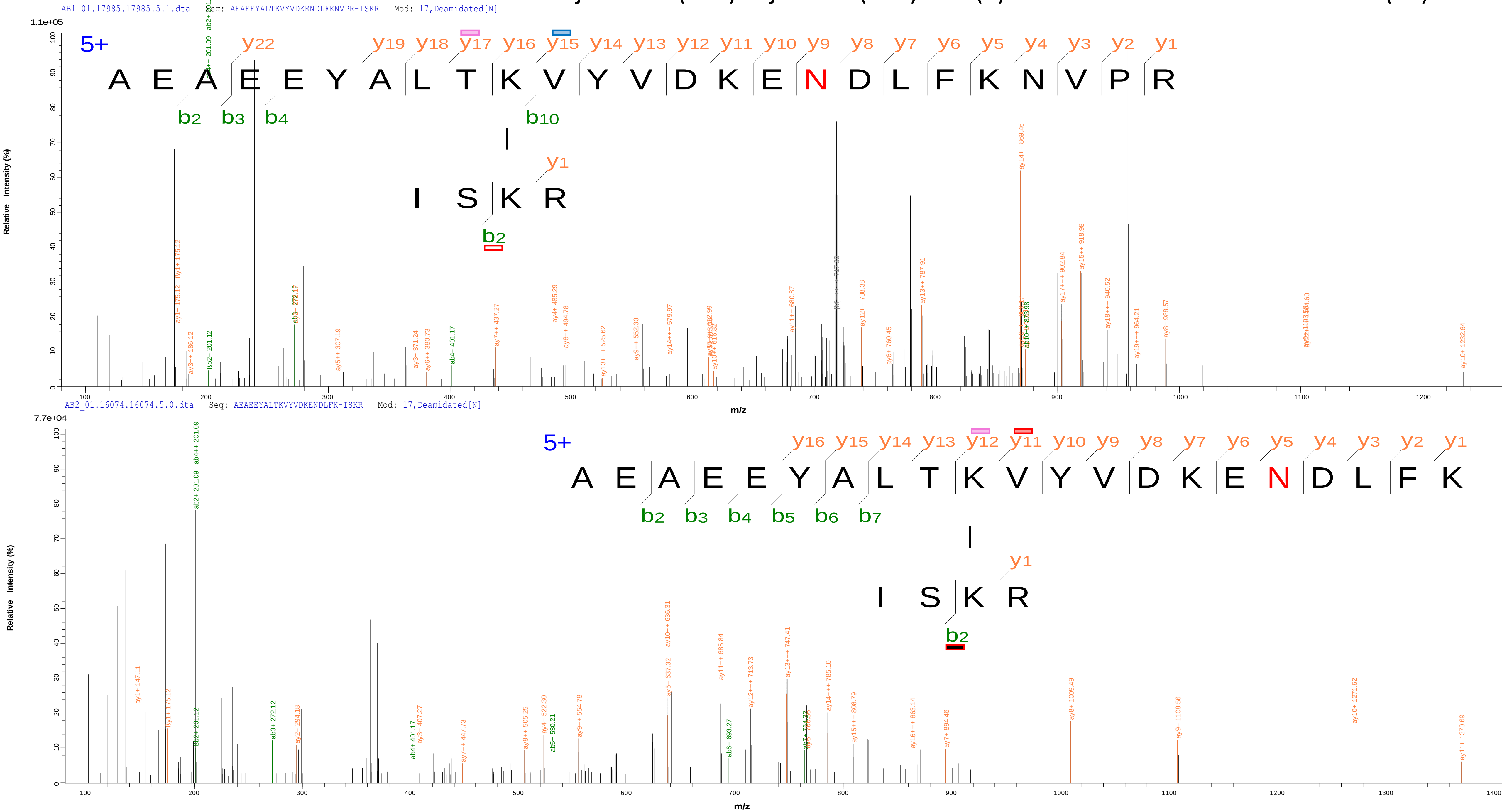

### 21 MjATPPase(283)-MjGATase(126) ISKR(3)-EFNAWASHKDEVK(9)

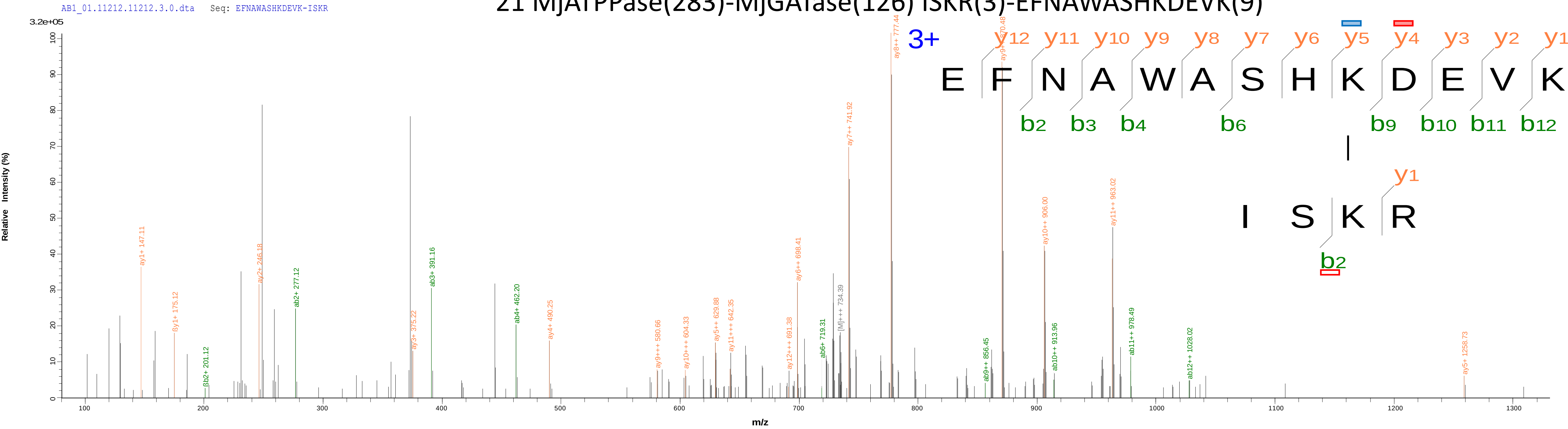


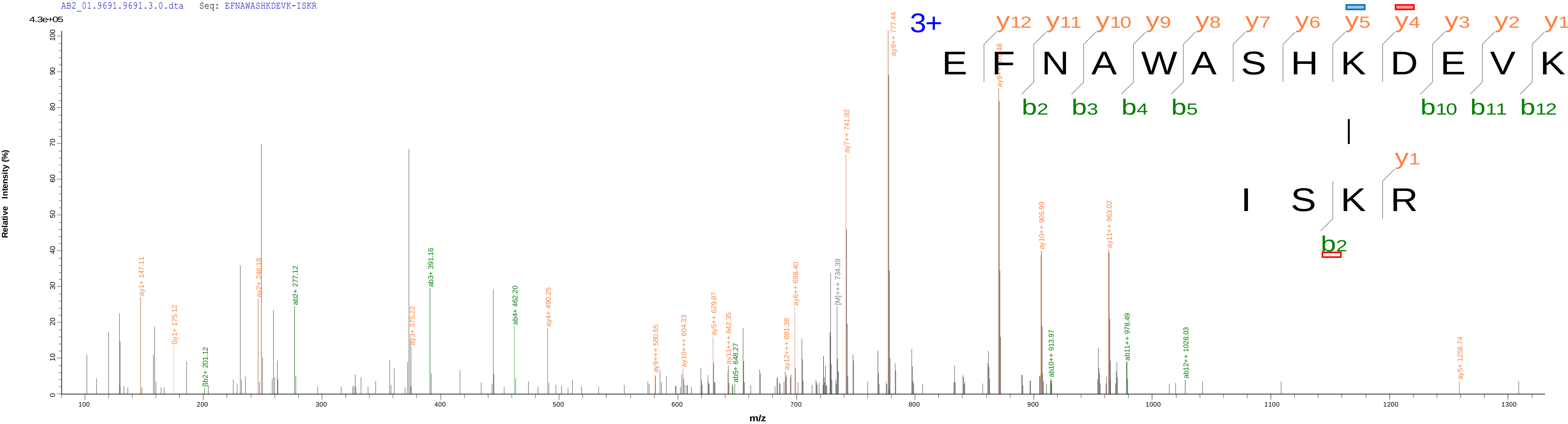
