## Supplementary file 3B for "Mechanistic insights into the functioning of GMP synthetase: a two-subunit, allosterically regulated, ammonia tunnelling enzyme"

**Supplementary file 3B: Annotation of the MS/MS spectra of inter-subunit cross-linked peptides identified in the MjATPPase-MjGATase heterodimer complex (AB).**

The crosslinked peptides that are common across the replicate experiments are listed in the Supplementary excel file. These were examined by visualizing the MS/MS spectra using the program pLabel (doi: 10.1002/rcm.3173), which annotates the b ions and y ions of the cross-linked peptides. The spectra were further analysed to confirm the presence of fragment ions that enable unambiguous identification of the crosslinked Lys residues or N-terminus. The presence of such fragment ions is indicated using the notation given below.

- 1+ charged
- 1+ charged with isotopic distribution
- 1+ charged, with water loss or $NH_3$ loss
- 2+ charged
- 2+ charged with isotopic distribution
- 2+ charged, with water loss or $NH_3$ loss
- 3+ charged
- 3+ charged with isotopic distribution

In each of the following pages, the identity of the crosslinked peptide is shown at the top. The MS/MS spectra from the replicate 1 and 2 are shown at the top and bottom, respectively.

1 MjATPPase(6)-MjGATase(20) KFIDEAVEEIKQQISDR(1)-SLKYIGVSSK(3)

Serial number of the cross-link

MjATPPase cross-linked residue

Position of the cross-linked residue in the MjATPPase tryptic peptide

MjGATase cross-linked residue

Position of the cross-linked residue in the MjGATase tryptic peptide

### 1 MjATPPase(6)-MjGATase(20) KFIDEAVEEIKQQISDR(1)-SLKYIGVSSK(3)

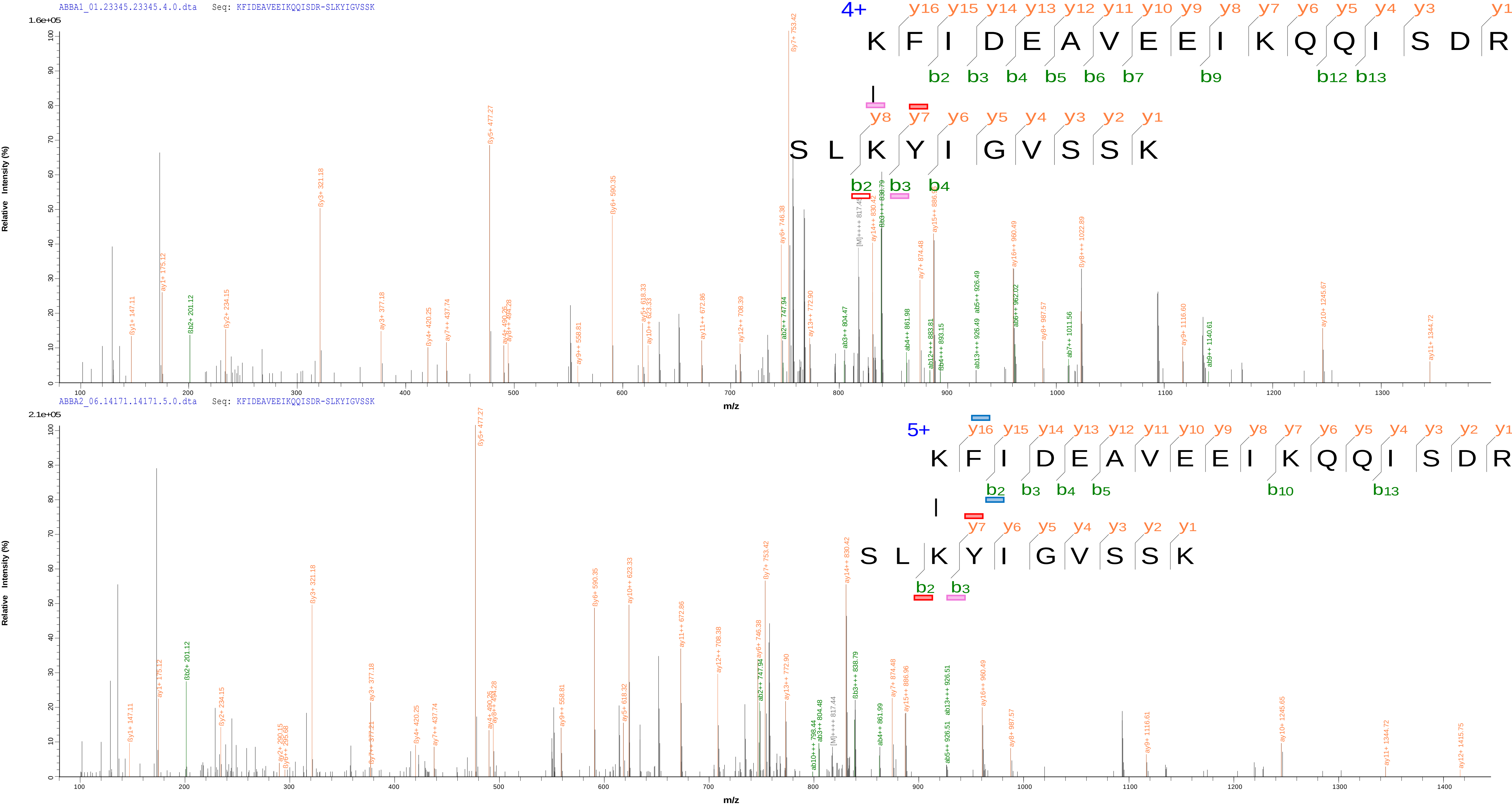

### 2 MjATPPase(1)-MjGATase(20) MFDPKK(1)-SLKYIGVSSK(3)

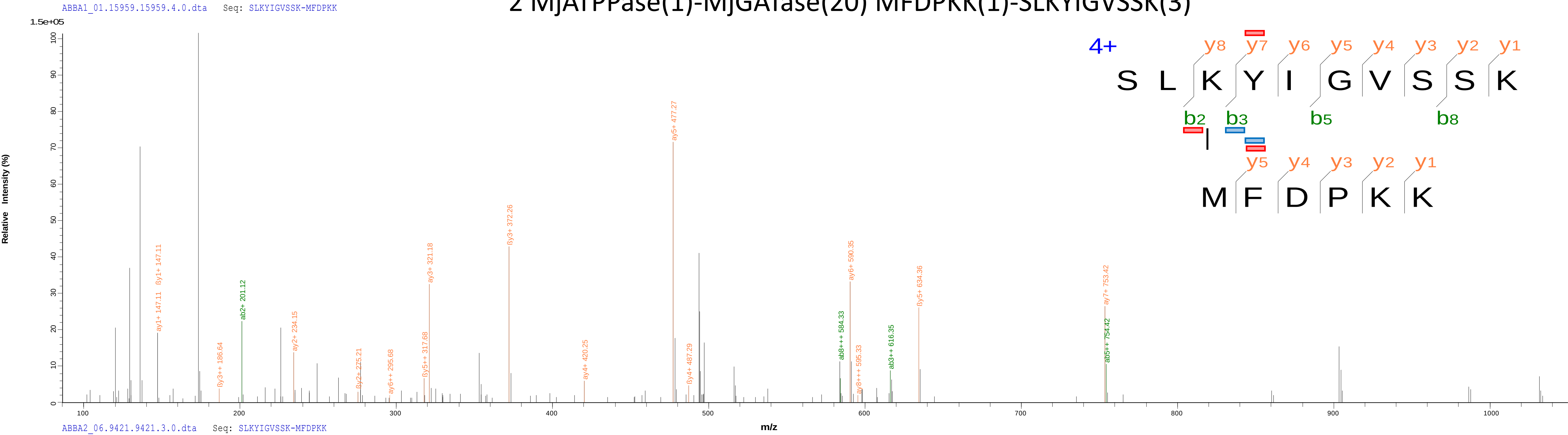


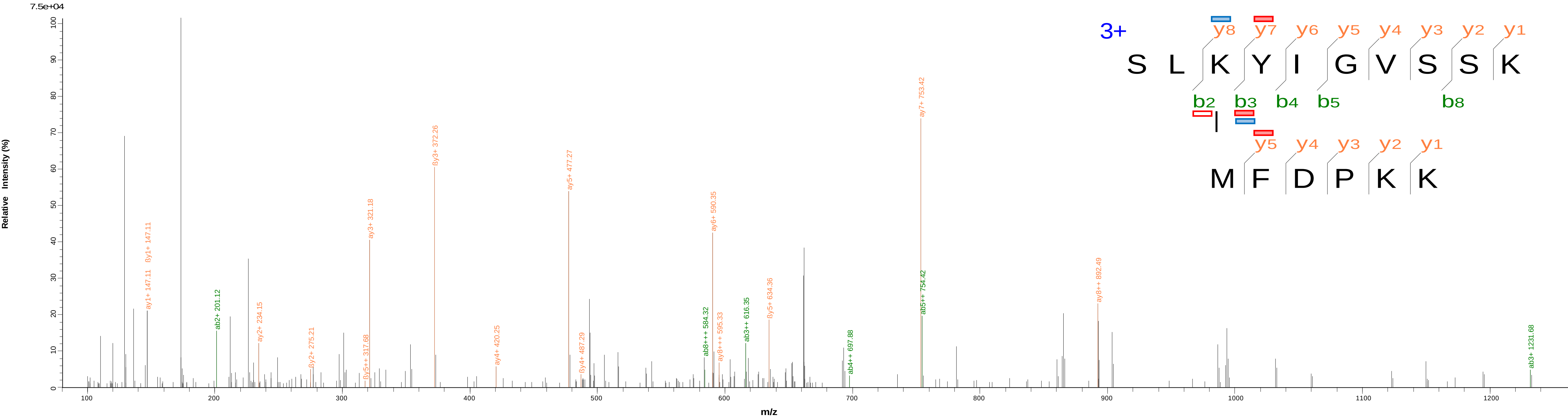

### 3 MjATPPase(174)-MjGATase(20) LLAKELGLPDSIVYR(4)-SLKYIGVSSK(3)

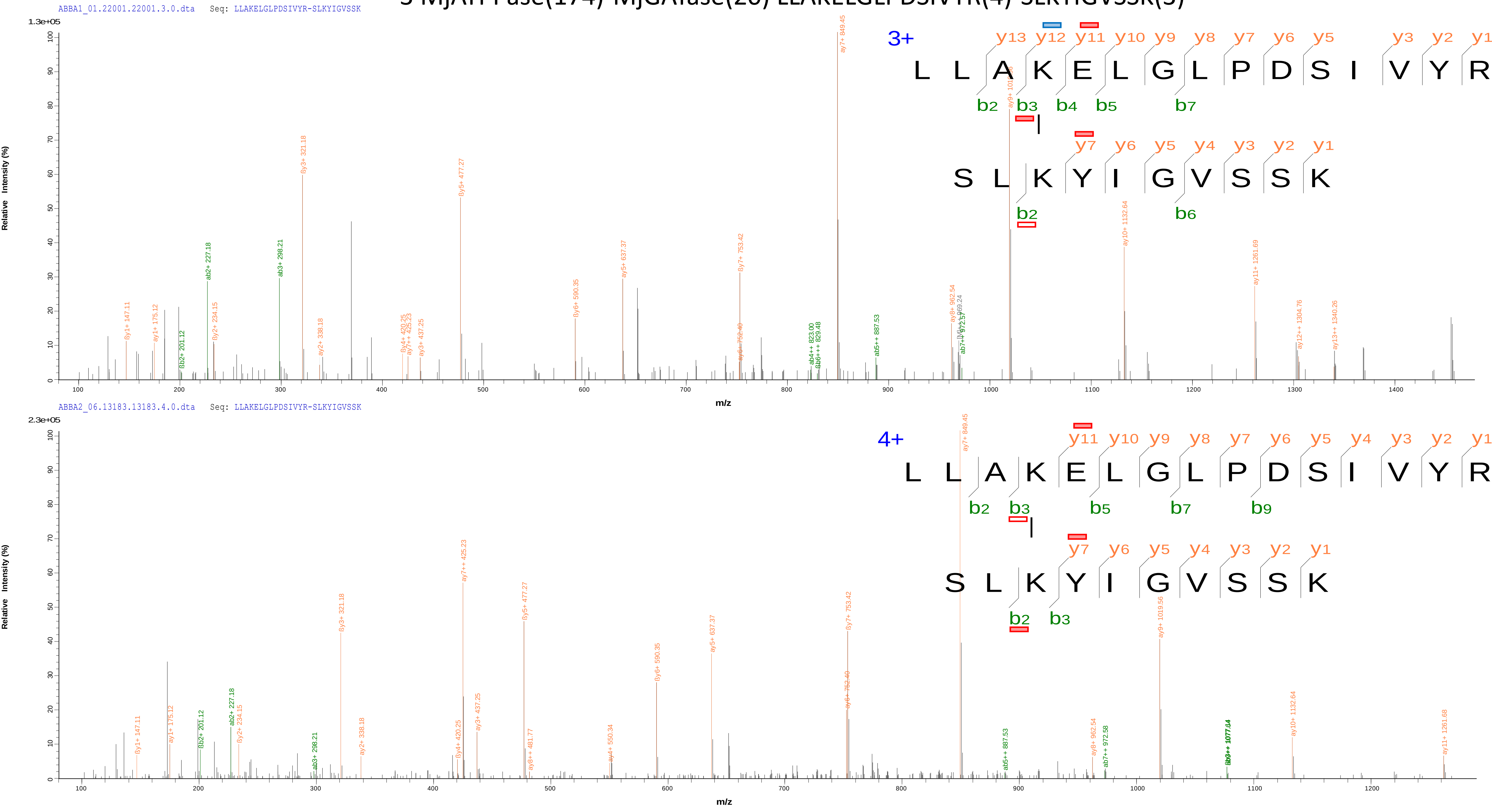

### 4 MjATPPase(16)-MjGATase(20) FIDEAVEEIKQQISDR(10)-SLKYIGVSSK(3)

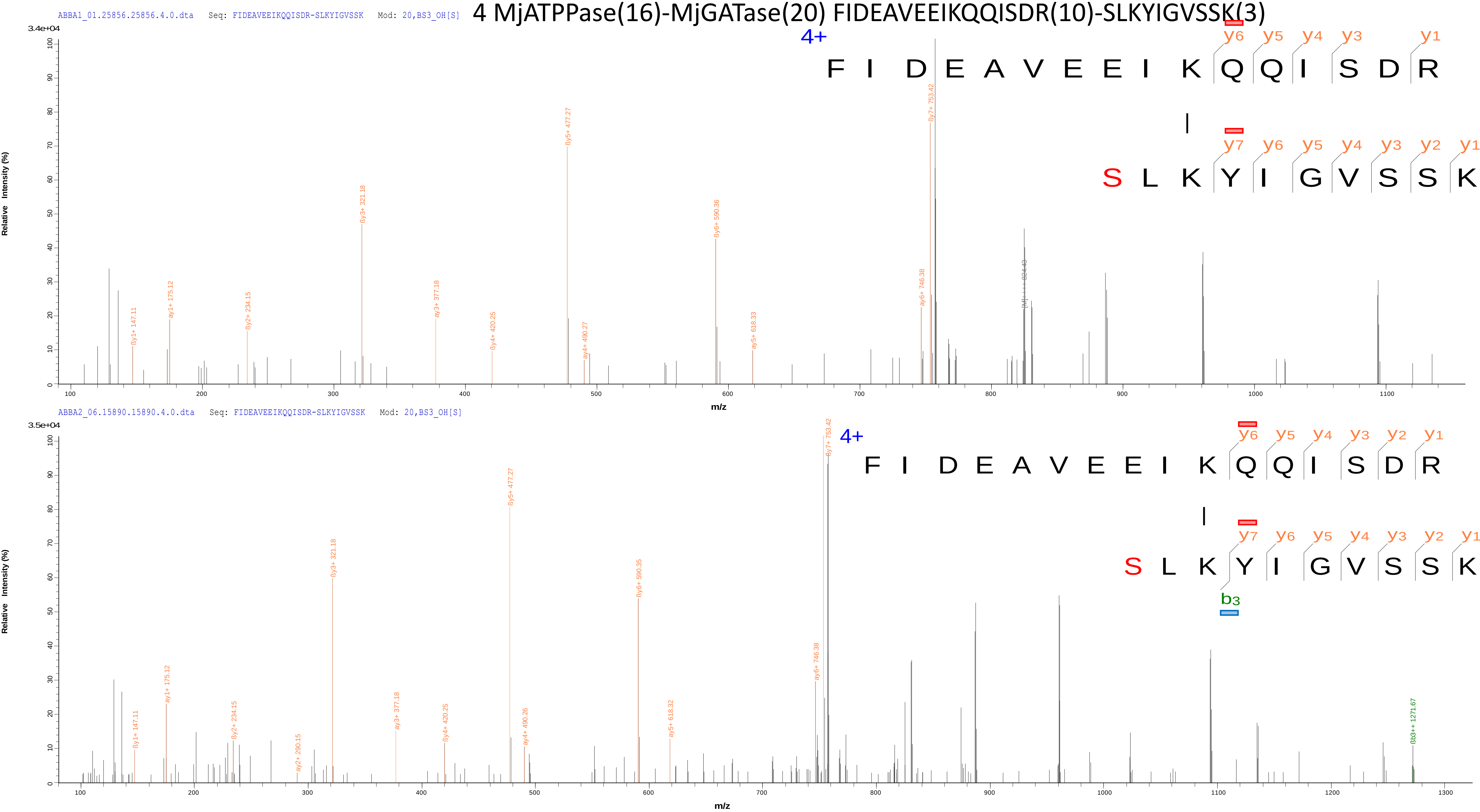

5 MjATPPase(1)-MjGATase(27) MFDPKK(1)-SLKYIGVSSKIVPNTTPLEEIESNKEVK(10)

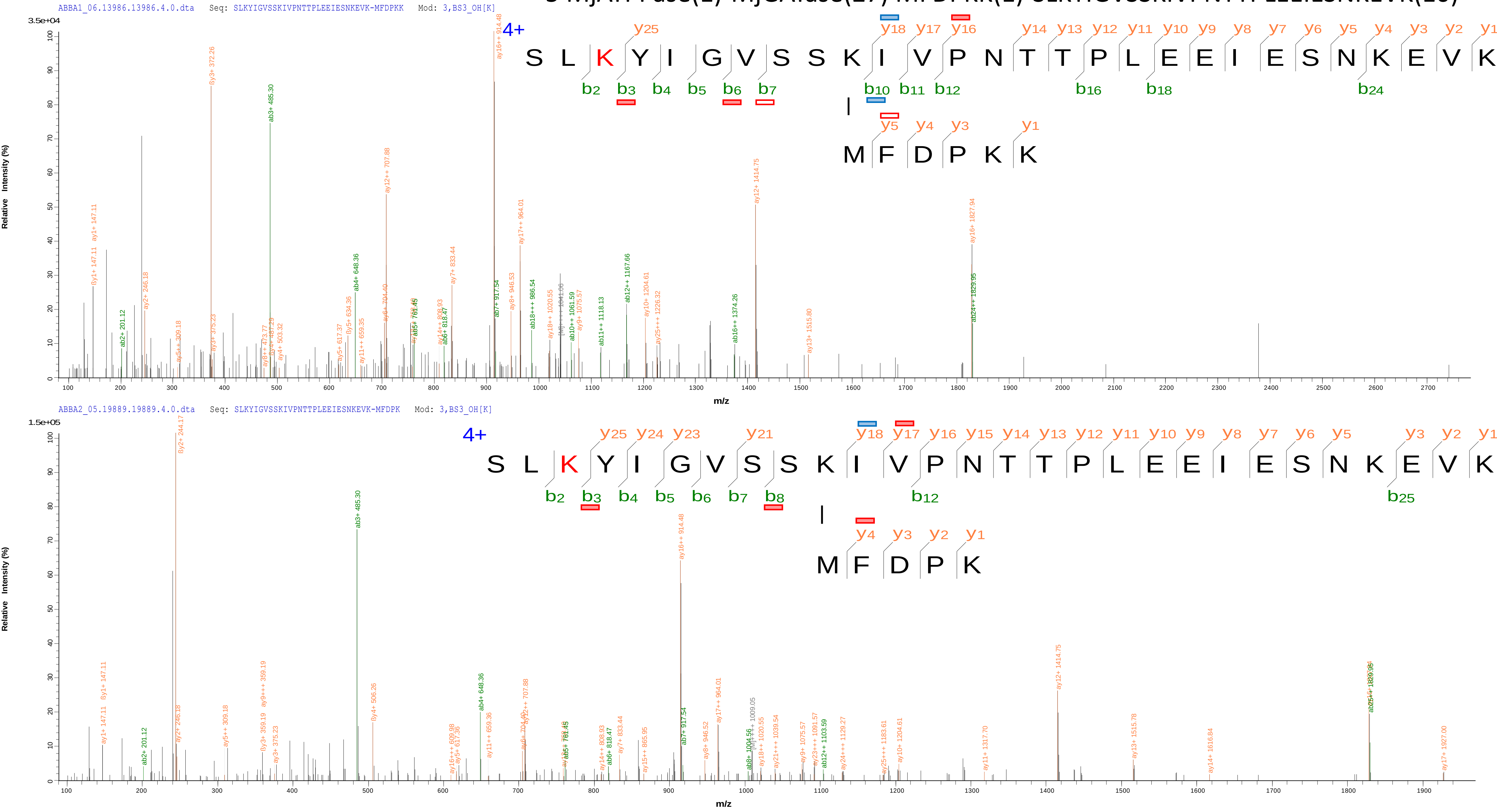

6 MjATPPase(302)-MjGATase(20) VVFDITDKPPATIEFE(8)-SLKYIGVSSK(3)

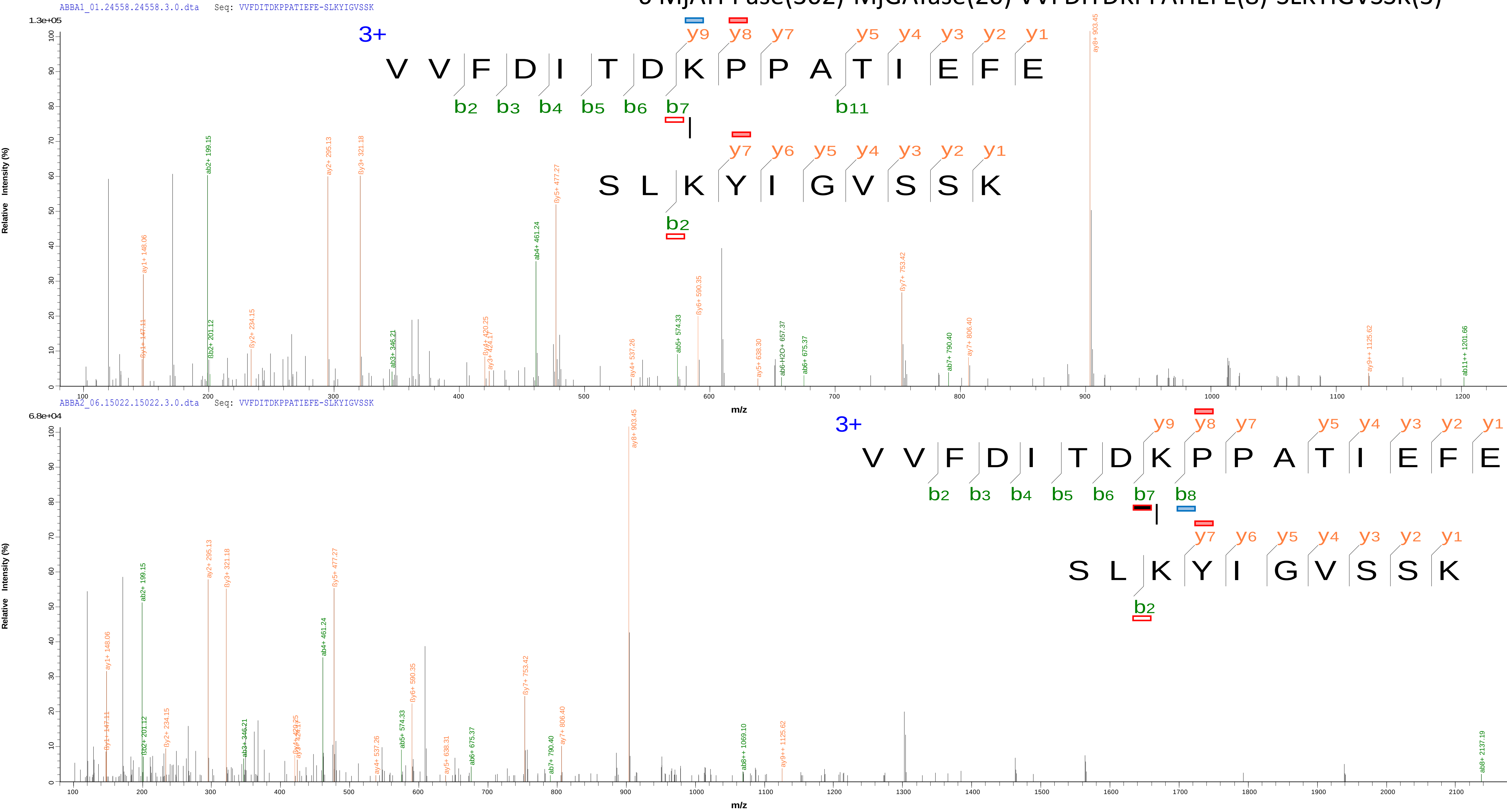

### 7 MjATPPase(1)-MjGATase(1) MFDPKK(1)-MIVILDNGGQYVHR(1)

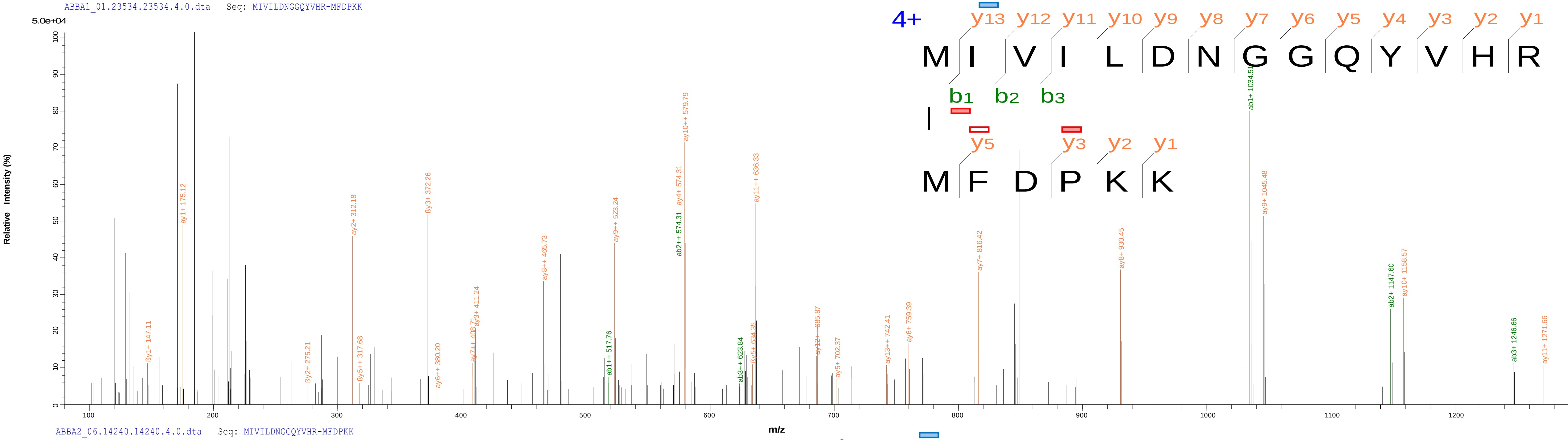


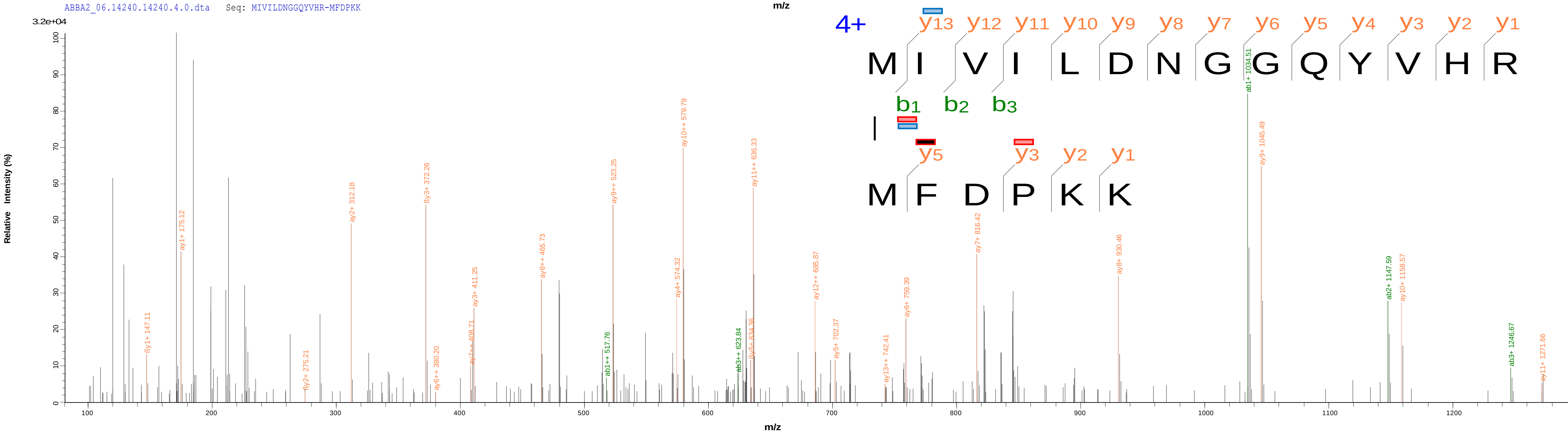

8 MjATPPase(279)-MjGATase(126) SLDAMTAHVPEIPFDLLKR(18)-EFNAWASHKDEVK(9)

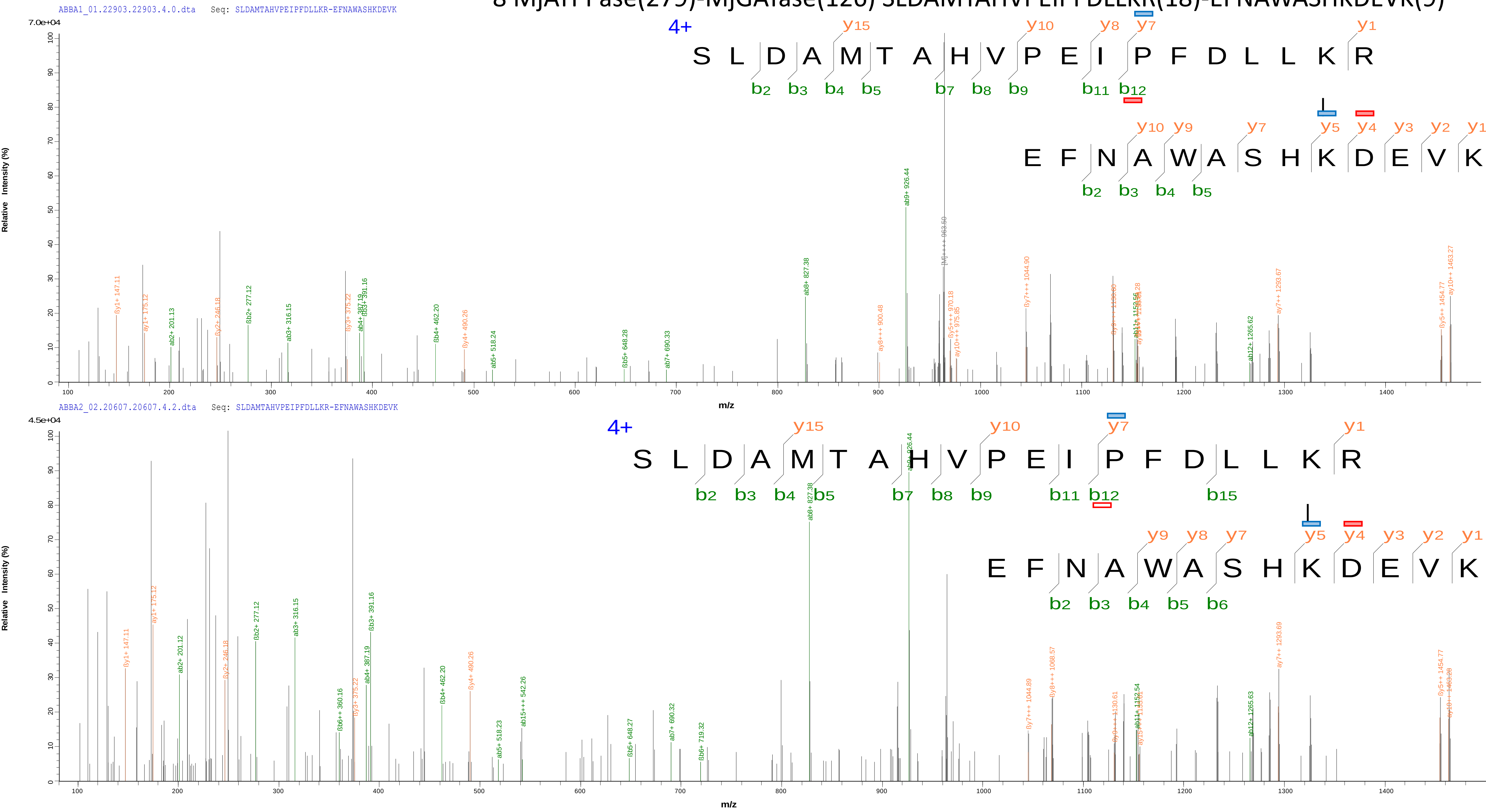

### 9 MjATPPase(1)-MjGATase(177) MFDPKK(1)-TKPIYGVQFHPEVAHTEYGNEILKNFCK(24)

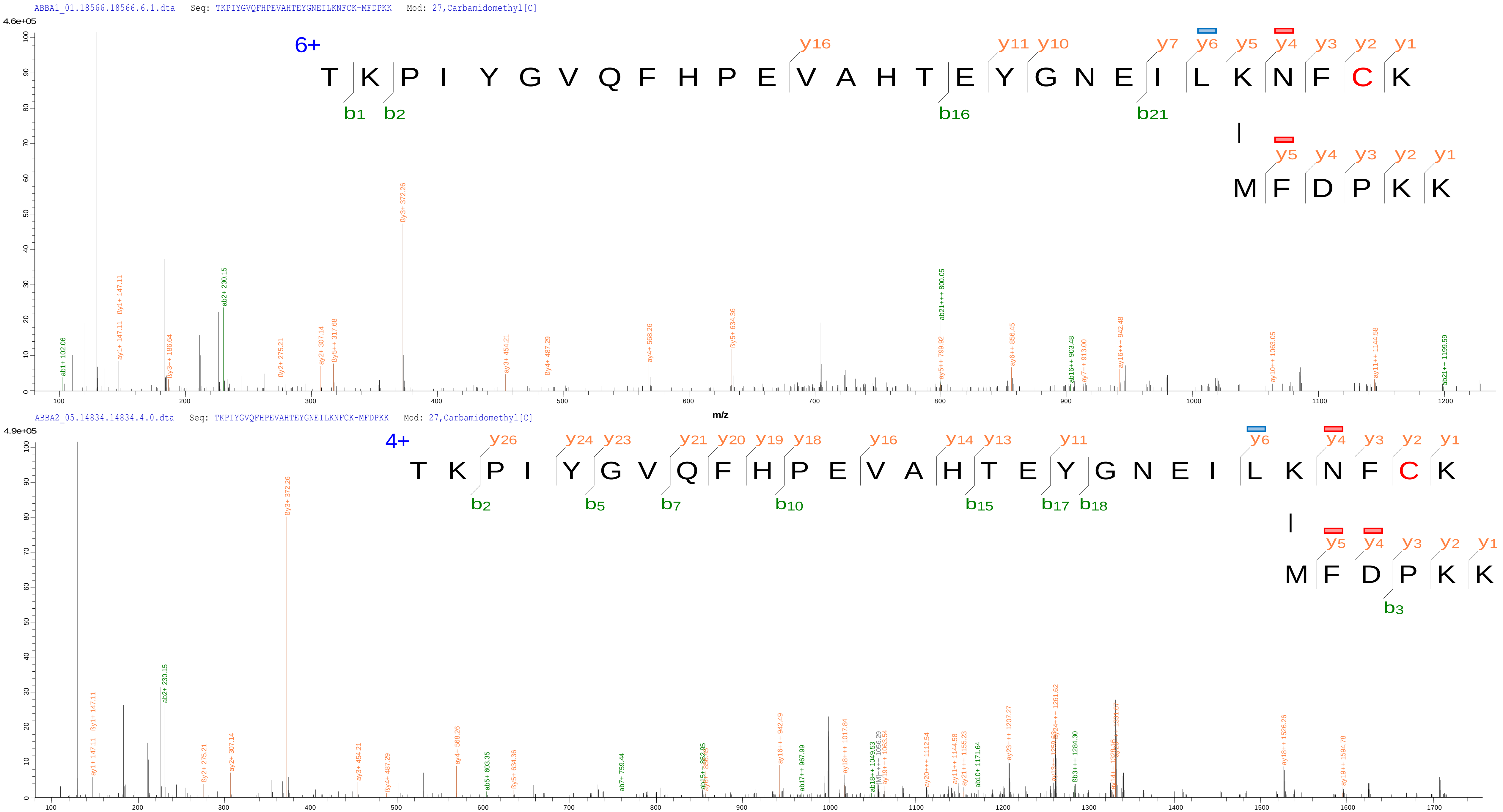

### 10 MjATPPase(5)-MjGATase(20) MFDPKK(5)-SLKYIGVSSK(3)

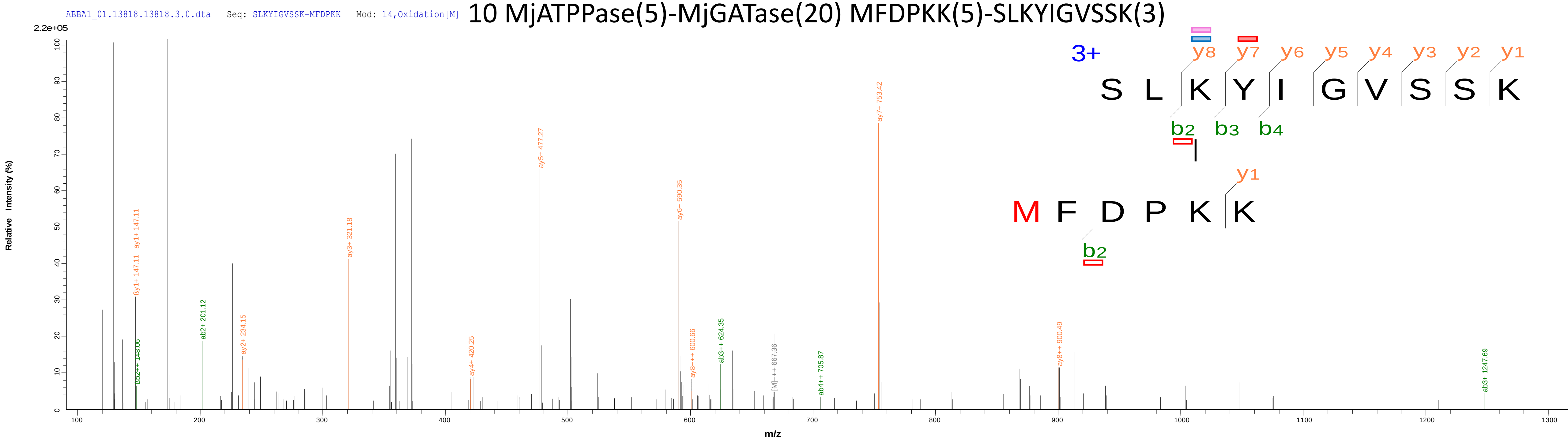


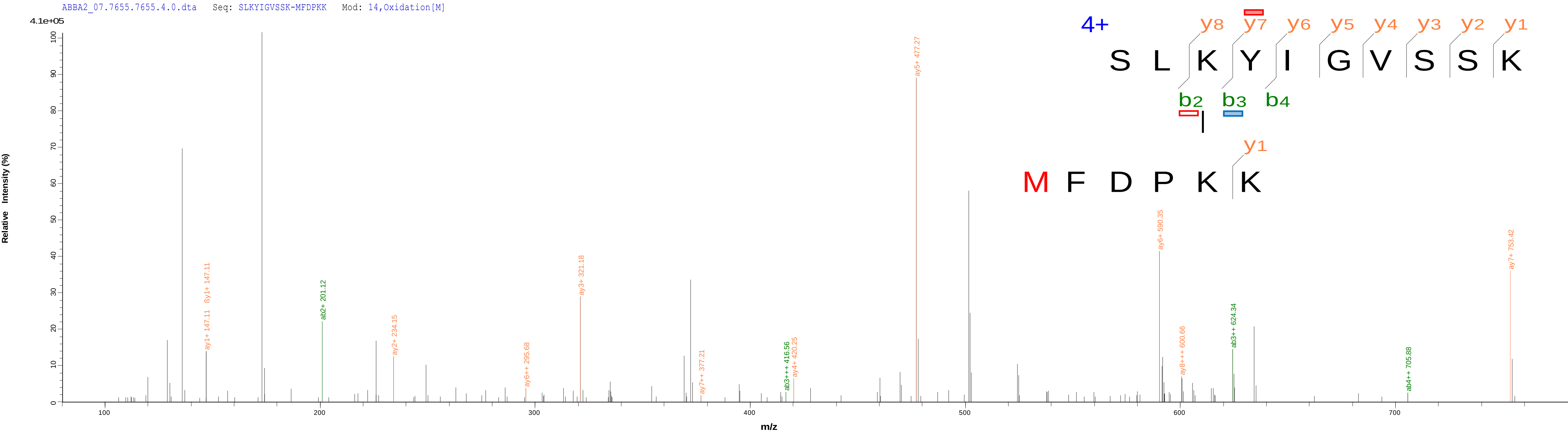

### 11 MjATPPase(222)-MjGATase(1) EANAIVEEEVKKANLDK(12)-MIVILDNGGQYVHR(1)

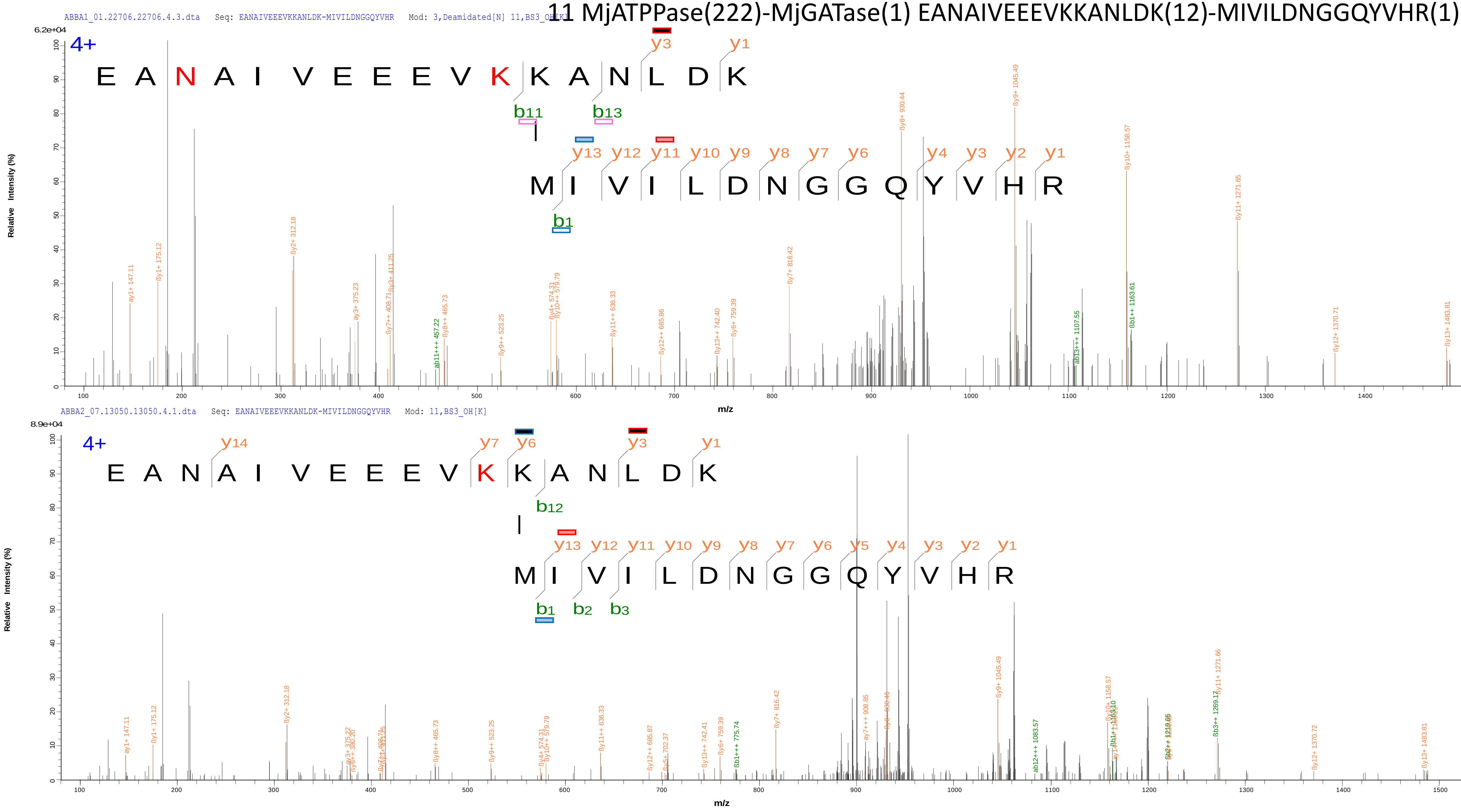

### 12 MjATPPase(1)-MjGATase(186) MFDPKK(1)-VCGYKFE(5)

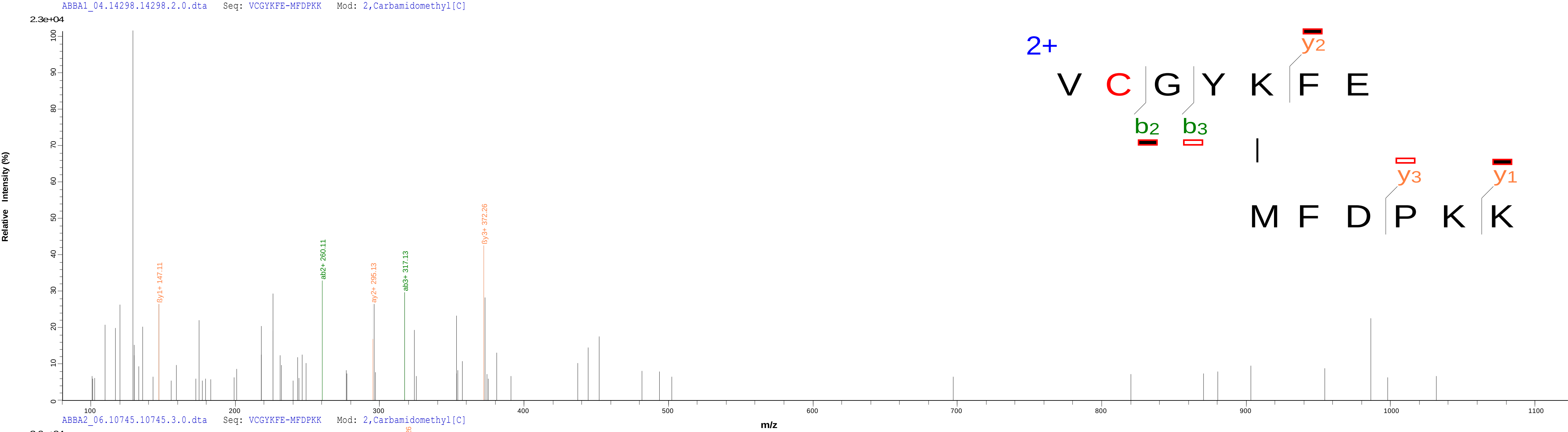


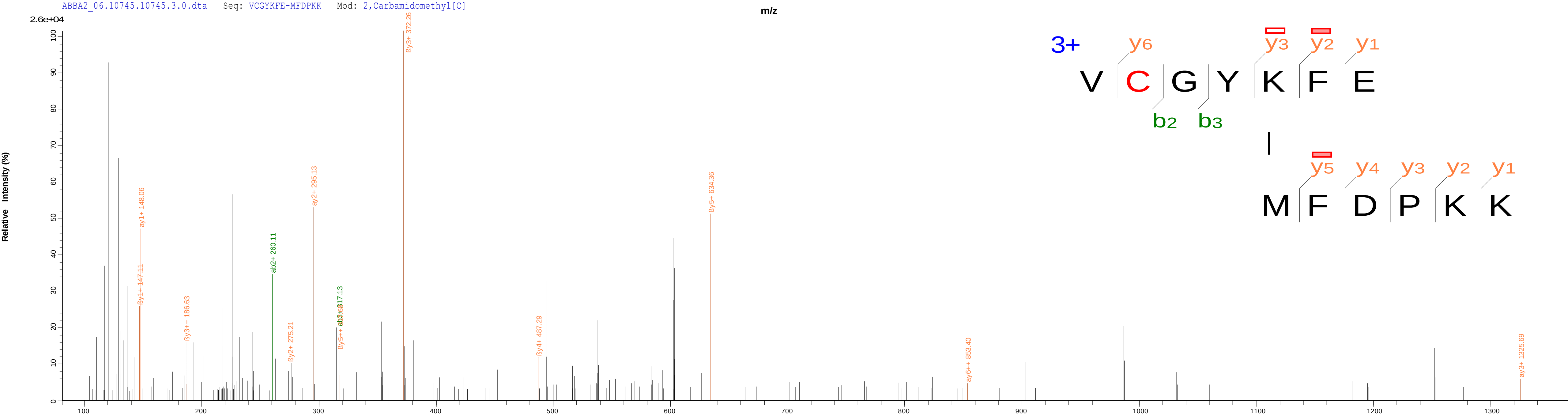

### 13 MjATPPase(93)-MjGATase(155) FLNALKGVTDPEEKR(6)-TKPIYGVQFHPEVAHTEYGNEILK(2)

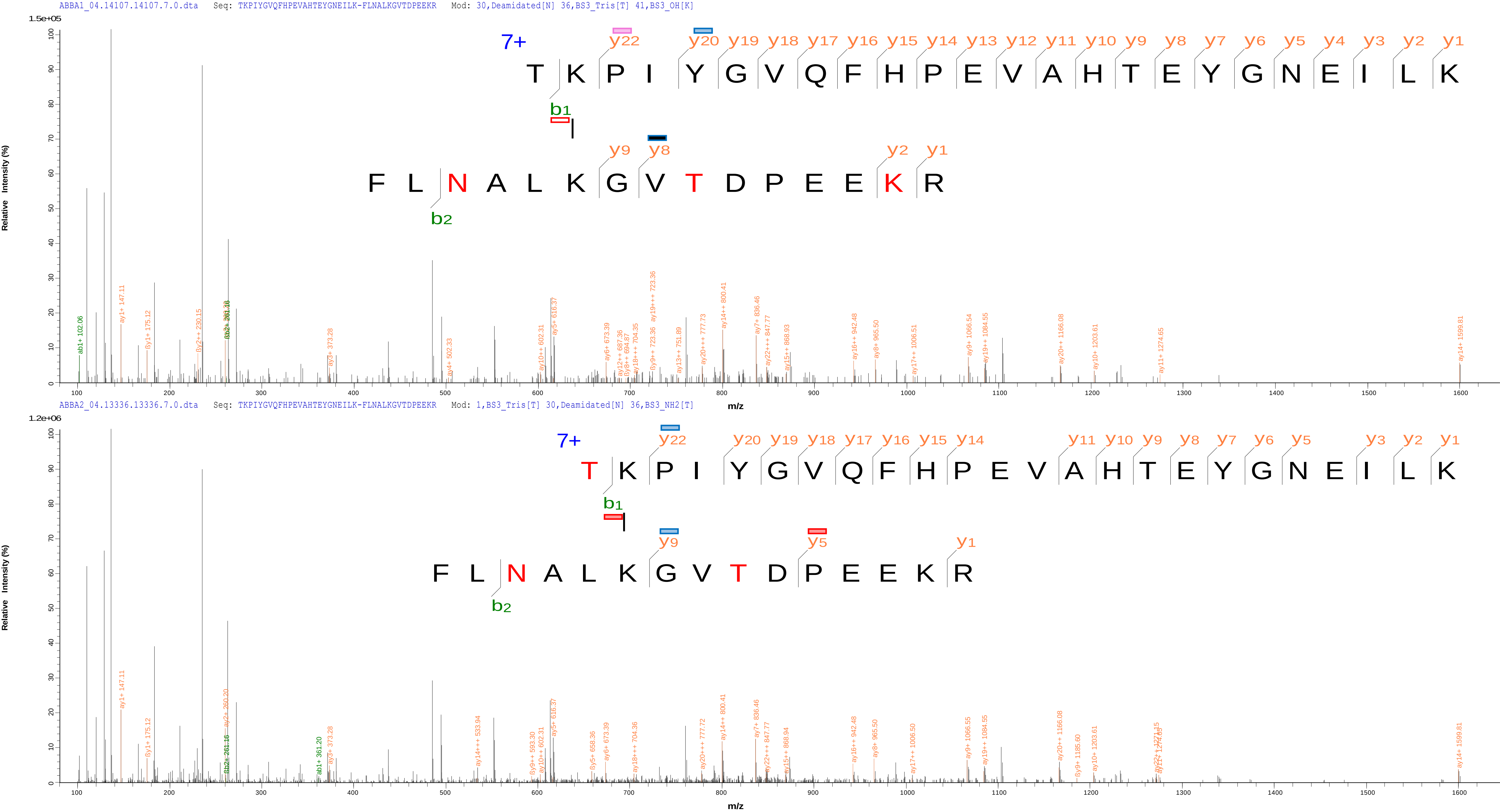
